# Structural stability of mutualistic networks over large geographic and temporal scales

**DOI:** 10.1101/2025.10.08.681159

**Authors:** Benoît Perez-Lamarque, Jérémy Andréoletti, Baptiste Morillon, Orane Pion-Piola, Amaury Lambert, Hélène Morlon

**Affiliations:** Université de Toulouse, Toulouse INP, CNRS, IRD, CRBE, Toulouse, France; Institut de biologie de l’École normale supérieure (IBENS), École normale supérieure, CNRS, INSERM, Université PSL, 46 rue d’Ulm, 75 005 Paris, France

**Keywords:** species interactions, macroevolution, bipartite network, plant mutualism, latent trait, Brownian motion

## Abstract

Mutualistic interactions form species-rich, complex networks that play essential roles for ecosystem function. Over macroevolutionary time scales, global- and continental-level networks change as species emerge and go extinct, yet the stability of their structural organization remains poorly understood. Here, we develop ELEFANT, a novel method for reconstructing ancestral interaction networks based on the hypothesis that species interactions are shaped by unobserved traits that evolve over evolutionary time. We show that ancestral interaction networks can be reliably reconstructed from present-day phylogenetic and interaction data. We infer the ancestral networks of plant mutualisms involving arbuscular mycorrhizal fungi, bat pollinators, and bird seed dispersers at large biogeographic scales. We find that these mutualistic networks exhibit a modular structure that seems to have persisted for millions of years, in part maintained by the evolutionary conservatism of species interactions. As species diversify, they tend to show limited shifts in mutualistic partners, which results in a remarkable long-term stability of mutualistic network structure at large spatial scales.

## Introduction

Mutualistic interactions play a key role in ecosystems (1), and have also been central to the evolution of life on Earth (2, 3). Species involved in mutualistic interactions are interconnected through shared partners, forming complex bipartite interaction networks (4). The structure of these networks is typically non-random, often displaying nestedness – where specialist species preferentially interact with generalists, and/or modularity, in which distinct subsets of species interact more frequently among themselves than with species outside their group (5–7), two patterns that can be present simultaneously (8–10). Historically, research has focused on local, community-scale networks, revealing that non-random structures can influence community stability. More recently, networks compiled at regional to global scales have revealed a tendency toward modularity (11). However, this tendency is not universal (12, 13), and its underlying causes remain poorly understood.

Present-day networks at regional to global scales result from ecological and evolutionary processes operating over millions of years (6, 14). Fundamental yet unresolved questions include when their structural organization emerged, what processes shaped it, and whether these structures are evolutionary stable (15–17), despite the ecological lability of certain interactions (18). If species distributions tend to be restricted to specific biogeographic areas, for example due to dispersal limitation or ecological filtering, we may expect the formation of stable modules aligned with biogeographic regions (11). Likewise, if mutualistic interactions are largely conserved as species diversify, we may expect the formation of stable modules composed of closely-related species (19). Conversely, if a significant fraction of species is ubiquitous and mutualistic interactions are more evolutionarily labile, large-scale networks may be more nested, a structure that enhances robustness to species extinctions (5).

The degree to which species biogeographic distributions are restricted, and mutualistic interactions evolutionary conserved, is therefore expected to have important implications for network structure at large spatial scales. Species distributions depend mostly on the dispersal capacity and niche breath of species. Periods of environmental stability may favor ecological specialization, which may reduce niche breadth and favor modular structures. As for the phylogenetic conservatism in interaction partners, we may expect selection to favor the retention of existing beneficial associations (20). Coevolution may then reinforce reciprocal specialization and compartmentalization (21). However, several processes can counteract this stabilizing selective force, favoring partner shift and reducing phylogenetic conservatism. These include opposing selective pressures during periods of environmental instability, competition for shared partners between closely-related species (22), pairwise coevolution across geographically distinct regions resulting in independent local adaptations (23), and the exploitation of mutualism by cheaters, which can destabilize mutualistic interactions (24). Indeed, empirical evidence for phylogenetic conservatism of mutualistic interactions is mixed (25).

Hypotheses on the emergence and evolutionary stability of network structures are particularly challenging to test. Indeed, direct observation of the fossil record is limited by both low taxonomic resolution and a lack of evidence with respect to the interactions. Only a few networks have been reconstructed from fossil deposits (26, 27), and these primarily represent trophic interactions, which leave more readily interpretable evidence than mutualistic ones. An alternative is to reconstruct ancestral networks (*i.e.* probable ancestral interactions) using knowledge of present-day interactions and phylogenetic relationships. Inspired by models of historical biogeography (28), Braga *et al.* (29) developed a probabilistic model that allows reconstructing ancestral host repertoires (*e.g.* the set of plant families a given butterfly species feeds upon) along a phylogeny (29, 30). However, this approach considers fixed host repertoires (*e.g.* a fixed set of plant families) rather than co-diversifying bipartite interaction networks and is computationally costly, hampering a fine-grained characterization of ancestral network structure, particularly at large biogeographic scales.

Here, we begin by developing an efficient method to reconstruct ancestral networks, based on the idea that interactions are shaped by unobserved traits evolving over the course of history (31, 32). We then apply this approach to four large scale networks of plant mutualisms. For hundreds of millions of years, plants have engaged in diverse and persistent mutualistic interactions with animals and microorganisms, which play crucial roles in their nutrition, protection and reproduction (33–35). We reconstruct ancestral mutualistic networks of plants with arbuscular mycorrhizal fungi at the global scale over the past 150 Myr, bat pollinators in the Neotropics over the past 12 Myr, and bird seed dispersers in Europe and in the Americas over the past 10 Myr, and track how their structure changes through time.

## Results

We developed the “Evolution of LatEnt traits For Ancestral NeTwork reconstruction” model (ELEFANT, Fig. 1, see Methods). We assume that interspecific interactions are shaped by species-specific traits, such as the production and detection of floral scents, or the morphological complementarity of spur and proboscis lengths in pollination systems. However, such traits are often unknown and/or difficult to measure in most empirical systems. Thus, instead of relying on directly measured traits, ELEFANT infers unobserved “latent” traits from the present-day interaction matrix, using singular value decomposition (31, 32) (see Methods ; Fig. 1.1). The methods directly follow (31, 32), although with the goal to infer past interactions rather than unobserved, present-day ones, as in (31, 32, 36). By construction, each species is represented by a vector of several latent trait values, and an interaction between a pair of species is inferred when the product of their corresponding latent trait values exceeds a network-specific threshold *J*, chosen to maximize fit to the observed interaction network. This interaction rule does not match classical formulations of trait matching (as, for example in (6, 37)), yet as we will show, it is effective in representing who interacts with whom. Next, we “augment” the phylogenetic trees associated with the present-day interaction network in order to account for lineages that are, or were, likely involved in the system considered (e.g., bird seed dispersers in Europe over the past 10 Myr) but are not represented in the trees, because they went extinct or were not sampled (Fig. 1.2). When diversification occurs *in situ* at the spatial scale of the network, such as in large (continental to global) networks over relatively short timescales, or in close communities shaped by local radiations (e.g., on remote islands or lakes), phylogenetic trees can be modeled by assuming that clades evolve according to a birth-death process representing speciation and extinction events (38). We use data augmentation under this process to obtain “complete” trees with probable configurations of missing lineages (39). The latter step is specific to ELEFANT: reconstructing lineages that were likely involved in ancestral interactions but are missing in reconstructed trees is not necessary when inferring unobserved, present-day interactions (31, 32, 36), but ignoring them would result in a massive undersampling of ancestral networks. Finally, as in (32), we assume that the latent traits evolve according to traditional phylogenetic models of trait evolution, such as the Brownian motion process. We use ancestral trait reconstruction to infer latent trait values along the augmented phylogenies (32, 40)(Fig. 1.3), and deduce ancestral networks (Fig. 1.4). This “ancestral” trait reconstruction infers latent traits on all branches of the augmented phylogenies, including at present for species which interacting partners are not known. An additional specificity of ELEFANT is to use stochastic character mapping to propagate uncertainties in the character reconstruction on the inferred ancestral networks. Like any model, ELEFANT relies on strong assumptions, and *at a minimum,* requires interactions to be evolutionary conserved to some extent. This condition may not always be met; therefore, the final step of our procedure is specifically designed to assess whether the model can be reliably applied to the network under investigation (Fig. 1.5 and S1). The principle of this “cross-validation” procedure is to assess the ability of the model to recover known interactions. Specifically, we artificially remove (subsample) a given percentage of present-day species, estimate the latent traits using the network formed by the remaining species, and then reconstruct these traits along the complete phylogeny, including at present for the subsampled species. From these reconstructions, we infer the interactions of the subsampled species. Comparing the inferred interactions with the known (“true”) ones yields a F1-score. We refer to this F1-score, computed at present, as the “cross-validation score”. Scores close to 0 indicate that the data violate one or more model assumptions (*e.g.* Brownian evolution of latent traits) and, consequently, that the model should not be used to infer ancestral networks. By diagnosing the predictive performance of ELEFANT using present-day data, the cross-validation score provides insight into its ability to infer ancestral interactions under its underlying assumptions.

**Figure 1:**
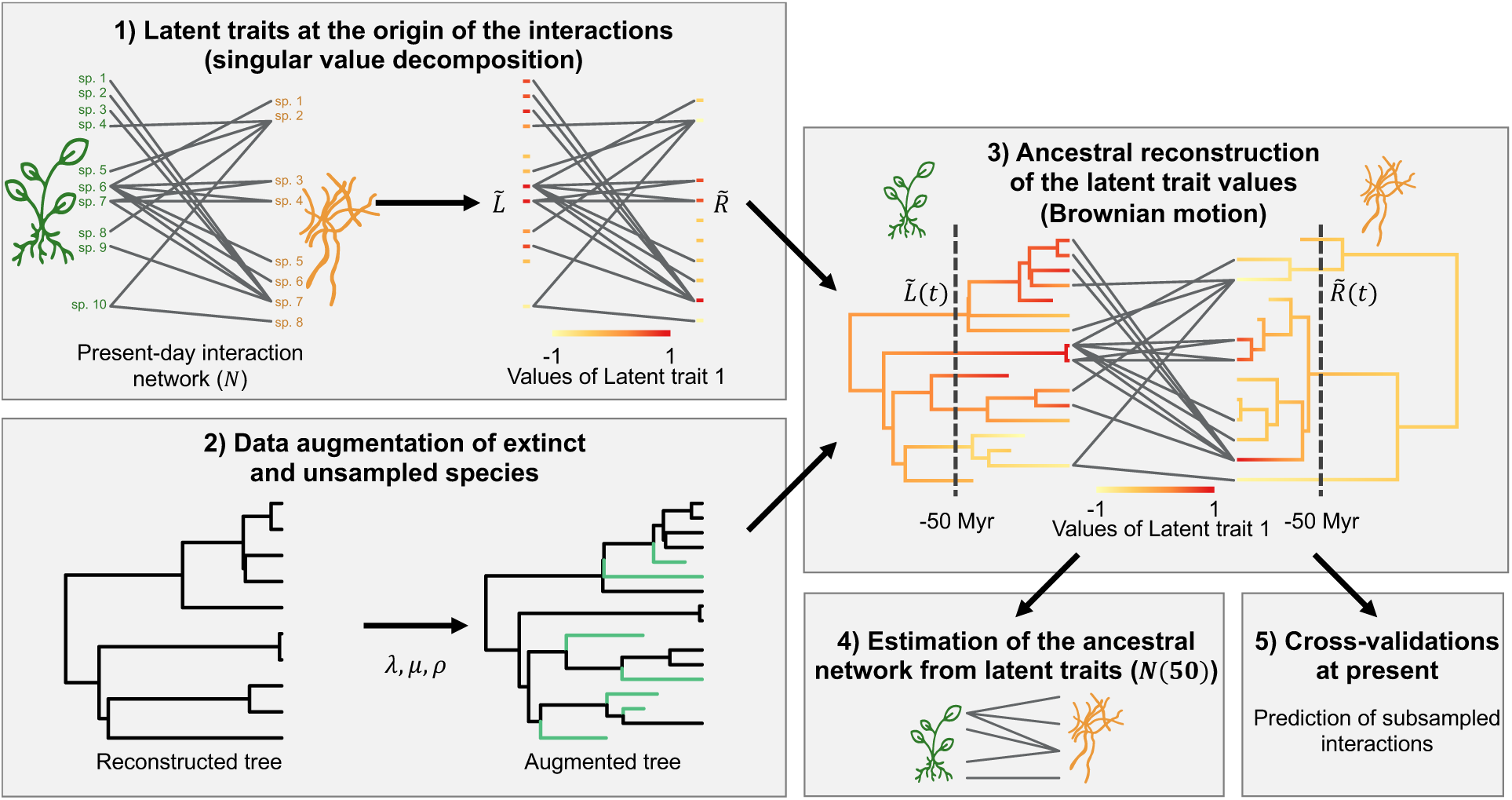
General framework used to reconstruct ancestral networks. The Evolution of LatEnt traits for Ancestral Network reconsTruction framework (ELEFANT), proceeds in five steps. 1) Given a bipartite interaction network (*N*), a vector of *d* latent trait values is estimated for each extant species on each side of the network; these vectors (compiled into matrices *L̃* and *R̃*) underlie present-day interactions (for clarity only the first latent trait value is represented here with the color code). 2) Data augmentation is used to add extinct and unsampled lineages (colored in green) on the phylogenetic trees, assuming constant rates of speciation (λ), extinction (μ), and sampling at present (ρ). 3) Ancestral values of the latent traits are estimated on the two phylogenetic trees using Brownian motion. 4) Ancestral interaction networks are reconstructed from the estimated ancestral latent trait values at any time in the past (e.g. here, 50 Mya) 5) A cross-validation procedure based on the ability to recover known interactions at present in a specific system provides a measure of confidence.

We thoroughly tested the ability of ELEFANT to reconstruct ancestral interaction networks using simulated data. We mimicked 30 Myr of network evolution during the diversification of two interacting guilds (hereafter referred to as “plants” and “mutualists” for simplicity, but ELEFANT can be applied to any type of bipartite network, including antagonistic ones) under five distinct scenarios (see Methods): i) The first scenario (“evolution by anagenetic shifts”) considers that interactions are inherited at speciation, but can change during a lineage’s lifespan. ii) The second scenario (“evolution by cladogenetic shifts”) adds the occurrence of cladogenetic shifts, where upon speciation of a mutualist species, one of the daughter lineages retains its plant partners, while the other forms an entirely new set of interactions with randomly selected plant species. iii) The third scenario (“Brownian evolution of latent traits”) corresponds to the model underlying ELEFANT, where traits evolve as Brownian processes on the plant and mutualist phylogenies and their products determine whether species interact. iv) In the fourth scenario (“host repertoire evolution”), inspired from Braga *et al.* (29), host repertoires (*i.e.* the set of plant lineages with which a mutualist interacts) are assumed to be fixed while mutualists diversify, as it may be the case when hosts are defined at higher taxonomic levels; host repertoires are inherited upon the speciation of mutualists, and they can be acquired or lost during the lifespan of a mutualist. v) Finally, the fifth scenario (“random evolution”) mimics very frequent anagenetic shifts. In scenarios i), iii) and iv) interspecific interactions tend to be conserved over macroevolutionary timescales, while in scenarios ii) and v) they are more labile. We applied ELEFANT to these simulated data and computed cross-validation scores. In addition, to assess the robustness of our reconstructions, we computed F1-scores from ancestral networks.

We found that our cross-validation procedure effectively discriminates between cases in which ancestral networks can be robustly reconstructed—at least in the recent past (scenarios i, iii, and iv, characterized by conserved interactions)—and cases in which they cannot (scenarios ii and v, characterized by evolutionary labile interactions) (Fig. S2 & S3). Cross-validation scores are above 0.75 for scenario i), and above 0.5 for scenarios iii) and iv), but around 0.20 for scenario ii) and below 0.10 for scenario v) (Fig. S2). The ability to recover correct ancestral interactions (F1-scores from ancestral networks) is well correlated with the cross-validation scores at present (Fig. S3). A cross-validation score above 0.5 indicates that the inferred ancestral interactions are highly accurate (F1-score >0.75), and a score between 0.25 and 0.5 typically indicates that the inferred ancestral interactions are likely to be correct in the recent past (F1-score >0.5; Fig. S3). These results suggest that a cross-validation score of 0.25 provides a good operational minimum threshold for the application of ELEFANT to a given empirical system. When interactions are not too labile, which in practice is reflected by a cross-validation score above 0.25, ancestral interactions are well recovered, in particular in the recent past (Figs. 2 and S4). It follows that global metrics (connectance – *i.e.* the ratio of realized interactions –, nestedness, and modularity) and phylogenetic signal in species interactions are also well recovered, although with a tendency toward homogenization (*i.e.* the lowest and highest simulated metrics tend to be estimated closer to the average; Figs. 2, S5 & S6). The estimations of global metrics are particularly precise close to the present (5 Myr ago: mean R^2^=0.79) and become noisier back in the past (20 Myr ago: mean R^2^=0.32), with a good recovery of temporal trends (see Supplementary results; Fig. S7). These results suggest to be careful with the interpretation of ancestral networks constructed past halfway through the clades’ age. In all cases, ELEFANT did not produce artefactual detection of structural patterns (e.g., nestedness, modularity, phylogenetic signal) when these patterns were not present in the simulations (Fig. S7).

**Figure 2:**
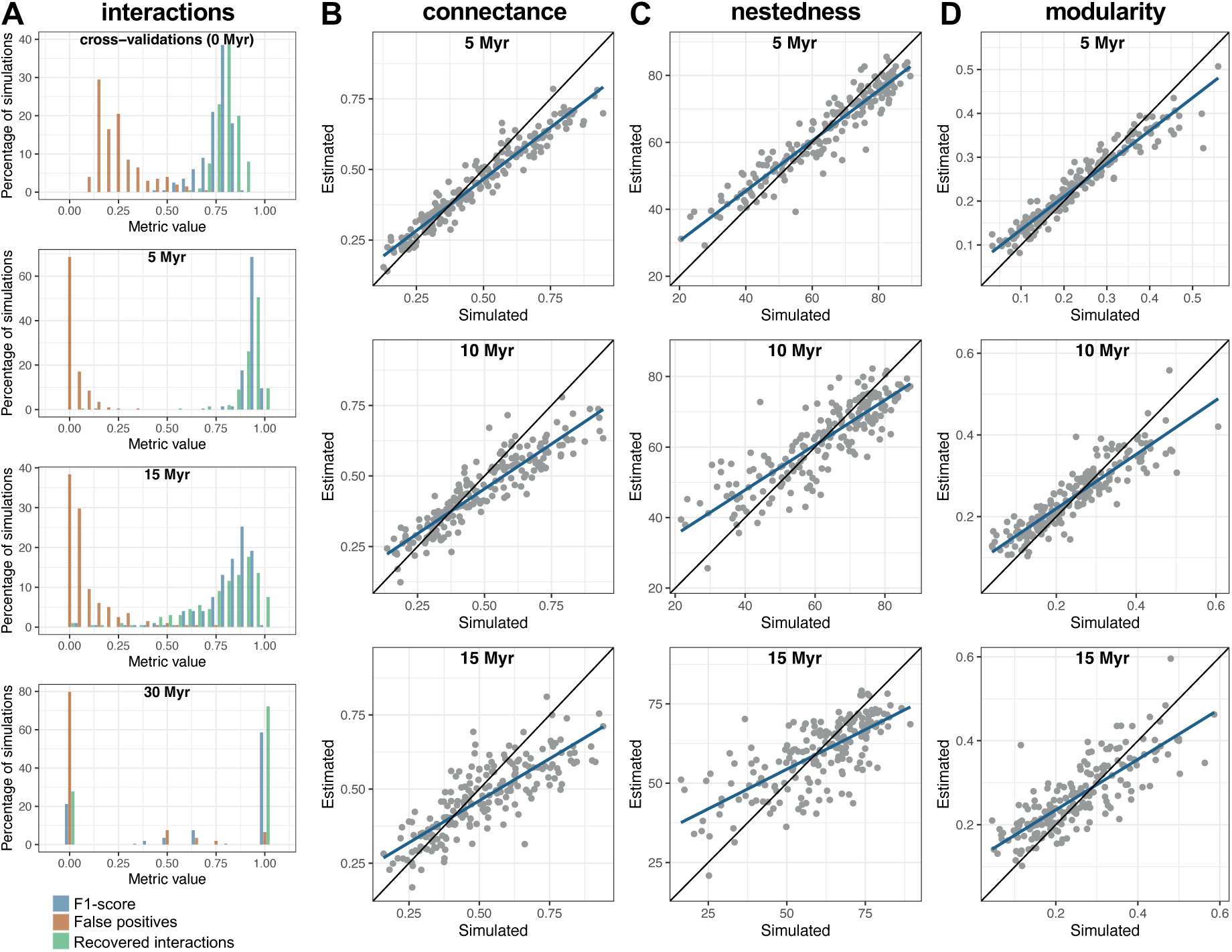
Accurate reconstruction of ancestral networks. **A:** Bar charts indicate the F1-score (blue), the proportion of false positives (red), and the proportion of recovered interactions (green) inferred in the cross-validations at present (upper row) and in the consensus ancestral networks (from 5 to 30 Myr ago; lower rows). 30 Mya corresponds to the most recent common ancestor of the mutualists, which interacted with a single plant species. These consensus networks assemble interactions that are recovered in at least *J*% of the ‘augmented ancestral networks’ where *J* is the classification threshold maximizing the Youden’s *J* statistic. **B-D:** Estimates of the ancestral connectance (B), nestedness (C), and modularity (D) are represented at different times in the past (from 5 to 15 Myr ago) as a function of the simulated values. Each dot corresponds to one simulation, and for each simulation, we reported the estimated values averaged over 250 ‘augmented’ ancestral networks. The black lines represent the line y=x, while the blue lines correspond to the fit of a linear model between the estimated and simulated values. The figure reports results obtained for simulations under the scenario of evolution by anagenetic shifts (see Fig. S4 for other types of simulations).

Confident in our framework, we inferred the ancestral networks for four large plant-mutualist systems (see Methods): i) the global-scale mutualism with arbuscular mycorrhizal fungi (115 plant families and 351 fungal taxa over the past 150 Myr), ii) interactions with bat pollinators in the Neotropics (42 plant families and 56 bat species over the past 12 Myr), iii) bird-mediated seed dispersal in Europe (28 plant families and 70 bird species over the past 10 Myr), and iv) bird-mediated seed dispersal in the Americas (115 plant families and 586 bird species over the past 10 Myr). Cross-validation scores were all above 0.25 (0.32, 0.27, 0.70, and 0.46 for i), ii), iii) and iv) respectively). Incorrect inferences were related to false positives more than to false negatives (Table S1), and at least part of these “false positives” could in fact correspond to true but undocumented interactions (36). Regardless of the reasons for these relatively modest scores (such as undocumented interactions and ecological lability in the interactions), they fall within the range where inferred network metrics are reliable, at least in the recent past. Our reconstructions reveal consistently significant modularity across the four systems, despite variation in species richness and composition over time (Figs. 3, 4 & 5). We did not detect nestedness patterns deviating consistently from null expectations. Plant median degrees (*i.e.,* the median number of partners per species) tend to be lower than expected under null models in three of the four systems (significantly so in two), reflecting a tendency toward specialization. Species interactions appear to be generally evolutionarily conserved to some extant (significant although low phylogenetic signal, except in the smaller size European bird dispersal network). These results are robust to diversification rates and sampling fractions used to augment the trees (Fig. S8). They suggest a scenario where mutualistic associations are partially evolutionary conserved, favoring specialization and contributing to the formation of relatively stable modules, where closely-related taxa tend to interact with similar partners.

**Figure 3:**
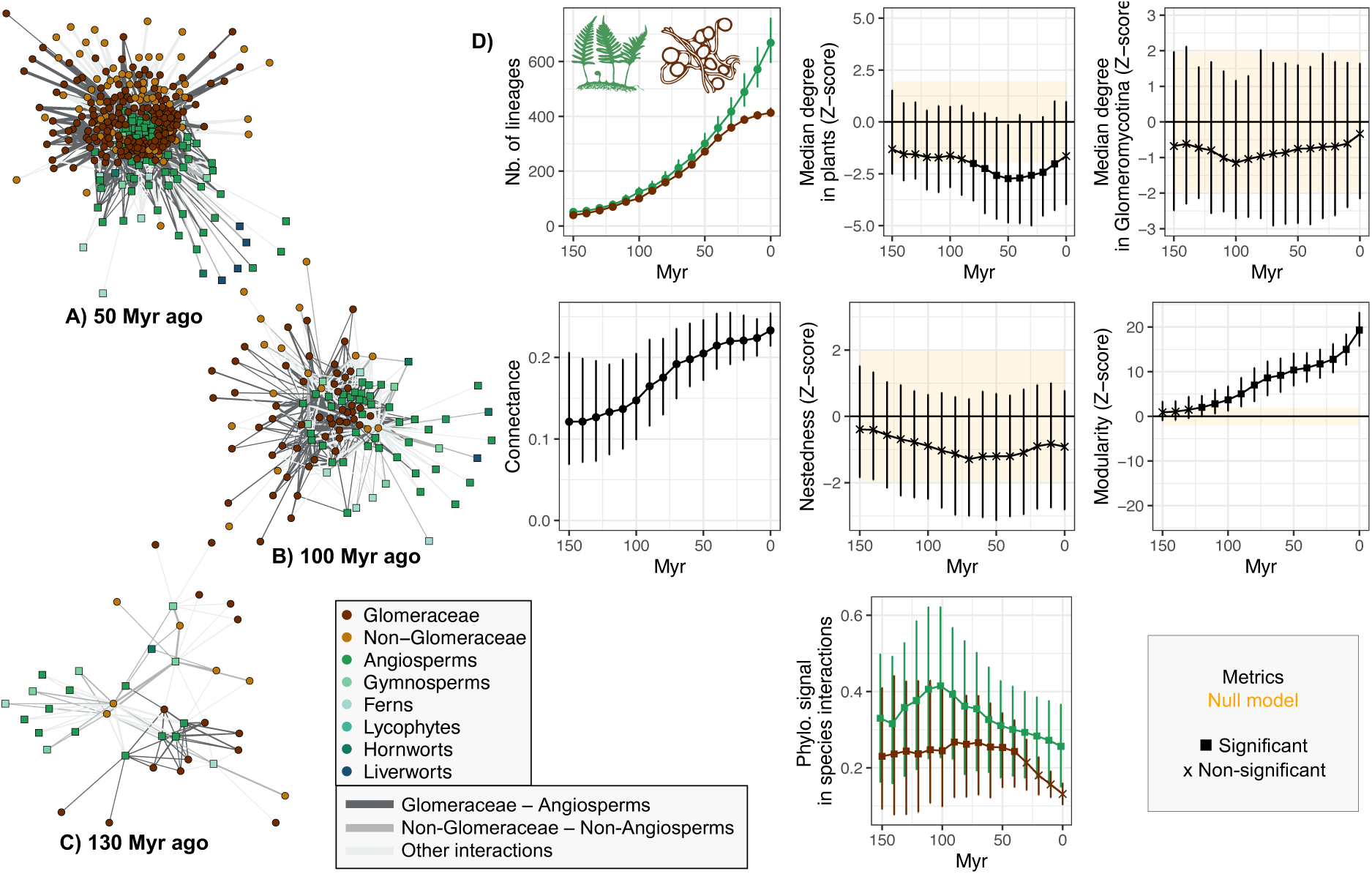
Global scale ancestral network reconstruction of arbuscular mycorrhizal symbiosis. Consensus ancestral networks 50 (A), 100 (B), or 130 (C) Myr ago. Ancestral network metrics (D). Squares indicate a significant departure from the null model (95% central range shown in orange), whereas the crosses indicate non-significance. Error bars represent the 95% central range of values obtained from the set of augmented ancestral networks.

**Figure 4:**
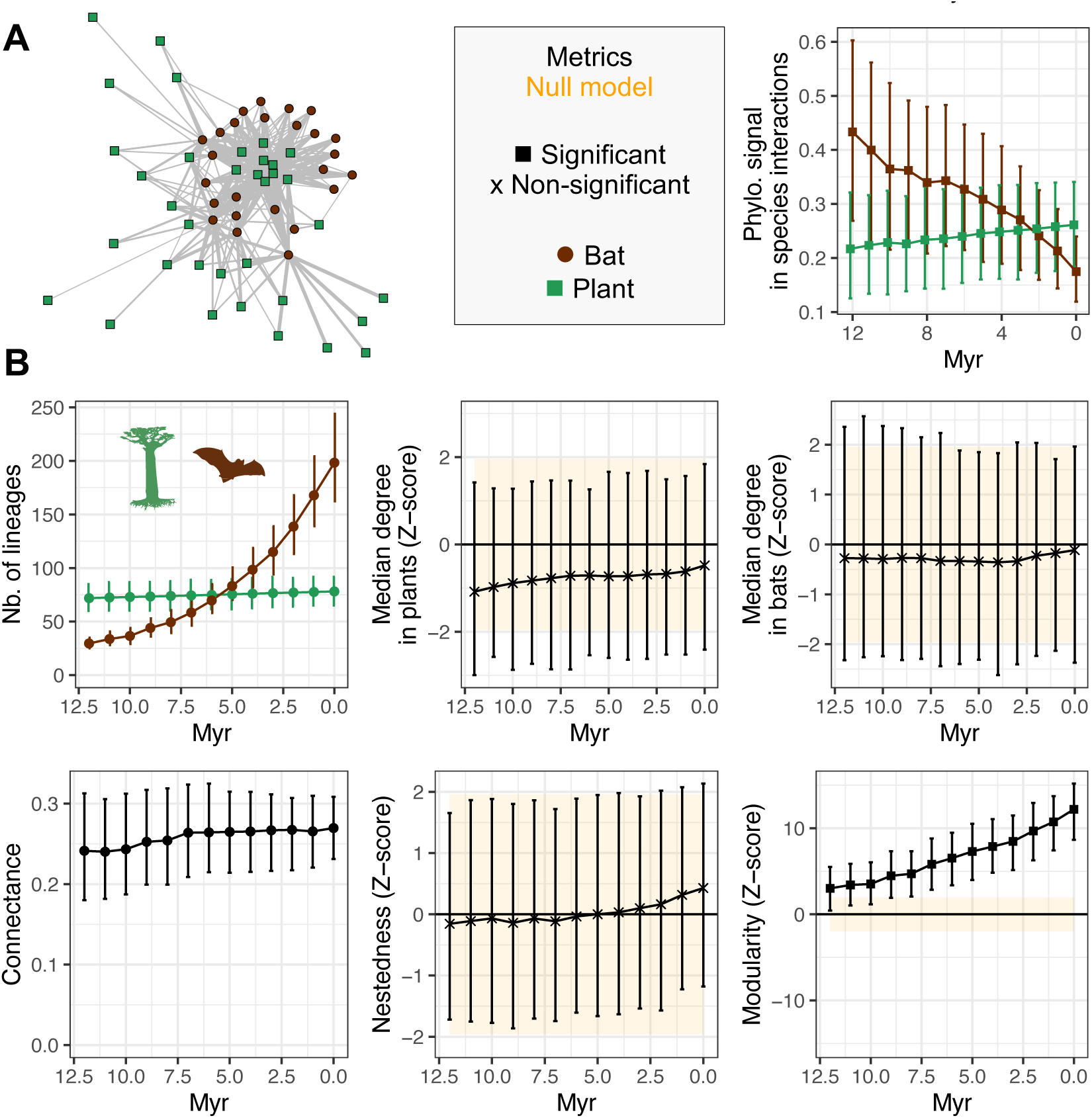
Ancestral network reconstruction of bat pollination in the Neotropics. Consensus ancestral network inferred 10 Myr ago (A). Ancestral network metrics (B). Squares indicate significant, and crosses non-significant departure from the null model (shown in orange), as in Fig. 3.

**Figure 5:**
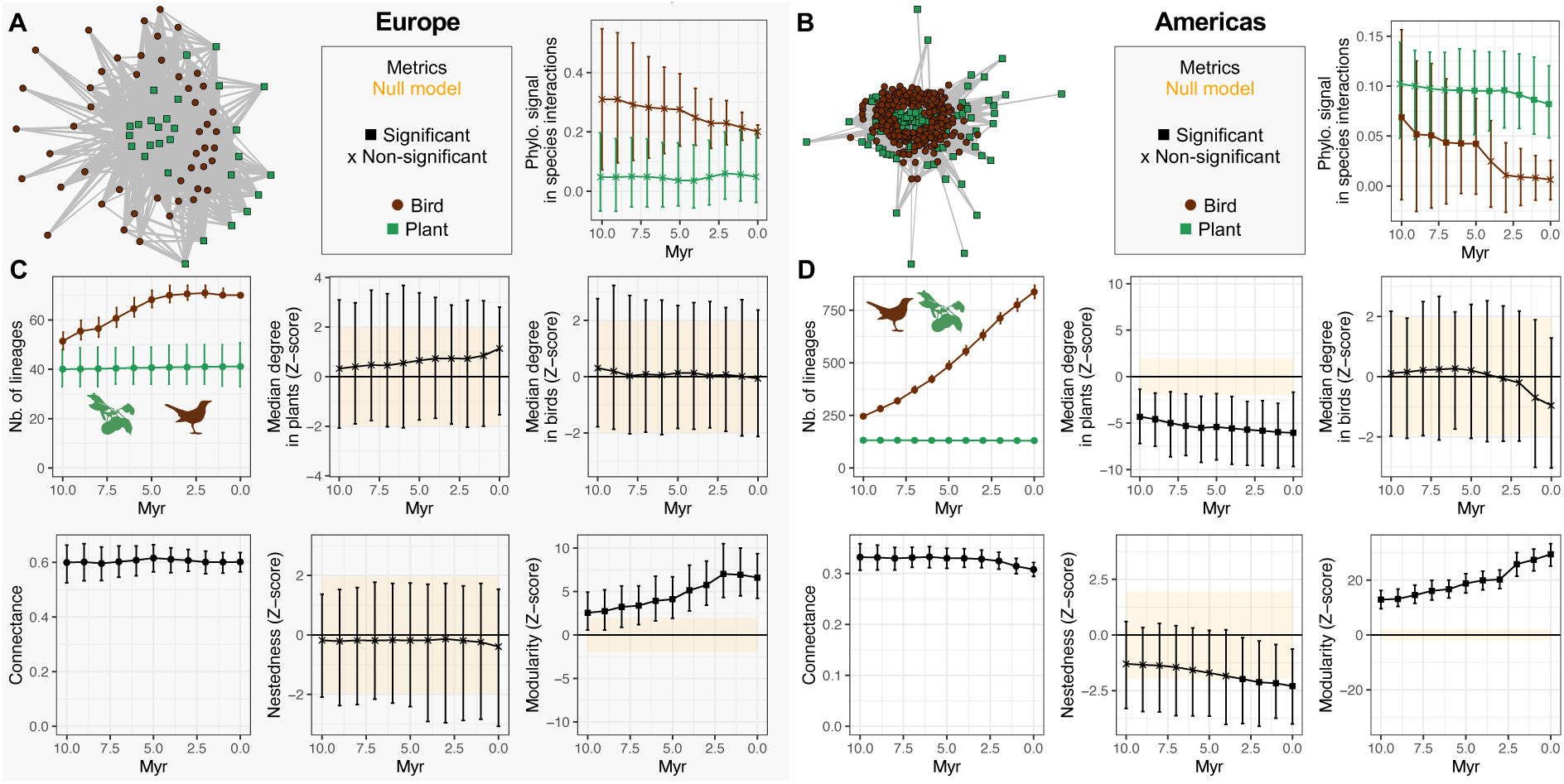
Ancestral network reconstruction of bird seed dispersal. Consensus ancestral networks inferred 10 Myr ago for the European (A) and American (B) bird seed dispersal networks. Ancestral network metrics (C & D). Squares indicate significant, and crosses non-significant departure from the null model (shown in orange), as in Figs. 3 & 4.

Our ancestral networks indeed reveal that angiosperms have preferentially formed associations with Glomeraceae, and non-angiosperms with early-diverging Glomeromycotina fungi, a pattern that has persisted since the origin of flowering plants (>150 Mya; Fig. 3A-C, Table S2). This tendency of flowering plants to interact with Glomeraceae (“recent plants with recent fungi”) and non-flowering plants with non-Glomeraceae (“ancient plants with ancient fungi” (41)) has emerged more than 150 Myr, at the emergence of flowering plants, reinforcing the idea of an entangled diversification dynamic in these clades (42, 43). Similarly, modularity was already evident in bat pollination networks 10 Mya, corresponding to the divergence of bat subfamilies (Fig. 4). Compared with other mutualistic systems, seed dispersal networks exhibited elevated connectance (44), a pattern already present 10 Mya (Fig. 5A-B). Only a minority of taxa are specialists (Fig. 5C-D), yet the networks present a significant modularity, potentially shaped by the partial phylogenetic conservatism of functional traits that mediate frugivory (45).

The use of latent traits is a strength of our approach, as it allows reconstructing ancestral networks without requiring the formulation of specific rules for the evolution of interactions as clades diversify, or large datasets of traits putatively determining who interacts with whom. The drawback is that the latent traits lack direct biological interpretation and do not necessarily capture the effect of interaction traits *per se*. By construction, latent trait values likely reflect, to some extent, the degree of generalism, as two species are more likely to interact if they exhibit high values for the same latent traits. Consequently, generalist species are expected to exhibit higher latent trait values. Latent traits may also capture biogeographic effects if species tend to be generalist and biogeography plays a major role in determining which species interact. Therefore, the evolutionary conservatism of latent traits may reflect the conservatism of specialization or biogeographic distributions, rather than directly reflecting that of interaction traits. In an attempt to better interpret the biological meaning of latent traits in the mutualistic networks studied here, we investigated their associations with the degree of generalism and biogeography. We indeed detected a strong association between the sum of latent trait values and species degree across all networks, except for the European bird seed dispersal network (Fig. S9). In the three other networks, generalist species exhibited higher latent trait values on average, whereas specialist species tended to have lower values. The association of latent traits with species’ bioregions was quite weak in the arbuscular mycorrhizal and bat pollination networks (Fig. S10). Although this association was stronger in the bird dispersal networks, inferred modules were, on average, composed of species spanning two to four bioregions (Table S3), indicating that biogeography alone cannot explain the observed modular structure.

## Discussion

Present-day large-scale networks result from millions of years of coevolution among interacting species, but we lack direct observations of their deep-time evolution. We addressed this issue by developing a phylogenetic approach to reconstruct ancestral interaction networks. Ancestral continental and global networks of plant mutualisms involving arbuscular mycorrhizal fungi, bat pollinators, and bird seed dispersers show a consistent modularity structure that has persisted over macroevolutionary time scales.

The main idea underlying our approach, compared with previous attempts (29), is to introduce latent traits supposed to determine who interacts with whom, and to reconstruct these latent traits rather than directly the interactions. This is justified by the fact that interspecific interactions generally require a minimal level of compatibility between morphological, physiological, phenological, and/or behavioral traits to establish (1, 46). Hence, ideally, latent traits partially reflect biological characteristics that shape biotic interactions (4, 31). To estimate these latent traits, we used a singular value decomposition of the present-day interaction matrix, following (31, 32). By construction, the latent traits are therefore independent, determine different axes of variation in present-day interactions, and two species are more likely to interact if they exhibit high values for the same latent traits (47, 48). While this interaction rule does not match traditional formulations of trait matching or phenotypic differences, we showed using simulations that the approach reliably recovers ancestral interactions across diverse simulated scenarios of network evolution when species interactions are evolutionary conserved, demonstrating the flexibility of latent traits in capturing who interacts with whom. We of course do not suggest that trait matching is not important in mutualism, but trait matching rules are often violated (e.g., small pollinators that do not use some flowers with short floral corolla, and to the contrary use flowers with long floral corolla tubes), such that a specific single-trait trait matching scenario is often biologically unrealistic. In contrast, the multidimensional latent trait space captures the heterogeneity of factors shaping mutualistic interactions.

Latent traits reflect diverse aspects beyond specific interaction traits. For example, latent traits correlate with species degree, which can reflect generalism, abundance, or sampling structure (49). At the scale considered in the paper, latent traits could also reflect biogeography, which can constrain co-occurrence, and therefore potential interactions (50–52), although we did not find a strong biogeographic signal in the latent traits. Latent traits could also reflect the combined expression of pleiotropic genes required for interactions, such that interaction can occur when complementary gene sets are highly expressed in both species. Regardless of their specific biological interpretation, which is likely complex and multifactorial, latent traits provide an operational and flexible framework to characterize who is likely to interact with whom.

The introduction of latent traits greatly simplifies the ancestral network reconstruction problem, as ancestral trait values can easily be obtained on the phylogenies of the interacting guilds under classical models of trait evolution. In addition, the independency of the traits guaranteed by the singular value decomposition means that traits can be assumed to evolve independently. We used here independent Brownian motion processes, a general framework for trait evolution (53) that has previously been used to infer unknown present-day interactions (32). The Brownian process is prone to smoothing, which may explain the observed tendency towards homogenization of the structural properties of reconstructed ancestral networks (*i.e.* under-estimation of high metrics and over-estimation of low ones). Other models of trait evolution could be used if more appropriate; for example, Ornstein-Uhlenbeck processes may better reflect processes of stabilizing selection and phenotypic convergence operating in mutualisms (6), and models with lineage-lineage trait coevolution (54) may better reflect the establishment of mutualistic syndromes – coevolved sets of complementary traits that tend to stabilize beneficial interactions – which are well-documented in the case of pollination ((55) but see (45)). Future developments could also explicitly account for biogeographic range evolution and restrict potential interactions to species co-occurring in the same region (as, for example, in (56)).

Another strength of our approach is to explicitly account for extinct and unsampled lineages. While we found that our results are robust when not augmenting the trees, neglecting these missing lineages results in a large undersampling of past diversity, and could affect measures of ancestral network structure (57). We used here a constant-rate homogeneous birth-death model to complement the reconstructed phylogenies. This resulted in an increase in the number of reconstructed lineages through time, except for plant families in the bat pollination and bird seed dispersal networks. Our framework could take advantage of recent developments that provide “augmented trees” under more realistic diversification models with heterogenous speciation and extinction rates through time and across lineages (39, 58), some of which can integrate information from the fossil record (59, 60). This would improve ancestral reconstructions and could allow accounting for mass extinction events, which may have strongly affected ancestral network structures (61). Considering “geographic” birth-death models, such as GeoSSE (62) or extensions (63) could in the future allow accounting for immigration and extirpation events that add or remove species from the geographic region considered, although such models have not yet been implemented with data augmentation. This step is necessarily imperfect given the current stage of development of phylogenetic diversification models, yet it is an improvement over ignoring missing lineages in phylogenies altogether.

Despite the flexibility of our approach, its performances vary: It performs best when modifications in the interactions occur anagenetically rather than cladogenetically, and when interactions are evolutionary conserved. When frequent interaction shifts occur, ancestral species are often inferred to interact while they do not (Fig. S4). Our simulations obviously do not (and cannot) cover all the processes known to play a role in the evolution of mutualisms, such as convergence (9). Yet, they mimic diverse scenarios that can be found in nature and result in a wide range of evolutionary conservatism, from absent to strong (25), thereby allowing us to draw conclusions beyond the specific scenarios considered. For example, we can expect the approach to perform well even in the presence of convergence, if this convergence does not entirely erase phylogenic signal, and to not perform well in the opposite case. Although this is typically not known *a priori* for a given empirical system, our cross-validation procedure provides a general and robust method for assessing the reliability of the ancestral reconstructions for any empirical system. If the cross-validation score falls below 0.25, mutualistic interactions may be too evolutionarily labile or too strongly shaped by convergent evolution to be reliably inferred in the past using ELEFANT. If the score exceeds 0.25, the reconstructions can be trusted at least in the recent past. Indeed, we found in this case that F1-scores of ancestral interactions in the recent past were above 0.5. It may *a priori* seem surprising to find better F1-scores in the recent past than in the present-day cross-validation, but this is in fact expected from the phylogenetic reconstruction, as estimates for “recent” internal nodes benefit from more direct observed information (from their descendants) than unsampled phylogenetic tips. The estimates are however less precise as we go deeper into the past, as expected from all ancestral reconstruction techniques based on stochastic processes. Phylogenetic uncertainty, which also increases further back in time, also reduces confidence in the reconstructions. As a rule of thumb, we would not consider ancestral networks constructed past halfway through the clade’s age to be reliable. Importantly, our cross-validation framework assumes that the present-day rules underlying predictions of extant interactions have not changed through time. For example, we assume a fixed threshold in the product of latent traits above which interactions occur. Although our simulations show that the results are robust to threshold choice, this threshold may have varied over evolutionary time due to intrinsic or environmental factors that facilitate or constrain interactions. We therefore recommend focusing on relatively recent timescales, as in our empirical applications. Ultimately, deep-time inferences of ancestral interactions predicted by ELEFANT will require validation using independent lines of evidence, such as fossil records of interactions.

Cross-validation scores obtained on the four empirical mutualistic networks we analyzed are relatively low, although above the 0.25 threshold at which we consider structural metrics to be trustable. These modest scores are primarily related to false positives, which may in fact correspond to true but undocumented interactions (36). Indeed, empirical surveys often under sample interactions despite intense sampling efforts, and the latent-based framework has in fact precisely been used to estimate missing, unobserved interactions (36). Additional work would be required to assess to which extent false positives are in fact real, *versus* artifacts due, for example, to the fact that biotic interactions are more ecologically labile, context-dependent, and plastic than intrinsic species traits (*e.g.* morphological traits) (18, 64).

The global or continental ancestral networks of plant-mycorrhizal, pollination, and seed dispersal mutualisms reveal a remarkably stable modular structure in the last millions of years, despite continual changes in species composition driven by speciation and extinction, and variations in environmental conditions. These results were obtained by considering plants at the family level, given scarce interaction data at the species level and computational limitations. The networks also had the tendency to be phylogenetically structured, with closely related species interacting with similar subsets of partners. Patterns related to the median species degree were more contrasted, with a slight but inconsistent tendency toward specialization. Finally, nestedness was globally not distinct from null expectations. Hence, we report in all plant mutualisms a stable modularity with a certain degree of phylogenetic conservatism. Such signal has been reported previously (46), but we show here that it is an evolutionarily stable characteristic of these mutualistic networks, rather than a feature that emerged only on recent ecological time scales. These results are not an artifactual consequence of using a phylogenetic inference method: although ELEFANT assumes Brownian evolution of the latent traits, and thus a certain degree of phylogenetic conservatism, its cross-validation procedure evaluates whether this assumption is accurate. In the absence of phylogenetic signal, cross validation scores would be low. The structural characteristics of local mutualistic networks, which tend to be more nested (5) and to exhibit moderate phylogenetic signal (45, 65), may be quite different and potentially less evolutionary stable. At large geographic scales, our results suggest that selection favors the retention of existing beneficial associations over macroevolutionary time scales (20), even if biogeographic barriers may also play a role. This in turn may favor diffuse coevolution, reinforcing reciprocal specialization and compartmentalization (21). The extent to which the latent traits inferred by ELEFANT correspond to classical mutualistic syndromes remains an open question (55) and could be explored in future work, for instance by correlating the latent trait values with measurable species traits that influence interactions (45). Across the different mutualistic networks, the influence of biogeography on network assembly was heterogeneous. It was rather weak in arbuscular mycorrhizal and bat pollination networks, and slightly more pronounced in bird seed dispersal networks. Although comparisons should be made cautiously given the differing spatial scales of these systems, our results support the idea that biogeography only weakly contributes to shaping mutualistic network assembly (11). Our findings provide further support for the stability of mutualisms across macroevolutionary timescales associated in part with niche conservatism in species interactions (16, 25, 34). Such evolutionary stability of mutualistic network structure over millions of years now stands in sharp contrast to its rapid erosion under human pressures, which are driving an unprecedented, worldwide homogenization of ecological communities (11, 66).

By reconstructing ancestral networks, we can follow Darwin’s ‘entangled bank’ through deep time. Tracing back the mutualistic side of this entangled bank suggests that the benefits of retaining established mutualistic relationships outweigh its costs, resulting in stable modules of relatively closely-related species. Similar analyses of the antagonistic side may reveal much more instability, with Red Queen dynamics driving frequent partner shifts. This deep-time perspective on interspecific interactions lays the groundwork for understanding their role in species diversification – that is, how the entangled bank modulates global biodiversity patterns.

## Methods

### The ELEFANT model

We consider two interacting guilds composed of *p* plant and *m* mutualist species sampled at present (‘plants’ and ‘mutualists’ are used here for simplicity, but our approach applies to any interacting groups of organisms, including antagonistic interactors). The binary matrix *N* (*p*, *m*) indicates whether each pair of plant and mutualist species interact (1) or not (0). The extant, sampled species of each guild have an associated reconstructed, time-calibrated phylogeny. We developed a framework, the “Evolution of LatEnt traits For Ancestral NeTwork reconstruction” (ELEFANT), illustrated in Fig. 1 and detailed in the next paragraphs, that infers ancestral network from these combined network and phylogenetic data. The framework proceeds in five steps, described below.

#### i) Inferring present-day latent traits

First, assuming that interactions are shaped by unobserved traits, we obtain the theoretical values of these latent traits for each extant species using the random dot product graph (RDPG) method (31, 32)(Fig. 1.1). In previous developments, the approach was used to infer present-day interactions of species for which interacting partners were not known by inputting latent trait values to extant species, *i.e.* phylogenetic tips (32, 40), or to augment unobserved, missing interactions within ecological networks (36). Here, we aim to infer ancestral interactions. This method performs a singular value decomposition of the present-day interaction matrix *N* by finding the matrix Σ (*p*, *m*) with null non-diagonal elements, and the matrices *L* (*p*, *p*) and *R* (*m*, *m*), such that *N* = *L* × Σ × *R^t^*. Each diagonal element of the Σ matrix corresponds to a singular value associated with a given share of the variance observed in present-day interactions. By construction, these singular values are independent and each of them defines an independent axis of variation of a latent trait. We selected the *d* first eigenvalues (Σ|*d*) that explain at least 90% of the total variance observed in present-day interactions. The values of the *d* independent latent traits associated to the *p* plants are given by the *p* × *d* matrix 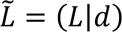, and those associated to the *m* mutualist s are given by the *d* × *m* matrix 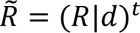. Each species is thus characterized by a vector of *d* latent trait values. The interaction matrix *N* is approximated by 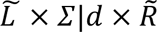. Thus, the sum of the products of each latent trait between plant *i* and mutualist 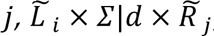, reflects their likelihood of interaction.

In other words, the probability of interaction between two species increases when both species exhibit high values for the same latent traits. Consequently, specialist species tend to have on average lower latent trait values, whereas generalist species tend to have higher latent trait values. To determine the threshold *J* above which we can consider that there is an interaction between two species, we use a binary classification approach between the observed interaction network (*N*) and its approximation 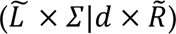 (67): we take the *J* value maximizing the Youden’s *J* statistic, defined as *J* = sensitivity (percentage of interactions correctly identified) + specificity (percentage of non-interactions correctly identified) – 1. This value is thus specific to each network.

#### ii) Augmenting phylogenetic trees

The second step (Fig. 1.2) is required to account for lineages that were likely involved in ancestral interactions but are missing in reconstructed trees, which lack all the lineages that left no descendants at present, either because they went extinct or were not sampled. For simplicity, we consider that the plant and mutualist phylogenies represent the diversification of two clades that follow each a constant-rate, homogeneous birth-death process, and we implement a data augmentation approach to simulate extinct and unsampled lineages on the plant and mutualist phylogenetic trees. Given the speciation rate *λ*, the extinction rate *μ*, and the sampling fraction at present *ρ*, all three assumed to be constant and known (in practice, *λ* and *μ* are estimated by maximum likelihood), we augment the phylogenetic trees with extinct and unsampled lineages (see Supplementary Methods). We account for stochasticity by independently replicating this data augmentation step 250 times. In the following steps, we thus work with a set of ‘augmented’ phylogenetic trees for each clade, which represent probable diversification histories. The assumption that the phylogenies on each side of the network represent the diversification of two clades following each a birth-death process corresponds best to a scenario where diversification occurs *in situ* at the spatial scale of the network, as in large (continental to global scale) networks over short enough timescales, or in close communities of *in situ* radiations, such as those found on remote islands or lakes. The assumption of a constant-rate homogeneous process can be relaxed with existing phylogenetic diversification models implemented with data augmentation (39, 58).

#### iii) Inferring ancestral latent traits

Next, for each pair of plant and mutualist augmented trees, we perform ancestral state reconstructions to estimate the latent trait values along each lineage (Fig. 1.3). We assume that the *d* latent traits of the plants (*L̃*) and their mutualists (*R̃*) evolved according to independent Brownian motion processes. For each latent trait, we infer the parameters of the Brownian motion process using maximum likelihood (function mvBM in the mvMORPH R-package (68)). Then, at each time point on the augmented phylogenetic trees (including at present), we determine the expected value of each latent trait and their variances (69, 70) and perform 250 stochastic character mappings to explicitly propagate uncertainty in ancestral trait reconstruction.

#### iv) Inferring ancestral networks

In the fourth step, we reconstruct the ancestral network *N*(*t*) between coexisting species at any given time *t* in the past (Fig. 1.4), based on the likelihood that two species interact given their inferred latent trait values 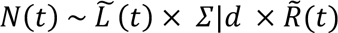 and according to Youden’s *J* threshold (see above). Note that this same procedure can be applied to infer the probable set of interacting partners for present-day species which interacting partners are not known. In the case of obligatory interactions, where a species cannot survive without at least one partner, we force every species to form at least one ancestral interaction: if the Youden’s statistic is not reached with any partner, we choose the partner species with the highest likelihood of interaction. This only happens in a limited number of cases (<2% of the species in the empirical applications).

#### v) Cross-validations

The last step of ELEFANT aims to provide a measure of confidence in the ancestral network reconstructions, given the strong assumptions made throughout: i) interactions determined by latent traits derived from a truncated singular value decomposition at present, ii) latent traits evolving according to independent Brownian motion processes, iii) latent traits, and therefore interactions, inherited at speciation, and (iv) ancestral lineages inferred to have interacted when the product of the latent traits exceeds the Youden’s J threshold, assumed to be fixed over evolutionary time across the clades. We use cross-validation on the observed present-day interaction data to obtain a measure of confidence in our model predictive accuracy (Fig. 1.5): we randomly subsample 90% of the extant species, predict the partners of the other 10% (using ancestral trait reconstruction at present), repeat this procedure 100 times, and compute the average F1-score, referred to as the “cross-validation score”, defined as the harmonic mean of the sensitivity (percentage of interactions correctly recovered) and the precision (the ratio of recovered interactions corresponding to true interactions). In other words, in this cross-validation step, we use ancestral trait reconstructions to predict the present-day interactions of the subsampled species, and test whether they match observed interactions.

Based on the results obtained across the different simulated scenarios, we adopt a cross-validation score of 0.25 as the operational threshold above which ancestral reconstructions can be considered reliable at least in the recent past (Fig. S3). Although this threshold may appear low, it reflects the intrinsic difficulty of our cross-validation design, in which ELEFANT must reconstruct ancestral interactions from present-day sampled species and subsequently predict present-day interactions of unsampled species from those ancestral reconstructions – that is, present to past to present. By contrast, the primary application of ELEFANT involves the reconstruction of ancestral interactions from present-day data (present to past), a task that consistently yields substantially higher F1-scores according to our simulations and thus greater reliability (see simulation results in Figs. 2A and S3).

Taking as input an interaction matrix (the network) and associated time-calibrated phylogenies, ELEFANT thus outputs at each time *t* in the past a set of ‘augmented’ ancestral networks representing probable interactions between both lineages present in the phylogeny and augmented lineages, as well as a measure of confidence in these reconstructions. When the two phylogenetic trees have different ages, ELEFANT provides reconstructions until the most recent common ancestor of the youngest clade.

In summary, ELEFANT’s overarching objective, *i.e.* reconstructing ecological networks through evolutionary time, parallels that of Braga et al. (29), but uses an orthogonal approach. Importantly, ELEFANT models the evolution of the two interacting clades jointly, whereas the approach of Braga et al. (29) considers a fixed set of host repertoires (*e.g.* plant families) that are gained or lost as the interactor clade (*e.g.* butterflies) diversifies. ELEFANT’s integration of latent traits with phylogenetic reconstruction is closely aligned with the framework proposed by Strydom et al. (32), who introduced a latent trait approach for phylogenetically informed transfer network representations (extending earlier work by Dalla Riva & Stouffer (31)), but used this framework to reconstruct present-day rather than past interactions. Important specificities of ELEFANT are to explicitly account for unsampled and extinct species to avoid undersampling ancestral networks, and to incorporate a cross-validation procedure to assess the confidence we can have in the reconstructions. Finally, in contrast to Strydom et al. (32), ELEFANT explicitly propagates uncertainty in ancestral trait reconstruction by performing stochastic character mappings based on the theoretical distributions of node states under a phylogenetic Brownian motion model.

### Summarizing ancestral networks

We summarized the structure of augmented ancestral networks by reporting several global metrics: i) median species degree, *i.e.* the median number of mutualists interacting with each plant and *vice-versa*, ii) connectance (the ratio of realized interactions), iii) nestedness (the tendency of specialist species to interact with generalist partners, measured using the NODF2 index (71)), iv) modularity (the density of interactions within modules relative to a null expectation – computed using the function computeModules (71)), and v) the phylogenetic signal of species interactions in each guild, *i.e.,* the tendency of closely related species to interact with similar partners, measured using Mantel tests with the function phylosignal_network (48). This function calculates a Pearson correlation between phylogenetic distances and binary Jaccard distances, with significance assessed using 10,000 permutations.

We also generated consensus ancestral networks constituted of frequently recovered interactions between pairs of species (30). We added an interaction in the consensus network if it was recovered in at least x% of the augmented ancestral networks, with x being either 10%, 25%, 50%, or *J*% (Youden’s *J* threshold). Consensus networks do not contain interactions involving augmented species, as these vary across augmented trees.

### Simulations

To evaluate the accuracy and robustness of ELEFANT in reconstructing ancestral networks, we simulated the evolution of plant-mutualist interaction networks using five simulation designs that capture different scenarios of species interactions evolution. We designed them based on different assumptions about how interaction networks could evolve, with the aim to obtain realistic simulated networks that resemble empirical ones. Although additional, more specialized simulations could have been developed — for example, incorporating convergent evolution in mutualisms, codiversification, mass extinction events, or preferential extinctions of key hub species — our goal was not to exhaustively test ELEFANT across all possible modes of mutualism evolution. Instead, we sought to characterize its performance under a range of scenarios differing in the degree of evolutionary conservatism of species interactions (see below). For systems in which ongoing processes not represented in our simulations are suspected (*e.g.* convergent evolution), the cross-validation procedure (see above) provides a way to assess the ability of ELEFANT to recover subsampled interactions directly from the empirical dataset, without requiring additional simulations.

As in the main text, we use “plants” and “mutualists” to refer to the two interacting guilds for simplicity and to match our empirical examples, but these simulations can equally represent the evolution of any two interacting guilds, including those engaged in antagonistic interactions.

In each of the five scenarios, we simulated network evolution for 30 million years and recorded the simulated ancestral networks every 5 million years. Both the plants and the mutualists diversified according to a constant rate birth-death process with a speciation rate (λ) uniformly sampled between 0.05 and 0.2 and an extinction rate (μ) between 0.01 and 0.05. We chose these rates to span the range of empirical diversification rates reported in the literature (38). For computational reasons, we only kept simulated systems having between 25 and 150 extant species per guild. We generated a total of 200 simulations per scenario.

#### i) Evolution by anagenetic shifts

We started the process with the diversification of plants, with that of the mutualists starting 10 million years after; we considered that the most recent common ancestor of the mutualists interacted with one single plant lineage picked at random. Then, we assumed that interactions are inherited at speciation events (no cladogenetic changes, the two descendant lineages inherit the interacting partners of their ancestor) but can change during the life time of the lineages (anagenetic shifts): a mutualist lineage can acquire a new interaction with an additional plant species chosen at rate 0.015 (number of events Myr^-1^), or lose one of the plant species it interacts with at a rate of 0.015. The plant partners that are gained or lost are picked at random. When a lineage goes extinct, the species it interacts with lose their partner. If this was the only interaction of the mutualist, a new interacting plant lineage is picked at random. By construction, mutualists are obligately associated with at least one plant species, while the plants are only facultatively interacting with the mutualists (*e.g.* while plants may form mycorrhizal associations facultatively, arbuscular mycorrhizal fungi are obligate symbionts).

#### ii) Evolution by cladogenetic shifts

By modeling the evolution of latent traits using Brownian motions, ELEFANT assumes that the latent trait values are inherited at speciation and therefore that the two daughter species are likely to keep the same interactions. However, there are examples of closely related species that likely switched partners at speciation (72, 73) and such cladogenetic shifts may not be well handled by ELEFANT. We therefore simulated a scenario where network evolution happened through both anagenetic and cladogenetic shifts. We built upon the scenario “evolution by anagenetic shifts” with a rate of anagenetic acquisition of 0.15 and a rate of anagenetic loss of 0.05 and added cladogenetic shifts. At each mutualist speciation event, one of the two daughter lineages retains the same interacting plant species, while the other daughter lineage forms an entirely new set of interactions with randomly selected plants.

#### iii) Brownian evolution of latent traits

Next, we considered one simulated scenario corresponding to the underlying model of ELEFANT. We simulated the independent evolution of *d*=10 latent traits using Brownian motions on the plant and mutualist phylogenetic trees with 0 as ancestral values and a variance of 0.01. We attributed interactions between pairs of plant and mutualist species if the product of their latent traits was larger than 0.25. Here, both plant and mutualist species were considered to be obligate partners, a condition often met in mutualistic networks (*e.g.* pollination).

#### iv) Host repertoire evolution

In this scenario, we assumed that host taxa (plants) diversified before the most recent common ancestor of the mutualists (*e.g.* it can correspond to higher taxonomic levels, *e.g.* plant families) and we tracked the set of host taxa associated with each mutualist species during their diversification. Following Braga *et al.* (29), we referred to this set of associated host taxa as the host repertoire. Each mutualist obligatorily interacts with at least one host taxon, while the hosts only facultatively interact with mutualists. We started the simulation by randomly attributing one host taxon to the most recent common ancestor of the mutualists. Then, the host repertoire of each mutualist lineage evolved independently by acquiring new host taxa at a rate of 0.15 and losing host taxa at a rate of 0.05 following a continuous-time Markov process. Host repertoires are inherited at mutualist speciation.

#### v) Random evolution

Finally, we considered a scenario of fast interaction evolution where network evolution occurs by anagenetic shifts at very high rates, such that interspecific interactions are not conserved over macroevolutionary timescales. To simulate this scenario, we used the first simulation scenario with a rate of plant acquisition of 100 and a rate of plant loss of 200.

We assumed that not all extant species were sampled at present and used a sampling probability (ρ) uniformly sampled between 60% and 100%, as undersampling is a common phenomenon in empirical networks (57). We discarded extinct and unsampled species from the phylogenetic trees of each guild; we referred to these new trees as the ‘reconstructed trees’. In addition, we considered that not all interactions are sampled at present and that present-day interactions between sampled species have only a 90% chance of being observed while keeping at least one observed interaction per species. We referred to these subsampled present-day networks as the ‘observed networks’.

The values of the parameters used in the different simulations were chosen to obtain a range of realistic simulated networks mimicking the variability observed in empirical ones. Indeed, we obtained ranges of size, connectance, modularity, and nestedness values comparable to those of empirical networks (Fig. S5)(44). We also obtained a range of phylogenetic signals in species interactions (Supplementary Results, Fig. S11), with many networks showing low signals as often observed in nature (65).

We then ran ELEFANT on each simulation composed of one observed network and two reconstructed trees. For the step of data augmentation of extinct and unsampled lineages, we used the simulated parameters values λ, μ, and ρ, and generated a total of 250 augmented ancestral networks. Between 5 and 15 singular values were often necessary to explain at least 90% of the present-day interactions in the different types of simulated networks (Supplementary Results; Fig. S12).

We then compared the simulated *versus* inferred global metrics of the ancestral networks (number of species, connectance, nestedness, and modularity) by fitting linear models. We also evaluated whether the analyses recovered significant phylogenetic signals when present in the simulations. Finally, we looked at the inferred ancestral interactions present in the consensus ancestral networks and reported the sensitivity (percentage of interactions correctly identified), the percentage of interactions corresponding to false positives, and the corresponding F1-score. We replicated the analyses without data augmentation (μ=0 and ρ=1) and confirmed that adding extinct and unsampled lineages significantly increased the performance of ELEFANT (Supplementary Results; Fig. S13).

Finally, to assess the robustness of our thresholding approach based on Youden’s threshold, we conducted additional simulation analyses using the anagenetic scenario (scenario i). We applied ELEFANT across a wide range of thresholds (0.1 to 0.9, every 0.1) and evaluated performance using (i) the accuracy of global network properties (connectance, nestedness, and modularity), quantified as the R² between simulated and inferred values, and (ii) the accuracy of ancestral interaction reconstruction, quantified using F1-scores (see Supplementary Results; Fig. S14).

The implementation of ELEFANT and all simulations were performed in R (74).

### Empirical analyses

We applied ELEFANT to three different mutualistic systems involving plants and covering a broad range of temporal and spatial scales: the global-scale arbuscular mycorrhizal symbiosis, the pollination by bats in the Neotropics, and the seed dispersal by birds in Europe and the Americas. Although we presented ELEFANT as a species-level model applied to monophyletic clades, it can technically be applied at any taxonomic level (*e.g.* genus or family), and to non-monophyletic groups of organisms, as long as the taxa included in the network and in the phylogenetic trees match. Here, in order to reduce “missing” data (e.g. plant species which mycorrhizal partners have not been characterized) and better match the type of network structures obtained in our simulations, we considered plants at the family level. In addition, plant mutualists correspond to ecologically- and/or geographically-defined groups of taxa (*e.g*. nectar feeding bats, bird seed dispersers in Europe). The “augmentation” step of the framework should thus be considered as an imperfect attempt to account for missing lineages, rather than as a rigorous reconstruction of past diversification and diversity histories (as, in, *e.g.* (39, 58)). For example, the birth-death model is not ideal to represent the (dis)appearance of higher taxa (see, *e.g.* (75) for a better model), and only makes sense in the case of ecologically- and/or geographically-defined groups of taxa for which it can be assumed that ecological and/or geographical transitions have been rare in the time period over which networks are reconstructed. We limited our continental-scale reconstructions (for bats and birds) to relatively recent time periods (on the order of 10 Myr), on which the latter assumption is reasonable. This does exclude the possibility that important immigration events occurred (for example, the thrushes arrived in the Americas between 11 and 4 million years ago, (76)), but these rare events are unlikely to deeply affect the global structure of ancestral networks. Given these imperfections in the reconstruction of ancestral lineages, which could affect the reconstruction of ancestral networks, we assess the robustness of our results to choices made during the data augmentation step.

First, we reconstructed ancestral interaction networks of the arbuscular mycorrhizal symbiosis between land plants and Glomeromycotina fungi at the global scale. We gathered the interaction data from the MaarjAM database ((77); https://maarjam.ut.ee) filtered according to (13). We considered the network at the levels of virtual taxa (VT) for the Glomeromycotina (391 taxa delimited using DNA-based criteria; (77)) and plant families (115 families). We used a phylogenetic tree of Glomeromycotina from (42) and reconstructed the plant phylogeny by pruning the mega-tree of vascular plants using V-PhyloMaker2 (78). We ran ELEFANT with λ=0.012, μ=0.001, and ρ=0.86 for Glomeromycotina following the estimates of (42), and λ=0.02, μ=0.003, and ρ=0.19 for the plants (the two former parameters were estimated by fitting a constant-rate birth-death model in RPANDA on the family-level phylogeny of land plants (79) with a sampling fraction computed as the number of sampled plant families (115) divided by the estimated total number of plant families (600)). We looked at the ancestral networks for the past 150 Myr, which corresponds to the emergence of Angiosperms, and respects the “halfway through clades age” rule of thumb. At this temporal scale and with the birth-death models used in our data augmentation procedure (constant rates, with a net diversification rate of 0.011 and 0.017 for the Glomeromycotina and plants, respectively), we expect to infer a comparable increase in the number of lineages through time on both sides of the network.

Second, we investigated the evolution of plant-bat pollination networks. We gathered the interaction data from (80). We focused our analysis on the evolution of nectar-feeding bat pollinators from the Phyllostomidae family, a Neotropical bat family including many nectar-feeding species. We obtained the phylogenetic tree of the 56 bat species from (81) and used V-PhyloMaker2 for the phylogenetic tree of the 42 plant families. We ran ELEFANT with λ=0.20, μ=0.01 (81), and ρ=0.31 for the bats (obtained by assuming that there are 180 extant species of the nectar-feeding Phyllostomidae family (82)) and λ=0.02, μ=0.003, and ρ=0.63 for the bat-pollinated plants (83). We looked at the ancestral networks for the past 12 Myr, which is less than half the age of the Phyllostomidae family. At this temporal scale and with the birth-death models used in our data augmentation procedure (constant rates, with a net diversification rate of 0.19 and 0.017 for the bats and plants, respectively), we expect to infer an increase in the number of lineages through time on the bat side of the network one order of magnitude faster than on the plant side of the network. This reflects the consideration of plants at the family level, and associated expectation that very few families emerged during a 10 Myr period.

Third, we studied seed dispersal of fleshy fruits by birds in Europe and the Americas. We obtained the plant-bird interactions from (11), the bird phylogenetic tree from (84), and the plant phylogenetic tree from V-PhyloMaker2. Europe and Americas were picked as they formed two well-sampled, separated modules in the global-scale network of seed dispersal by birds (11). In Europe, the network contained 70 bird species and 28 plant families forming fleshy fruits, with sampling fractions (ρ) of 1 and 0.78, respectively (85, 86). In the Americas, it contained 586 bird species and 115 plant families forming fleshy fruits, with sampling fractions (ρ) of 0.73 and 0.93, respectively (85). We ran ELEFANT with λ=0.15 and μ=0.01 for the birds (87) and λ=0.02 and μ=0.003 for the plants. We looked at the ancestral networks for the past 10 Myr, which limited the number of potential bird immigration events while still encompassing a significant number of speciation events. Under our birth–death model, the speciation rate exceeds the extinction rate, so we expect an increase in the number of bird lineages toward the present over the last 10 million years. At this temporal scale and with the birth-death models used in our data augmentation procedure (constant rates, with a net diversification rate of 0.14 and 0.017 for the birds and plants, respectively), we expect to infer an increase in the number of lineages through time on the bird side of the network one order of magnitude faster than on the plant side of the network. This again reflects the consideration of plants at the family level, and associated expectation that very few families emerged during a 10 Myr period.

The three empirical networks showed different levels of phylogenetic signal. In the arbuscular mycorrhizal symbiosis, significant signal was detected in both plants (R=0.30, p<0.001) and fungi (R=0.11, p<0.001). Similarly, the pollination network exhibits significant phylogenetic signal in both plants (R=0.13, p=0.01) and bats (R=0.12, p<0.001). In contrast, in the European seed dispersal network, closely related birds interact with similar plant families (R=0.21, p<0.01), but not the reverse (R=-0.01, p=0.50), and no significant signal was observed in the American seed dispersal network (p-values>0.05). In these mutualistic networks, respectively, 38, 12, 4, and 18 latent traits were necessary to explain at least 90% of the present-day interactions.

To evaluate the significance of the structural properties of each augmented ancestral network (nestedness, modularity, median species degree, and phylogenetic signal), we compared the original metrics with those obtained under three null models. These null models account for the intrinsic relationships between network size, connectance, and other network properties by preserving some characteristics while allowing others to vary. To evaluate the significance of nestedness and modularity, we used the “quasiswap” null model (function *nullmodel* in the R-package vegan), which randomizes the interactions while maintaining network connectance and species degrees (*i.e.* the marginal sums of the networks). To evaluate the significance of the median species degree, following (30), we used the less stringent “r00” null model, which randomizes the interactions while maintaining only network connectance. Finally, to evaluate the significance of phylogenetic signal, we shuffled the tip labels in the phylogenetic trees (48). The first two null models were each replicated 250 times, whereas the third, being less computationally demanding, was evaluated using 1,000 replicates. Beyond assessing significance, we calculated Z-score-transformed metrics from the null models and focused exclusively on temporal variation in these standardized metrics to avoid directly comparing raw values across ancestral networks, which are strongly influenced by variation in network size and connectance.

To evaluate the robustness of our results, we reran ELEFANT without using data augmentation or with other rates of diversification for the step of data augmentation (Table S4; Fig. S8).

We tested the sensitivity of our findings to the use of Youden’s J as the interaction threshold by repeating the analyses using a range of fixed thresholds: 0.5, which showed the best performance in the simulations, as well as the lower and upper bounds of the well-performing range, 0.3 and 0.6 (Fig. S14). We found qualitatively similar results across all thresholds (Fig. S15).

We performed additional analyses to (i) assess the influence of biogeography on latent traits and (ii) quantify the extent to which biogeography explains the phylogenetically-conserved modularity observed in the networks (see Results). Bioregions for each species were assigned based on the original publications: major biogeographic realms for arbuscular mycorrhizal symbioses, continental regions for the bat pollination dataset (*e.g.* South America, Antilles), and country-level units for the plant dispersal dataset. For species occurring in multiple bioregions, we retained the region with the greatest number of recorded interactions as the primary bioregion. We first fitted phylogenetic generalized least squares (PGLS) models to estimate the contribution of biogeography to each latent trait within each clade, reporting both the explained variance (R^2^) and statistical significance (p-value). Second, for each inferred module, we quantified the number of bioregions represented and calculated, for each clade, the proportion of species pairs belonging to the same bioregion within modules.

## Author Contributions

BPL and HM designed the study. BPL performed the analyses and analyzed the data with help from JA, BM, OPP, and AL. BPL and HM wrote the first version of the manuscript and all authors contributed to the revisions.

## Competing Interest Statement

The authors declare no conflict of interests.

## Acknowledgments

The authors acknowledge G. Dalla Riva for helpful discussions of the RDPG methods as well as the members of the ‘Modeling Biodiversity’ team at IBENS for constructive feedbacks. We also acknowledge the Editor and the three anonymous reviewers for their helpful comments.

This work was performed using HPC resources from GENCI-IDRIS (Grant 2022-AD010313735).

## Data availability

All the functions developed for running ELEFANT are available on GitHub: https://github.com/BPerezLamarque/ELEFANT

All scripts and empirical data used in this manuscript are available on OSF with the following DOI: https://doi.org/10.17605/OSF.IO/K275U

## Notes

### Competing Interest Statement

The authors have declared no competing interest.

### Summary of Updates

We have clarified some aspects and limitations of our approach.

## References

1. J. L. Bronstein, Mutualism, J. L. Bronstein, Ed. (Oxford University Press, 2015) 10.1093/acprof:oso/9780199675654.001.0001 (December 12, 2018).

2. D. H. Hembry, M. G. Weber, Ecological interactions and macroevolution: A new field with old roots. Annu. Rev. Ecol. Evol. Syst. 51, 215–243 (2020).

3. L. J. Harmon, et al., Detecting the macroevolutionary signal of species interactions. J. Evol. Biol. 32, 769–782 (2019).

4. J. Bascompte, P. Jordano, Mutualistic networks (Princeton University Press, 2013).

5. E. Thébault, C. Fontaine, Stability of ecological communities and the architecture of mutualistic and trophic networks. Science (80-). 329, 853–856 (2010).

6. O. Maliet, N. Loeuille, H. Morlon, An individual based model for the eco-evolutionary emergence of bipartite interaction networks. Ecol. Lett. 23, ele.13592 (2020).

7. C. Fontaine, et al., The ecological and evolutionary implications of merging different types of networks. Ecol. Lett. 14, 1170–1181 (2011).

8. M. A. Fortuna, et al., Nestedness versus modularity in ecological networks: Two sides of the same coin? J. Anim. Ecol. 79, 811–817 (2010).

9. C. I. Donatti, et al., Analysis of a hyper-diverse seed dispersal network: Modularity and underlying mechanisms. Ecol. Lett. 14, 773–781 (2011).

10. R. B. P. Pinheiro, G. M. F. Felix, C. F. Dormann, M. A. R. Mello, A new model explaining the origin of different topologies in interaction networks. Ecology 100, e02796 (2019).

11. E. C. Fricke, J. Svenning, Accelerating homogenization of the global plant – frugivore meta-network. Nature 585 (2020).

12. J. Davison, et al., Global assessment of arbuscular mycorrhizal fungus diversity reveals very low endemism. Science (80). 349, 970–973 (2015).

13. B. Perez-Lamarque, M. A. Selosse, M. Öpik, H. Morlon, F. Martos, Cheating in arbuscular mycorrhizal mutualism: a network and phylogenetic analysis of mycoheterotrophy. New Phytol. 226, 1822–1835 (2020).

14. T. Poisot, D. Stouffer, How ecological networks evolve. bioRxiv, 071993 (2016).

15. S. T. Segar, et al., The role of evolution in shaping ecological networks. Trends Ecol. Evol. 35, 454–466 (2020).

16. G. Burin, P. R. Guimarães, T. B. Quental, Macroevolutionary stability predicts interaction patterns of species in seed dispersal networks. Science (80-). 372, 733–737 (2021).

17. J. L. Bronstein, The study of mutualism, past, present, and future. Am. Nat. in press. (2025).

18. L. F. Fuzessy, M. A. Pizo, Navigating a changing world: on the significance of rewiring for mutualistic interactions, caveats and future directions. Oikos 2025, e11230 (2025).

19. M. P. Braga, P. R. Guimarães, C. W. Wheat, S. Nylin, N. Janz, Unifying host-associated diversification processes using butterfly–plant networks. Nat. Commun. 9, 5155 (2018).

20. P. R. Guimarães, P. Jordano, J. N. Thompson, Evolution and coevolution in mutualistic networks. Ecol. Lett. 14, 877–885 (2011).

21. J. M. Olesen, J. Bascompte, Y. L. Dupont, P. Jordano, The modularity of pollination networks. Proc. Natl. Acad. Sci. 104, 19891–19896 (2007).

22. M. L. Stanton, Interacting guilds: Moving beyond the pairwise perspective on mutualisms in American Naturalist, (2003), pp. S10–S23.

23. J. N. Thompson, B. M. Cunningham, Geographic structure and dynamics of coevolutionary selection. Nature 417, 735–738 (2002).

24. J. Genini, L. P. C. Morellato, P. R. Guimarães, J. M. Olesen, Cheaters in mutualism networks. Biol. Lett. 6, 494–497 (2010).

25. J. M. Gómez, M. Verdú, F. Perfectti, Ecological interactions are evolutionarily conserved across the entire tree of life. Nature 465, 918–921 (2010).

26. J. A. Dunne, R. J. Williams, N. D. Martinez, R. A. Wood, D. H. Erwin, Compilation and network analyses of Cambrian food webs. PLoS Biol. 6, 693– 708 (2008).

27. J. C. S. Nascimento, F. Blanco, M. S. Domingo, J. L. Cantalapiedra, M. M. Pires, The reorganization of predator–prey networks over 20 million years explains extinction patterns of mammalian carnivores. Ecol. Lett. 27, e14448 (2024).

28. M. J. Landis, N. J. Matzke, B. R. Moore, J. P. Huelsenbeck, Bayesian analysis of biogeography when the number of areas is large. Syst. Biol. 62, 789–804 (2013).

29. M. P. Braga, M. J. Landis, S. Nylin, N. Janz, F. Ronquist, Bayesian inference of ancestral host-parasite interactions under a phylogenetic model of host repertoire evolution. Syst. Biol. 69, 1149–1162 (2020).

30. M. P. Braga, N. Janz, S. Nylin, F. Ronquist, M. J. Landis, Phylogenetic reconstruction of ancestral ecological networks through time for pierid butterflies and their host plants. Ecol. Lett. 24, 2134–2145 (2021).

31. G. V. Dalla Riva, D. B. Stouffer, Exploring the evolutionary signature of food webs’ backbones using functional traits. Oikos 125, 446–456 (2016).

32. T. Strydom, et al., Food web reconstruction through phylogenetic transfer of low-rank network representation. Methods Ecol. Evol. 2022, 1–12 (2022).

33. M.-A. Selosse, F. Le Tacon, The land flora: a phototroph-fungus partnership? Trends Ecol. Evol. 13, 15–20 (1998).

34. Y. Zeng, J. J. Wiens, Do mutualistic interactions last longer than antagonistic interactions? Proc. R. Soc. B Biol. Sci. 288 (2021).

35. D. Peris, et al., Evolutionary implications of a deep-time perspective on insect pollination. Biol. Rev. 100, 1452–1466 (2025).

36. A. Nunes Martinez, M. Mistretta Pires, Estimated missing interactions change the structure and alter species roles in one of the world’s largest seed-dispersal networks. Oikos 2024, e10521 (2024).

37. S. L. Nuismer, L. J. Harmon, Predicting rates of interspecific interaction from phylogenetic trees. Ecol. Lett. 18, 17–27 (2015).

38. H. Morlon, et al., Phylogenetic insights into diversification. Annu. Rev. Ecol. Evol. Syst. (2024) 10.1146/annurev-ecolsys-102722-020508 (October 26, 2024).

39. O. Maliet, H. Morlon, Fast and accurate estimation of species-specific diversification rates using data augmentation. Syst. Biol. 71, 353–366 (2022).

40. F. Massol, E. Macke, M. Callens, E. Decaestecker, A methodological framework to analyse determinants of host–microbiota networks, with an application to the relationships between Daphnia magna’s gut microbiota and bacterioplankton. J. Anim. Ecol. 90, 102–119 (2021).

41. W. R. Rimington, S. Pressel, J. G. Duckett, K. J. Field, M. I. Bidartondo, Evolution and networks in ancient and widespread symbioses between Mucoromycotina and liverworts. Mycorrhiza 29, 551–565 (2019).

42. B. Perez-Lamarque, et al., Analysing diversification dynamics using barcoding data: The case of an obligate mycorrhizal symbiont. Mol. Ecol. 31, 3496–3512 (2022).

43. F. Lutzoni, et al., Contemporaneous radiations of fungi and plants linked to symbiosis. Nat. Commun. 9, 1–11 (2018).

44. B. Pichon, R. Le Goff, H. Morlon, B. Perez-Lamarque, Telling mutualistic and antagonistic ecological networks apart by learning their multiscale structure. Methods Ecol. Evol. 15, 1113–1128 (2024).

45. L. Fuzessy, M. Verdú, M. A. Pizo, Ecological function over evolutionary legacy: The limited role of shared evolutionary history in shaping modern frugivory interactions. Funct. Ecol. 00, 3688–3703 (2025).

46. E. L. Rezende, P. Jordano, J. Bascompte, Effects of phenotypic complementarity and phylogeny on the nested structure of mutualistic networks. Oikos 116, 1919–1929 (2007).

47. A. R. Ives, H. C. J. Godfray, Phylogenetic analysis of trophic associations. Am. Nat. 168, E1–E14 (2006).

48. B. Perez-Lamarque, et al., Do closely related species interact with similar partners? Testing for phylogenetic signal in bipartite interaction networks. Peer Community J. 2, e59 (2022).

49. N. Blüthgen, M. Staab, A critical evaluation of network approaches for studying species interactions. Annu. Rev. Ecol. Evol. Syst. 55, 65–88 (2024).

50. B. Perez-Lamarque, H. Morlon, Distinguishing cophylogenetic signal from phylogenetic congruence clarifies the interplay between evolutionary history and species interactions. Syst. Biol. 73, 613–622 (2024).

51. F. Encinas-Viso, T. A. Revilla, R. S. Etienne, Phenology drives mutualistic network structure and diversity. Ecol. Lett. 15, 198–208 (2012).

52. M. A. R. Mello, et al., Insights into the assembly rules of a continent-wide multilayer network. *Nat*. Ecol. Evol. 3, 1525–1532 (2019).

53. L. J. Revell, L. J. Harmon, D. C. Collar, Phylogenetic signal, evolutionary process, and rate. Syst. Biol. 57, 591–601 (2008).

54. M. Manceau, A. Lambert, H. Morlon, A unifying comparative phylogenetic framework including traits coevolving across interacting lineages. Syst. Biol. 66, 551–568 (2017).

55. C. B. Fenster, W. S. Armbruster, P. Wilson, M. R. Dudash, J. D. Thomson, Pollination syndromes and floral specialization. Annu. Rev. Ecol. Evol. Syst. 35, 375–403 (2004).

56. J. P. Drury, et al., Contrasting impacts of competition on ecological and social trait evolution in songbirds. PLoS Biol. 16, e2003563 (2018).

57. C. Llopis-Belenguer, et al., Sensitivity of bipartite network analyses to incomplete sampling and taxonomic uncertainty. Ecology 104, e3974 (2023).

58. I. Quintero, N. Lartillot, H. Morlon, Imbalanced speciation pulses sustain the radiation of mammals. Science (80). 384, 1007–1012 (2024).

59. N. Chabrol, H. Morlon, J. Barido-Sottani, The Fossilized Birth Death Process with heterogeneous diversification rates unravels the link between diversification and specialisation to a carnivorous diet in Nimravidae (Carnivoraformes). bioRxiv, 2025.07.15.664897 (2025).

60. I. Quintero, J. Andréoletti, D. Silvestro, H. Morlon, Loss of macroevolutionary species fitness explains the rise and fall of clades. *Nat*. Ecol. Evol. 9, 2346–2357 (2025).

61. P. R. Guimarães, M. Galetti, P. Jordano, Seed dispersal anachronisms: Rethinking the fruits extinct megafauna ate. PLoS One 3, e1745 (2008).

62. E. E. Goldberg, L. T. Lancaster, R. H. Ree, Phylogenetic inference of reciprocal effects between geographic range evolution and diversification. Syst. Biol. 60, 451–465 (2011).

63. I. Quintero, M. J. Landis, W. Jetz, H. Morlon, The build-up of the present-day tropical diversity of tetrapods. Proc. Natl. Acad. Sci. 120 (2023).

64. T. Poisot, D. B. Stouffer, D. Gravel, Beyond species: Why ecological interaction networks vary through space and time. Oikos 124, 243–251 (2015).

65. E. L. Rezende, J. E. Lavabre, P. R. Guimarães, P. Jordano, J. Bascompte, Non-random coextinctions in phylogenetically structured mutualistic networks. Nature 448, 925–928 (2007).

66. I. Junk, et al., Archived natural DNA samplers reveal four decades of biodiversity change across the tree of life. Nat. Ecol. Evol., 1–12 (2025).

67. T. Poisot, et al., Network embedding unveils the hidden interactions in the mammalian virome. Patterns 4, 100738 (2023).

68. J. Clavel, G. Escarguel, G. Merceron, mvMORPH: An R package for fitting multivariate evolutionary models to morphometric data. Methods Ecol. Evol. 6, 1311–1319 (2015).

69. E. P. Martins, T. F. Hansen, Phylogenies and the comparative method: A general approach to incorporating phylogenetic information into the analysis of interspecific data. Am. Nat. 149, 646–667 (1997).

70. J. Clavel, L. Aristide, H. Morlon, A penalized likelihood framework for high-dimensional phylogenetic comparative methods and an application to New-World monkeys brain evolution. Syst. Biol. 68, 93–116 (2019).

71. C. F. Dormann, B. Gruber, J. Fründ, Introducing the bipartite package: analysing ecological networks. R News 8, 8–11 (2008).

72. D. M. de Vienne, et al., Cospeciation vs host-shift speciation: Methods for testing, evidence from natural associations and relation to coevolution. New Phytol. 198, 347–385 (2013).

73. N. E. Langmore, et al., Coevolution with hosts underpins speciation in brood-parasitic cuckoos. Science (80). 384, 1030–1036 (2024).

74. R Core Team, R: A language and environment for statistical computing. (2024).

75. L. L. Sánchez-Reyes, H. Morlon, S. Magallón, Uncovering higher-taxon diversification dynamics from clade age and species-richness data. Syst. Biol. 66, 367–378 (2017).

76. J. Nagy, Z. Végvári, Z. Varga, Phylogeny, migration and life history: filling the gaps in the origin and biogeography of the Turdus thrushes. J. Ornithol. 160, 529–543 (2019).

77. M. Öpik, et al., The online database MaarjAM reveals global and ecosystemic distribution patterns in arbuscular mycorrhizal fungi (Glomeromycota). New Phytol. 188, 223–241 (2010).

78. Y. Jin, H. Qian, V.PhyloMaker2: An updated and enlarged R package that can generate very large phylogenies for vascular plants. Plant Divers. 44, 335–339 (2022).

79. H. Morlon, et al., RPANDA: An R package for macroevolutionary analyses on phylogenetic trees. Methods Ecol. Evol. 7, 589–597 (2016).

80. K. González-Gutiérrez, J. H. Castaño, J. Pérez-Torres, H. R. Mosquera-Mosquera, Structure and roles in pollination networks between phyllostomid bats and flowers: a systematic review for the Americas. Mamm. Biol. 102, 21–49 (2022).

81. N. S. Upham, J. A. Esselstyn, W. Jetz, Inferring the mammal tree: Species-level sets of phylogenies for questions in ecology, evolution, and conservation. PLoS Biol. 17, e3000494 (2019).

82. T. Datzmann, O. Von Helversen, F. Mayer, Evolution of nectarivory in phyllostomid bats (Phyllostomidae Gray, 1825, Chiroptera: Mammalia). BMC Evol. Biol. 10, 1–14 (2010).

83. T. H. Fleming, C. Geiselman, W. J. Kress, The evolution of bat pollination: a phylogenetic perspective. Ann. Bot. 104, 1017–43 (2009).

84. W. Jetz, G. H. Thomas, J. B. Joy, K. Hartmann, A. O. Mooers, The global diversity of birds in space and time. Nature 491, 444–448 (2012).

85. W. D. Kissling, K. Böhning-Gaese, W. Jetz, The global distribution of frugivory in birds. Glob. Ecol. Biogeogr. 18, 150–162 (2009).

86. P. Vargas, R. Heleno, J. M. Costa, EuDiS - A comprehensive database of the seed dispersal syndromes of the European flora. Biodivers. data J. 11, e104079 (2023).

87. O. Maliet, F. Hartig, H. Morlon, A model with many small shifts for estimating species-specific diversification rates. Nat. Ecol. Evol. 3, 1086–1092 (2019).

