## Supplementary material for "Structural stability of mutualistic networks over large geographic and temporal scales": Sup Fig

**Supplementary Methods**
**Supplementary Results**
**Supplementary References**
**Supplementary Figures**
**Supplementary Tables**

**Supplementary methods:**

**Data augmentation of extinct and unsampled lineages on the plant and mutualist phylogenetic trees assuming a constant-rate birth-death process:**

For each guild, we considered that the reconstructed phylogenetic tree is the result of a constant-rate, homogeneous birth-death process with a speciation rate $\lambda$, an extinction rate $\mu$, and a sampling probability $\rho$ at present (t=0).

Let $u(t)$ be the probability that a lineage alive at time t has no representative lineage in the reconstructed tree. In particular, at present, we have $u(0) = 1 - \rho$ and in general, $u(t)$ is given by the following formula (Morlon *et al.* 2011):

$$u(t) = 1 - \frac{e^{(\lambda-\mu)t}}{\frac{1}{\rho} + \frac{\lambda}{\lambda - \mu}\left(e^{(\lambda-\mu)t} - 1\right)}$$

Given the reconstructed tree, the sub-trees that are not observed (either because they are unsampled at present or went extinct) have the following properties:

- they are independent;
- they are branched on the reconstructed trees according to an inhomogeneous Poisson point process with rate $2\lambda u(t)$;
- they follow an inhomogeneous birth-death process with a speciation rate of $\lambda u(t)$ and an extinction rate of $\mu/u(t)$.

If $\rho$=1 (all extant lineages are sampled), the sub-trees cannot reach the present (t = 0) because the extinction rate $\mu/u(t)$ tends toward infinity when t tends toward 0.

If $\rho$ is lower than 1, the sub-trees can reach 0 and the extant species are not sampled in the reconstructed tree.

Following this approach, we implemented the data augmentation of unsampled and extinct lineages on the reconstructed trees in R (R Core Team 2024) using the R-packages ape (Paradis *et al.* 2004) and phytools (Revell 2012). To simulate inhomogeneous Poisson processes, we employed the thinning method (Lewis & Shedler 1979). See Supplementary Figure S16 for examples of data augmented trees.

All the functions developed for this data augmentation step are available on GitHub: https://github.com/BPerezLamarque/ELEFANT

The same GitHub link contain a tutorial for running ELEFANT. Depending on the network size and the parameters of the data augmentation step ($\lambda$, $\mu$, $\rho$, and the number of replicates), the inferences can take from one hour to several days, with the majority of the computational time being spent calculating the global metrics of the ancestral networks.

**Supplementary Results:**

***Properties of the simulated networks:***

We verified that our simulations generated expected networks structures, in particular in term of phylogenetic signals (Fig. S11). Significant and high phylogenetic signals in both clades were correctly generated under simulations of network evolution by anagenetic changes and evolution of latent traits by Brownian motions. As expected, in simulations of host repertoire evolution, only phylogenetic signals in the symbionts were present (*i.e.* closely related symbionts interact with similar hosts) but not in the hosts. We only detected phylogenetic signals in the hosts when simulating network evolution by cladogenetic shifts but not in the symbionts, as one of the daughter symbiont species loses all its interactions at symbiont speciation. Finally, no phylogenetic signal was detected in simulations of random evolution. ELEFANT is unlikely to provide correct ancestral network estimations in these two last simulated systems presenting no or low phylogenetic signal on both sides of the network as it indicates that interspecific interactions were weakly evolutionarily conserved.

Using the random-dot product graph method (RDPG), between 5 and 15 singular values were often necessary to explain at least 90% of the present-day interactions in the different types of simulated networks (Fig. S12). Therefore, less than 15 latent traits ($d$) were needed to describe the simulated networks, except in the simulations of network evolution by cladogenetic shifts or random evolution, which often required $d$ > 15 latent traits (Fig. S12). A large number of latent traits is indicative of the high dimensionality of the present-day interaction network.

***Performance of ELEFANT:***

Although the estimations of global metrics may be more noisy back in the past (20 Myr ago), ELEFANT successfully recovered the temporal trends in the global metrics (Fig. S7): in the simulated scenarios i), iii), and iv), we correctly estimated a decrease (resp. increase) through time of the connectance in 91% (resp. 75%) of the cases, of the nestedness in 84% (resp. 81%) of the cases, and of the modularity in 84% (resp. 71%) of the cases (Fig. S7).

Adding extinct and unsampled lineages using data augmentation (DA) significantly increased the accuracy of the estimations of the ancestral numbers of species (always underestimated otherwise), the connectance (with DA: $R^2$=0.66; without DA: $R^2$=0.60), nestedness (with DA: $R^2$=0.50; without DA: $R^2$=0.41), and modularity (with DA: $R^2$=0.55; without DA: $R^2$=0.48; Fig. S13).

Regarding sensitivity to the threshold used to define interactions, both global network metrics and ancestral interaction accuracy remain consistently high across a broad intermediate range of thresholds (0.3-0.6; Fig. S14). Performance peaks around a threshold of 0.5, and thresholds selected using Youden's J fall within this same high-performing range. These analyses demonstrate that ELEFANT's results are robust to the choice of threshold and that Youden's J provides a suitable data-driven criterion for threshold selection.

**Supplementary references:**


Lewis, P.W. & Shedler, G.S. (1979). Simulation of nonhomogeneous Poisson processes by thinning. *Nav. Res. Logist. Q.*, 26, 403–413.

Morlon, H., Parsons, T.L. & Plotkin, J.B. (2011). Reconciling molecular phylogenies with the fossil record. *Proc. Natl. Acad. Sci.*, 108, 16327–16332.

Paradis, E., Claude, J. & Strimmer, K. (2004). APE: Analyses of phylogenetics and evolution in R language. *Bioinformatics*, 20, 289–290.

R Core Team. (2024). R: A language and environment for statistical computing.

Revell, L.J. (2012). phytools: An R package for phylogenetic comparative biology (and other things). *Methods Ecol. Evol.*, 3, 217–223.

**Supplementary Figures:**

**Supplementary Figure 1: Description of the cross-validation procedure of ELEFANT**

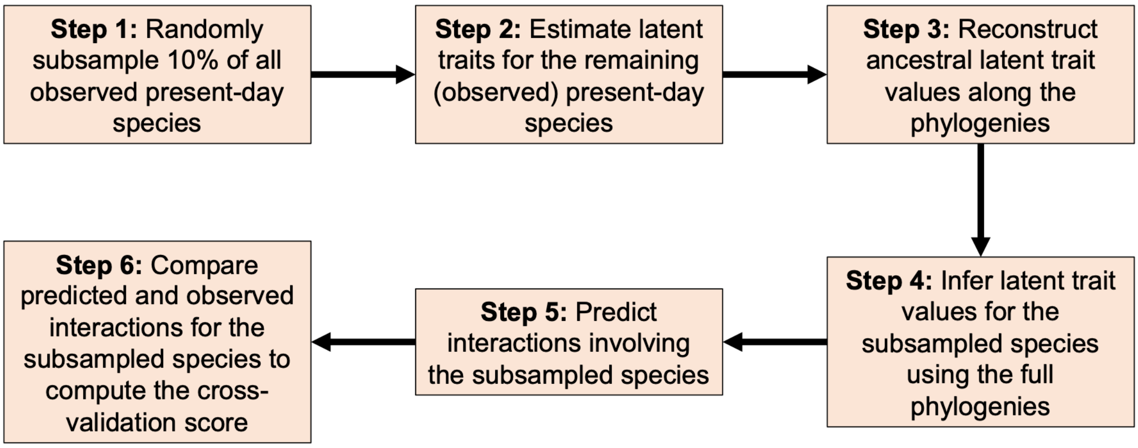

**Supplementary Figure 2: Results of the cross-validations performed on the different types of simulations.**

During cross-validations, we randomly removed 10% of the species of each guild and predicted their interactions based on ELEFANT: we reported the average cross-validation score (A), as well as the mean probabilities of true positives (B) and false positives (C).

**a) Network evolution by anagenetic shifts**

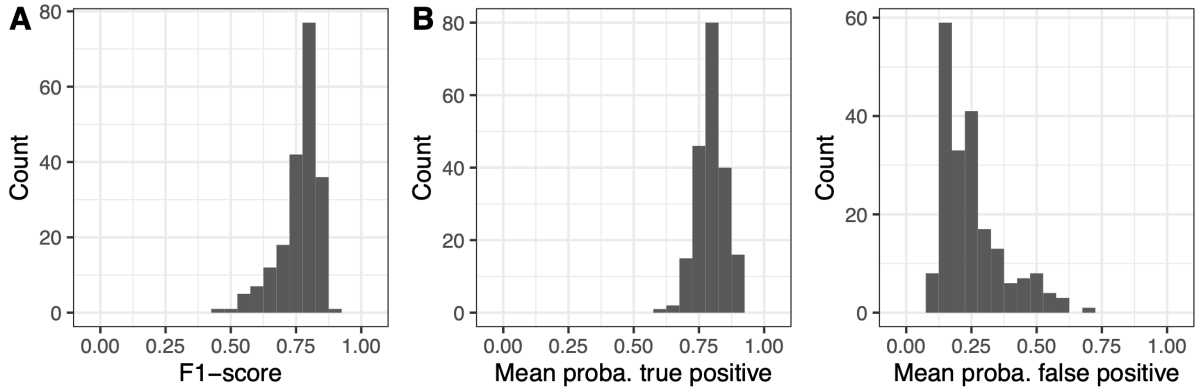


**b) Network evolution by cladogenetic shifts**

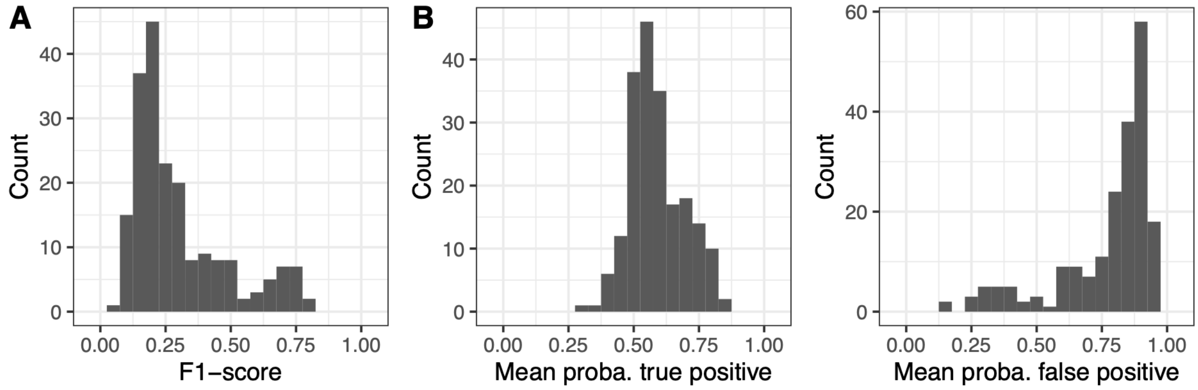


**c) Evolution of latent traits by Brownian motions**

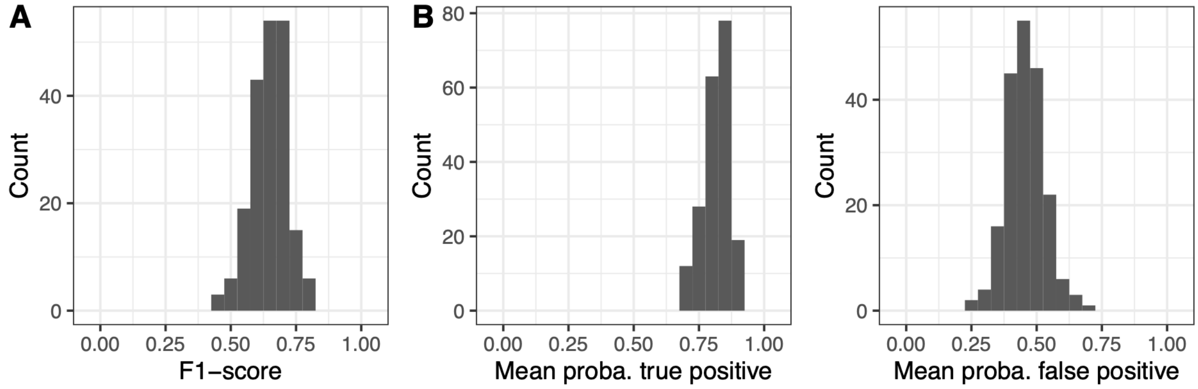

**d) Host repertoire evolution**

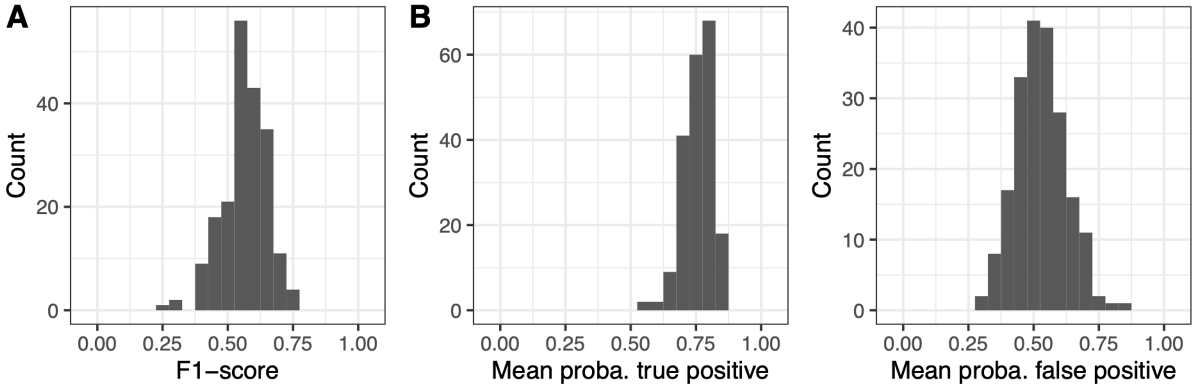


**e) Random evolution**

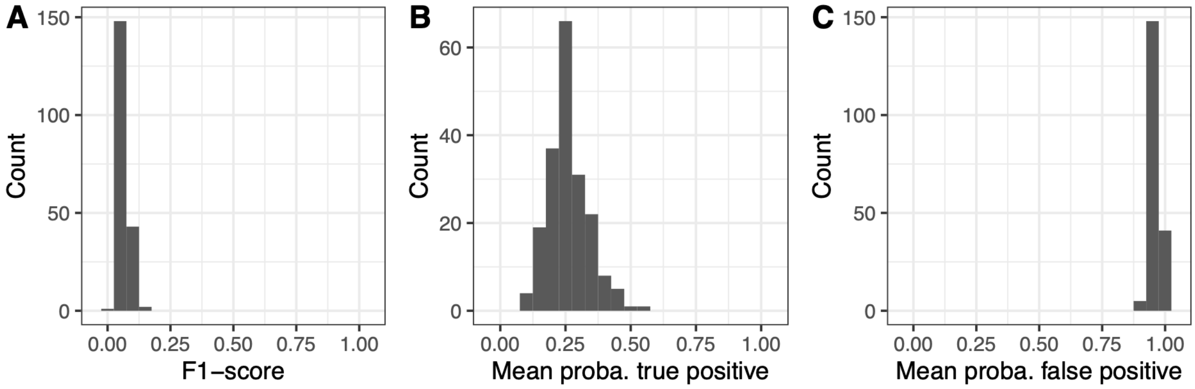

**Supplementary Figure 3: The ability to recover the simulated ancestral interactions using ELEFANT depends on the cross-validation scores:**

The plots indicate the F1-scores of the inferred ancestral interactions at a given time in the past (from 5 to 30 Myr ago) as a function of the cross-validation scores (F1-scores of the cross-validations at present).

F1-scores of the inferred ancestral interactions were computed from the consensus networks assembled from interactions that are recovered in at least J% of the augmented ancestral networks.

Each dot corresponds to one simulation and the dot color represents the cross-validation scores. The black lines represent the line y=x, while the blue lines correspond to the fit of a linear model.

The orange line (x = 0.25) indicates the cross-validation score below which predicted ancestral interactions are considered unreliable.

The last plot (f) summarizes all the results obtained across different simulations.

### a) Network evolution by anagenetic shifts

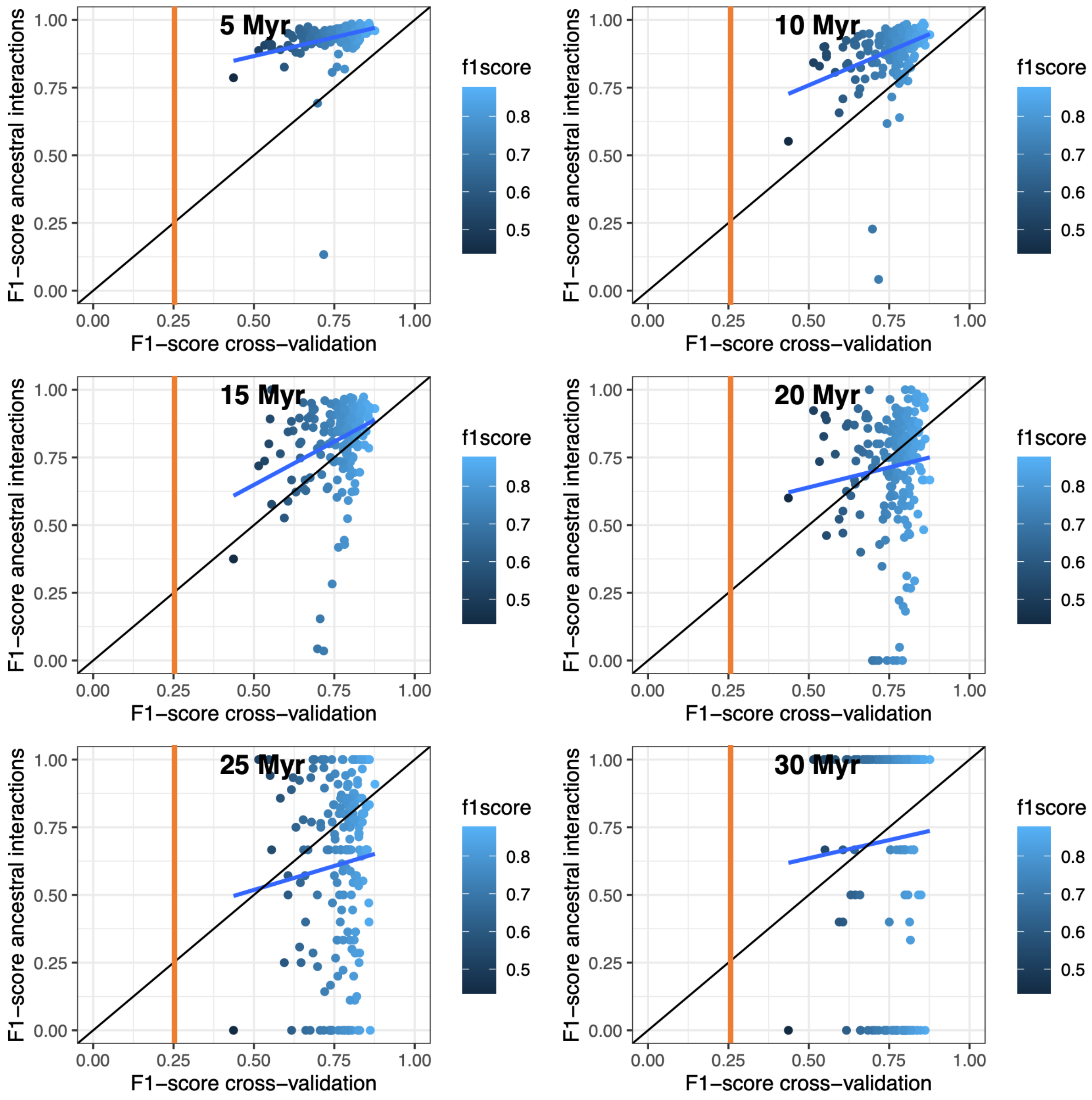

**b) Network evolution by cladogenetic shifts**

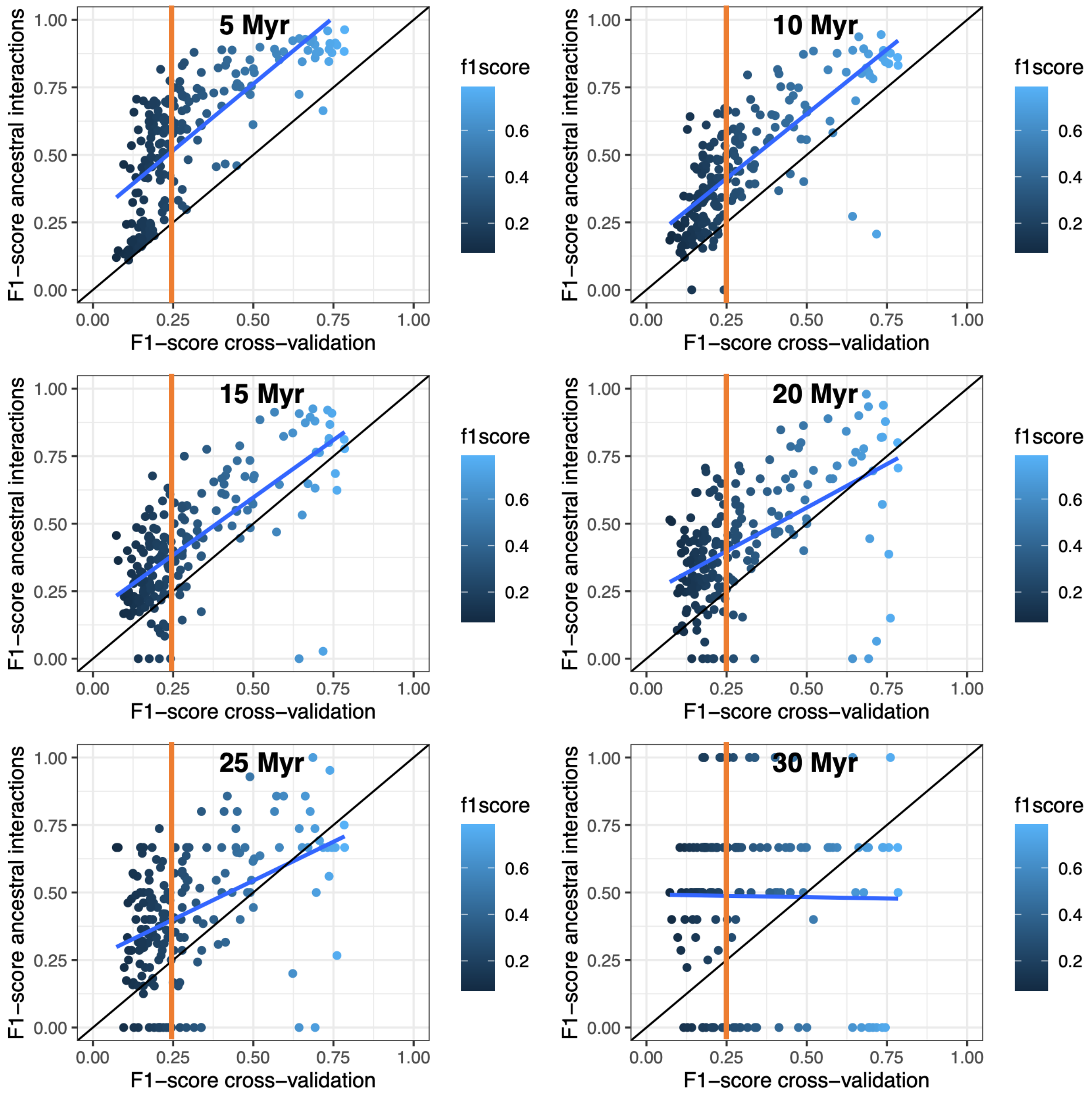

#### c) Evolution of latent traits by Brownian motions

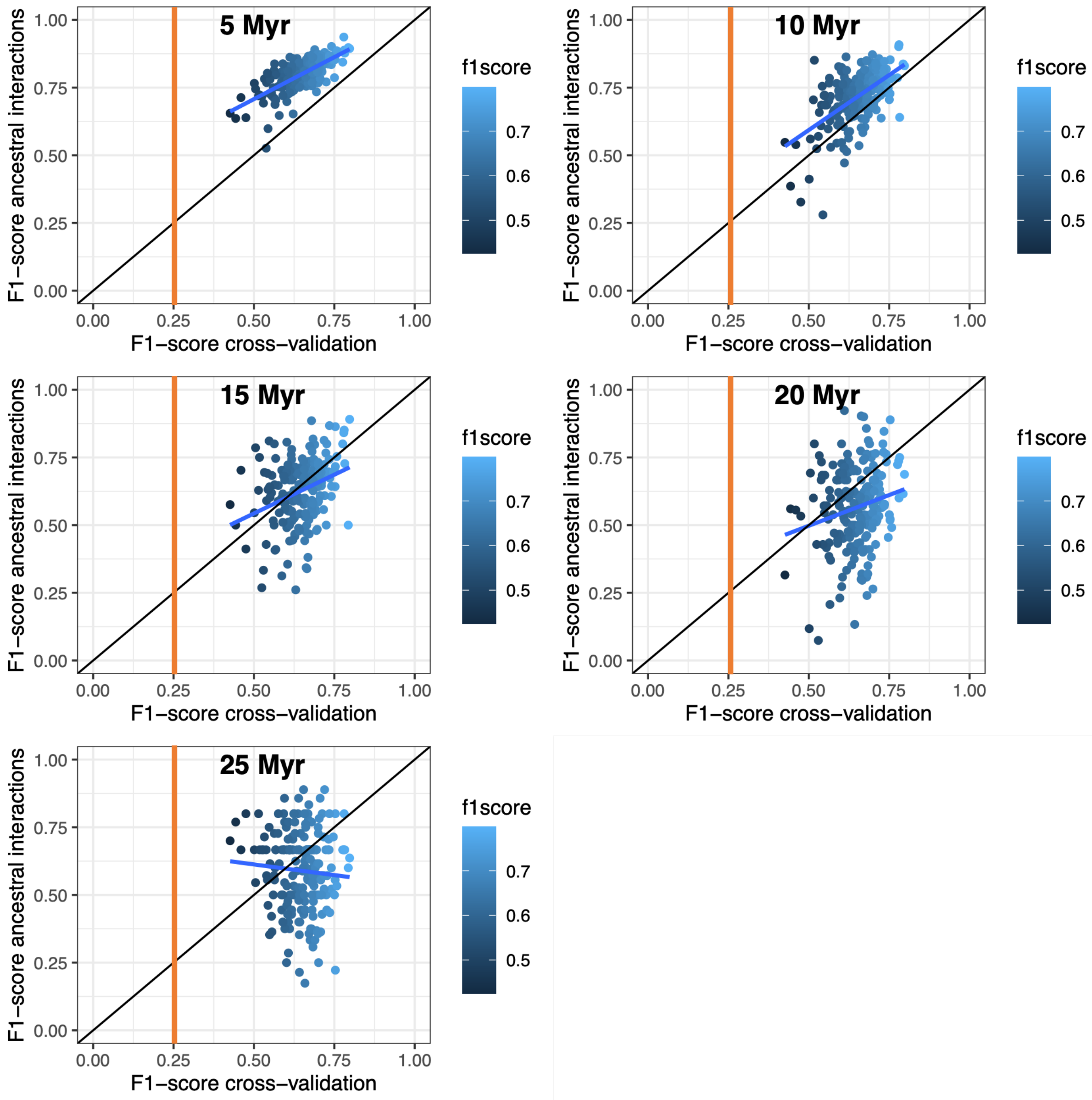

**d) Host repertoire evolution**

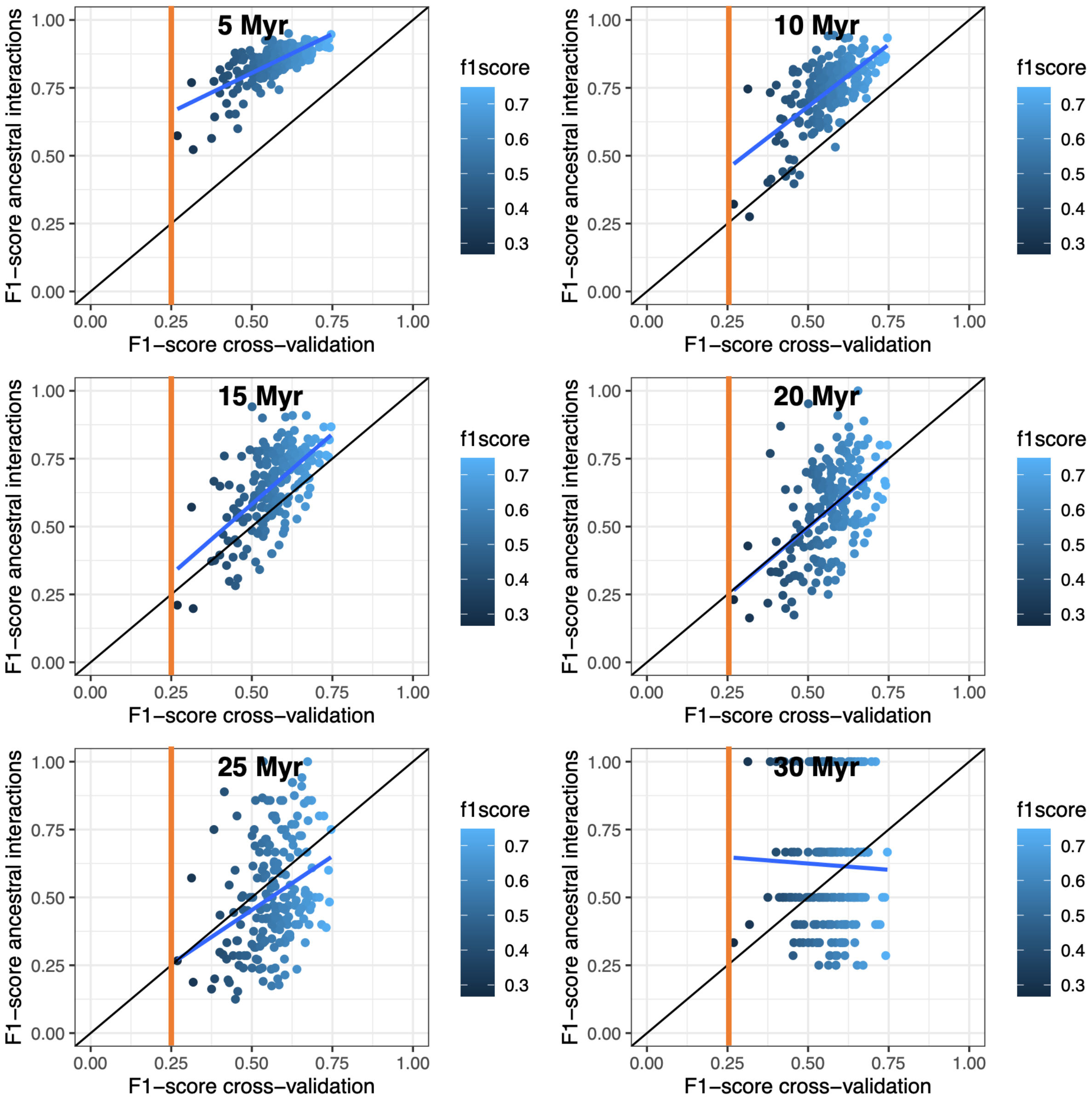

### e) Random evolution

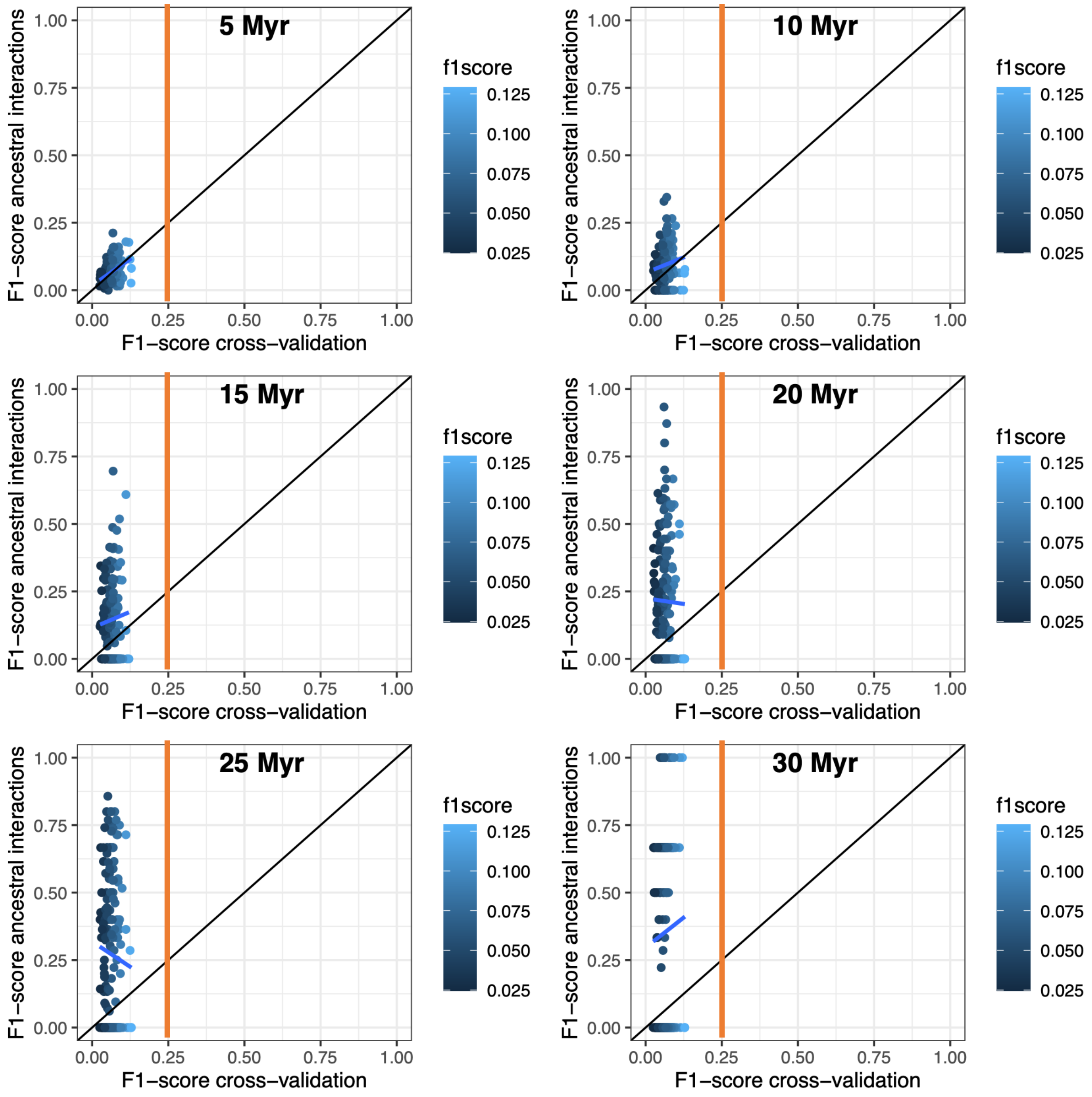

**(f) All simulations:**

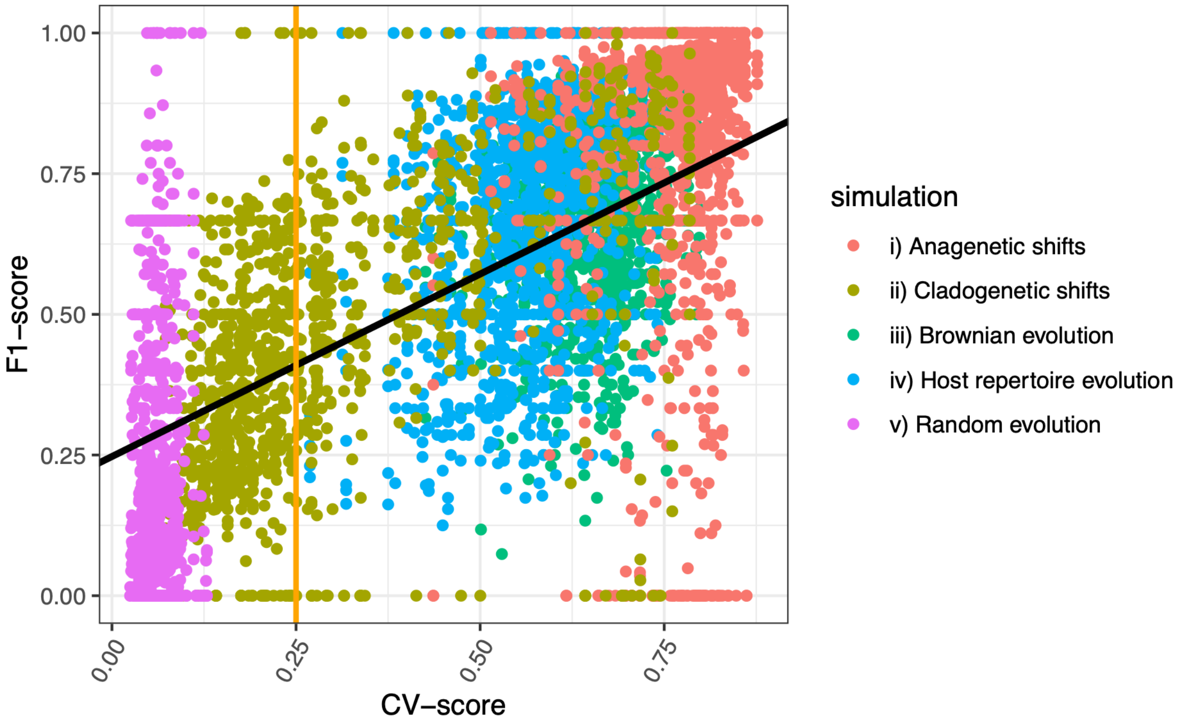


**All simulations but only looking at the recent past (less than 15 Myr in the simulations) :**

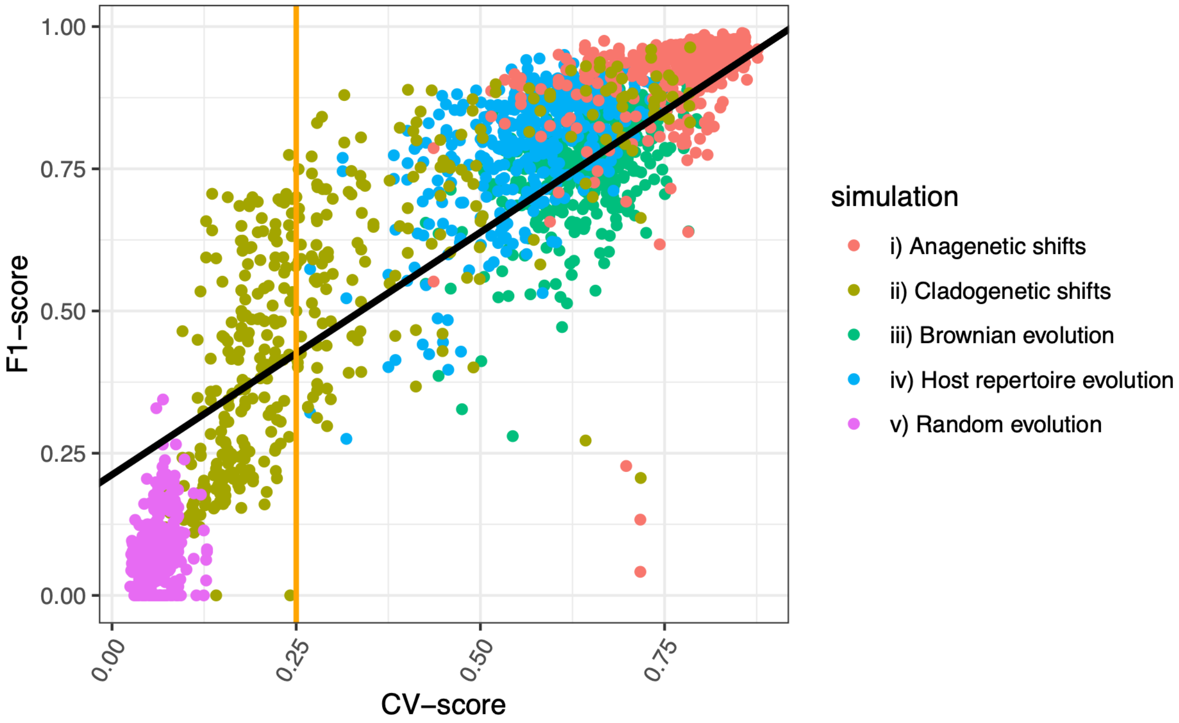

**Supplementary Figure 4: Ability to recover the simulated ancestral interactions for the different types of simulations using ELEFANT.**

The bar charts indicate the F1-score of the ancestral interactions (left panels), the percentage of recovered interactions (middle panels), and the percentage of false positives (right panels) obtained when estimating ancestral interactions in the past (from 5 to 30 Myr ago).

We considered here the 'consensus networks' assembled from interactions that are recovered in at least X% of the 'augmented ancestral networks' with X being 10% (orange), 25% (green), 50% (blue), or the threshold obtained maximizing the Youden's J statistic (grey).

### a) Network evolution by anagenetic shifts

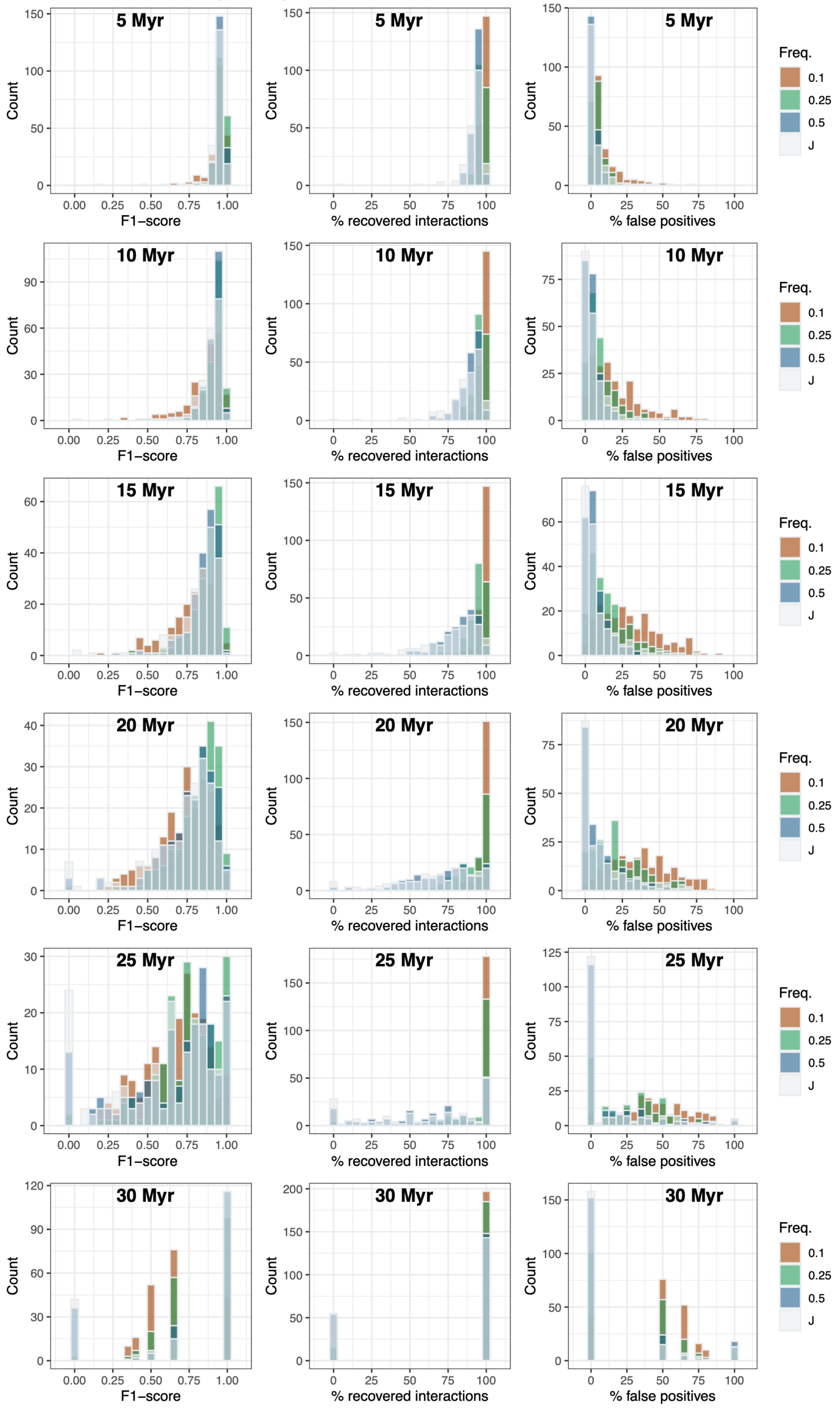

### b) Network evolution by cladogenetic shifts

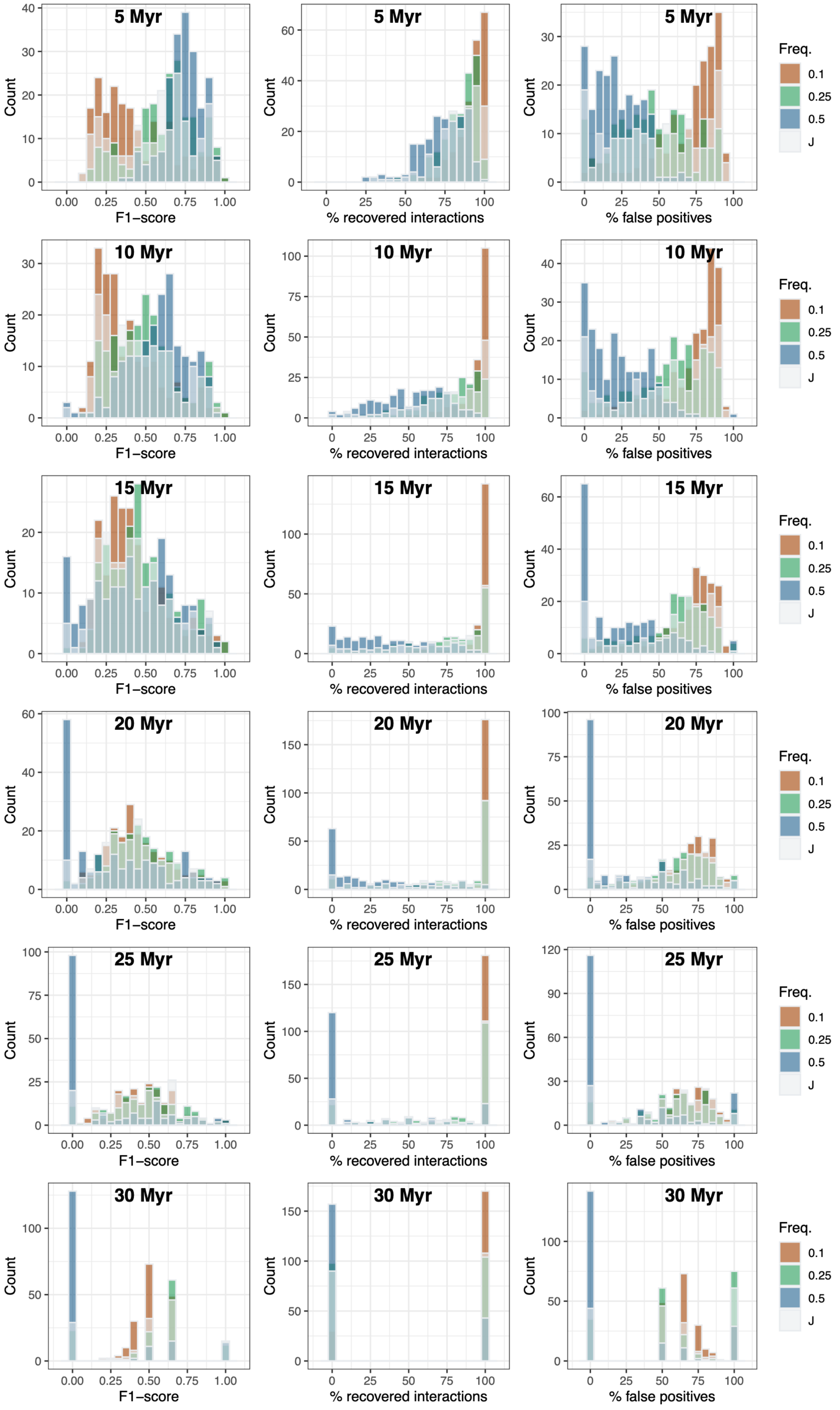

### c) Evolution of latent traits by Brownian motions

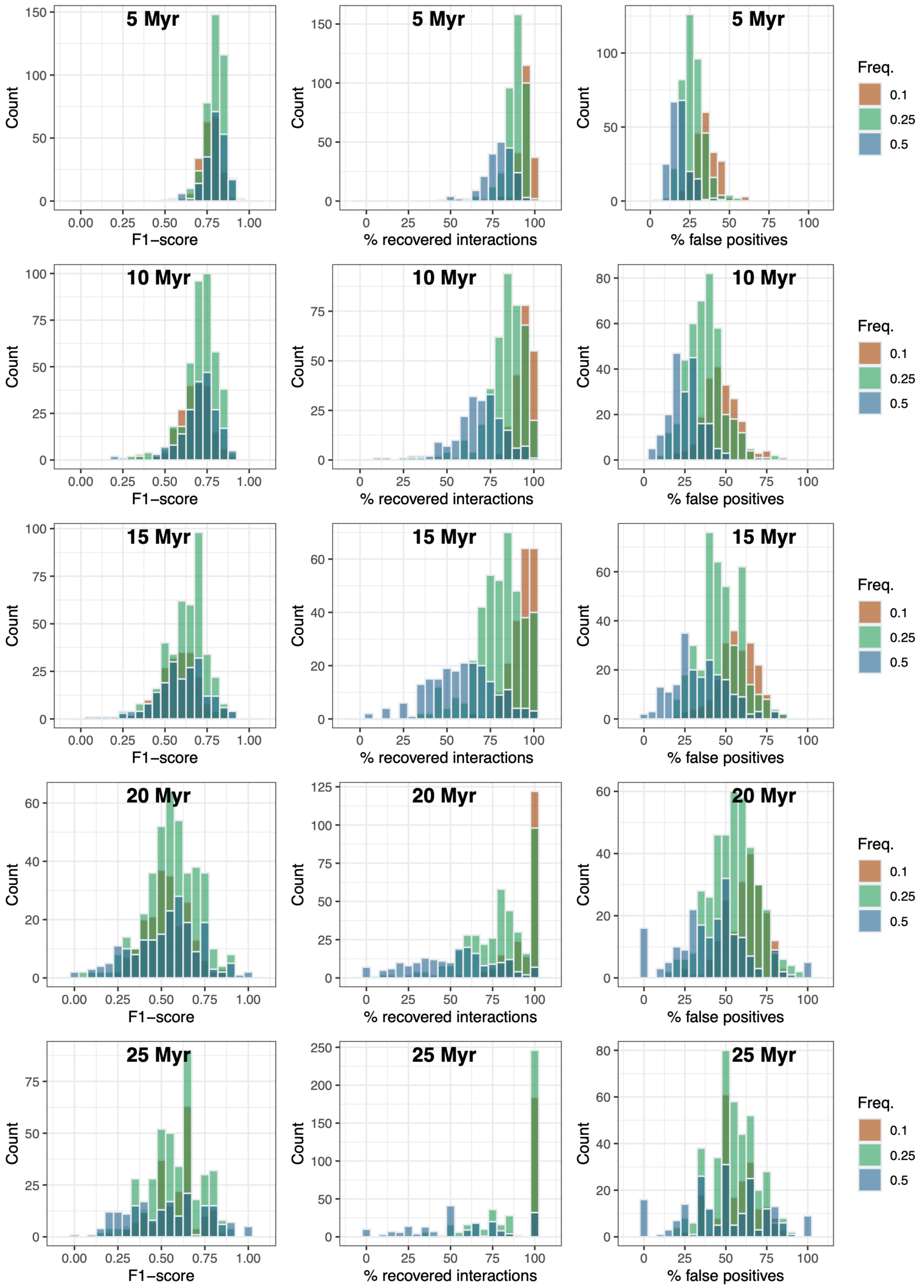

### d) Host repertoire evolution

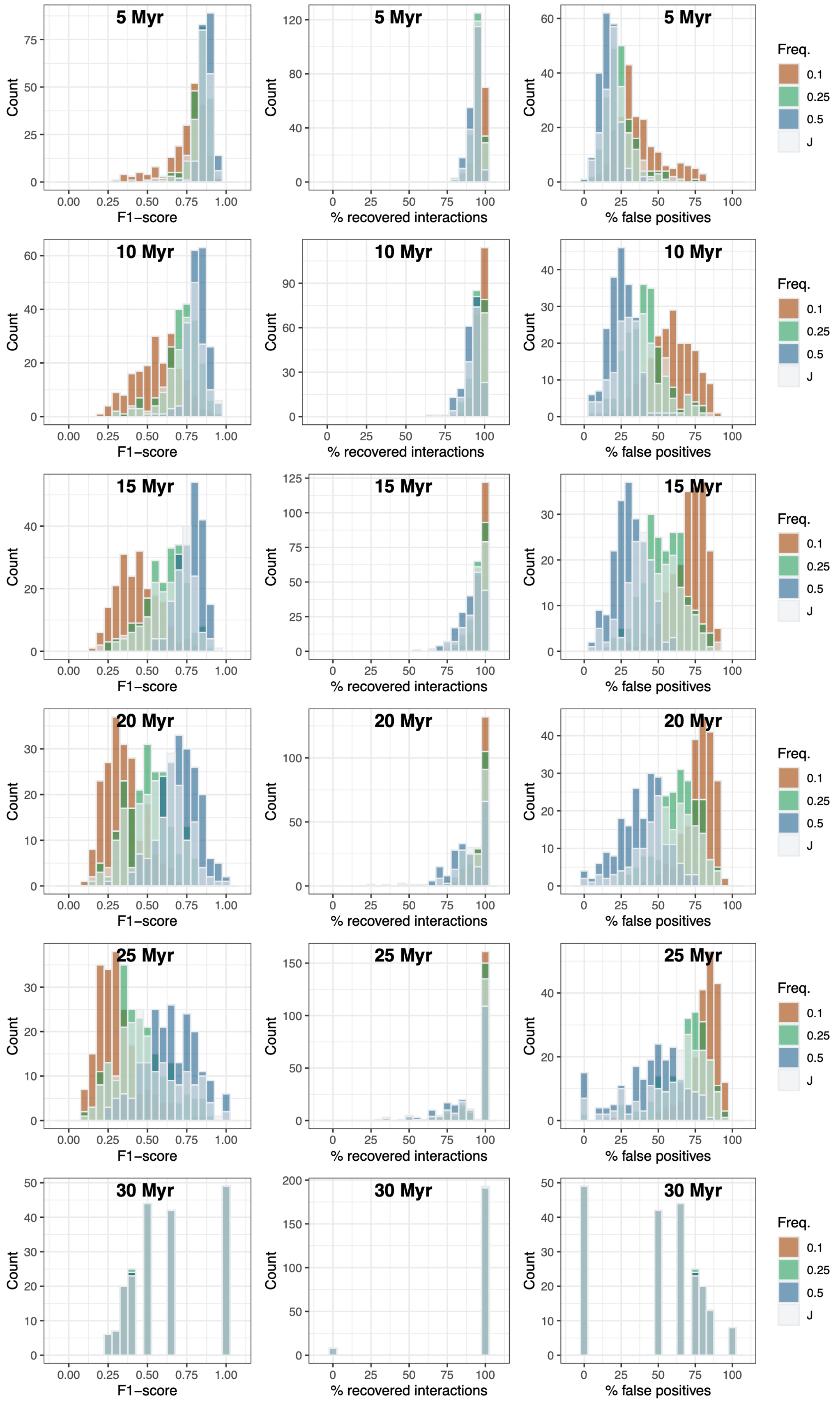

### e) Random evolution

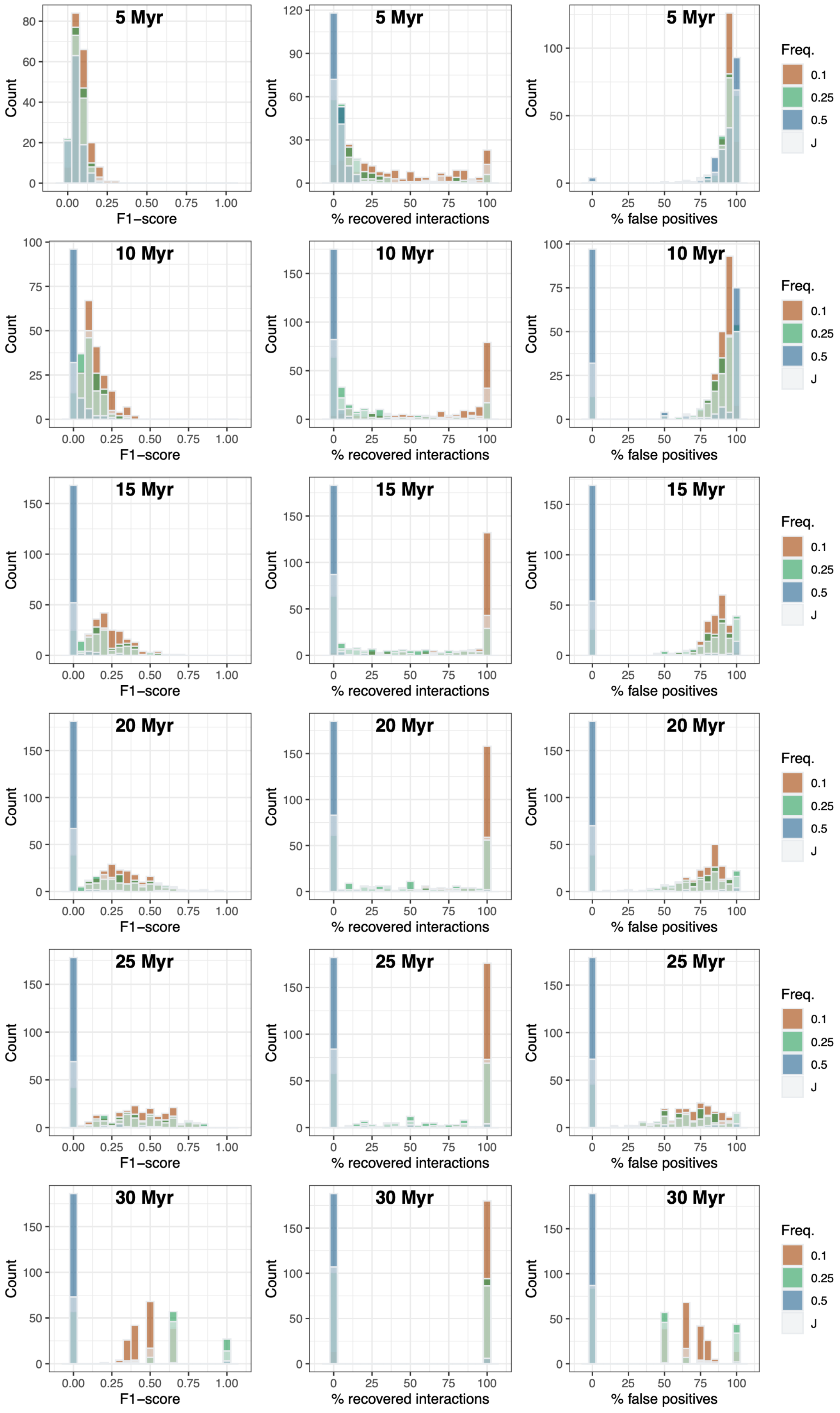

**Supplementary Figure 5: Estimations of the global metrics of the ancestral networks as a function of the simulated ones for the different types of simulations using ELEFANT.**

The estimations of the ancestral connectance (left panels), nestedness (middle panels), and modularity (right panels) are represented at different times in the past (from 5 to 20 Myr ago).

Each dot corresponds to one simulation, and for each simulation, we reported the estimated values averaged over 100 'augmented' ancestral networks.

The dot color represents the cross-validation scores. The black lines represent the line y=x, while the blue lines correspond to the fit of a linear model between the estimated and simulated values.

### a) Network evolution by anagenetic shifts

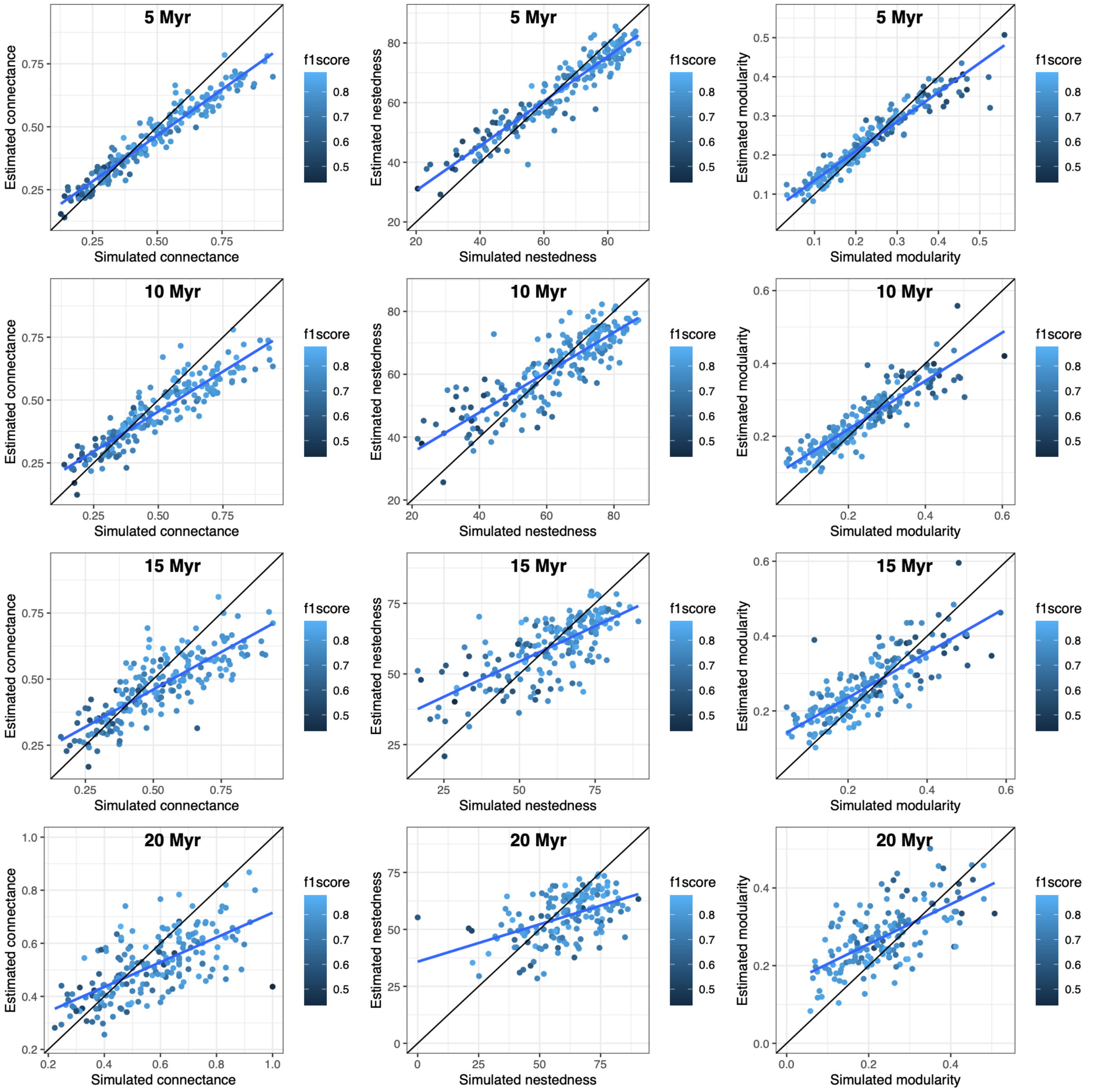

### b) Network evolution by cladogenetic shifts

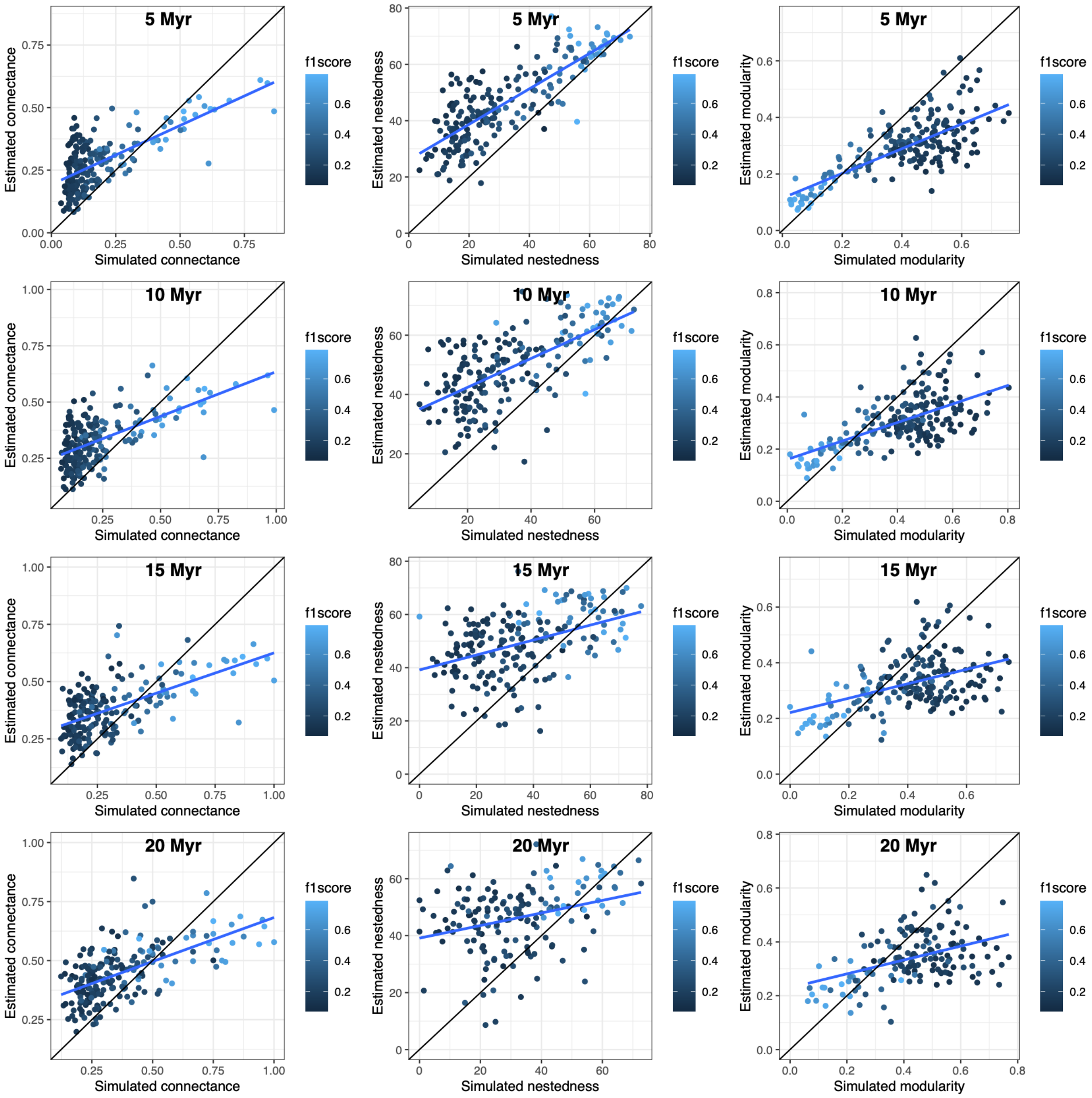

### c) Evolution of latent traits by Brownian motions

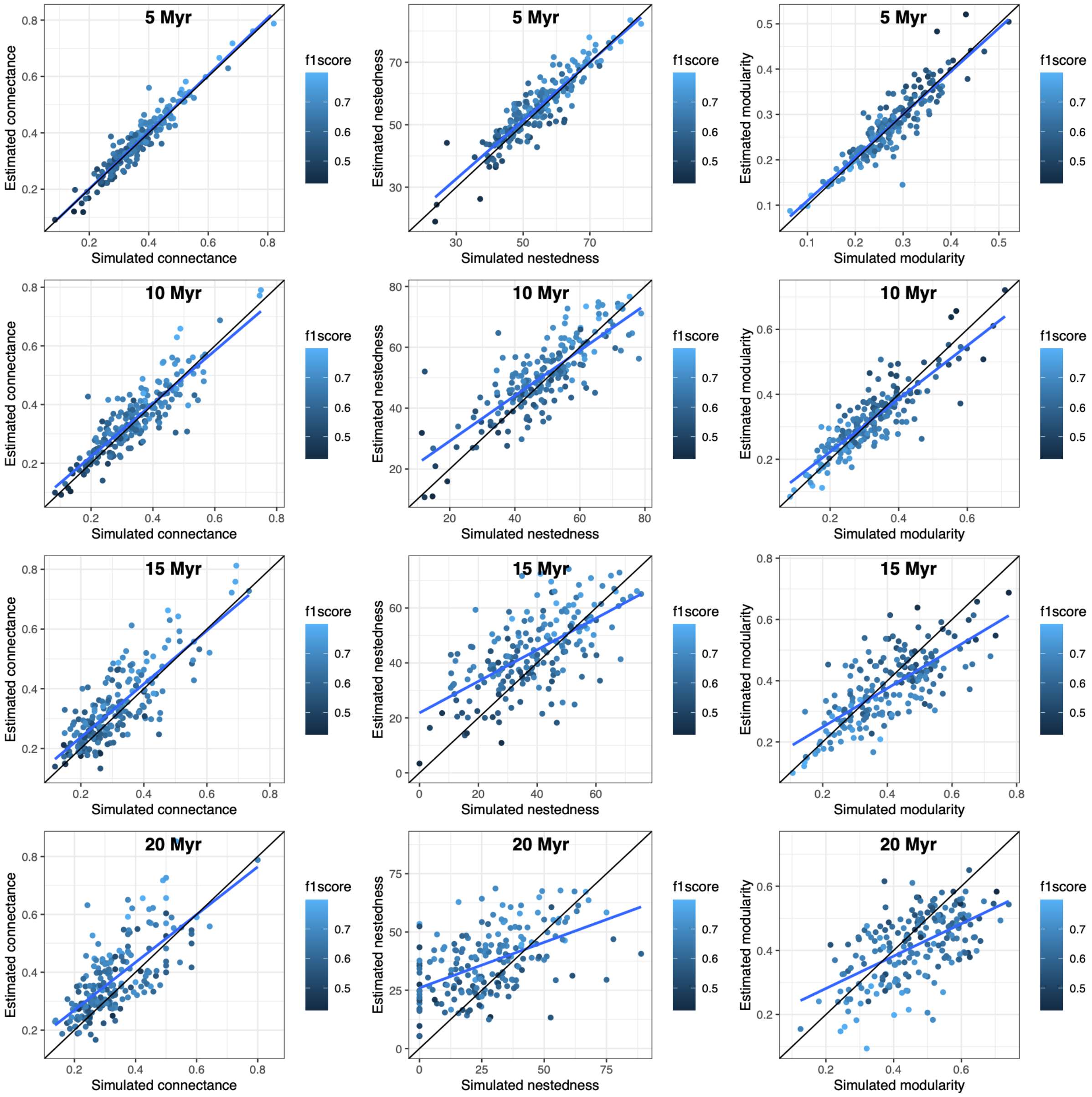

### d) Host repertoire evolution

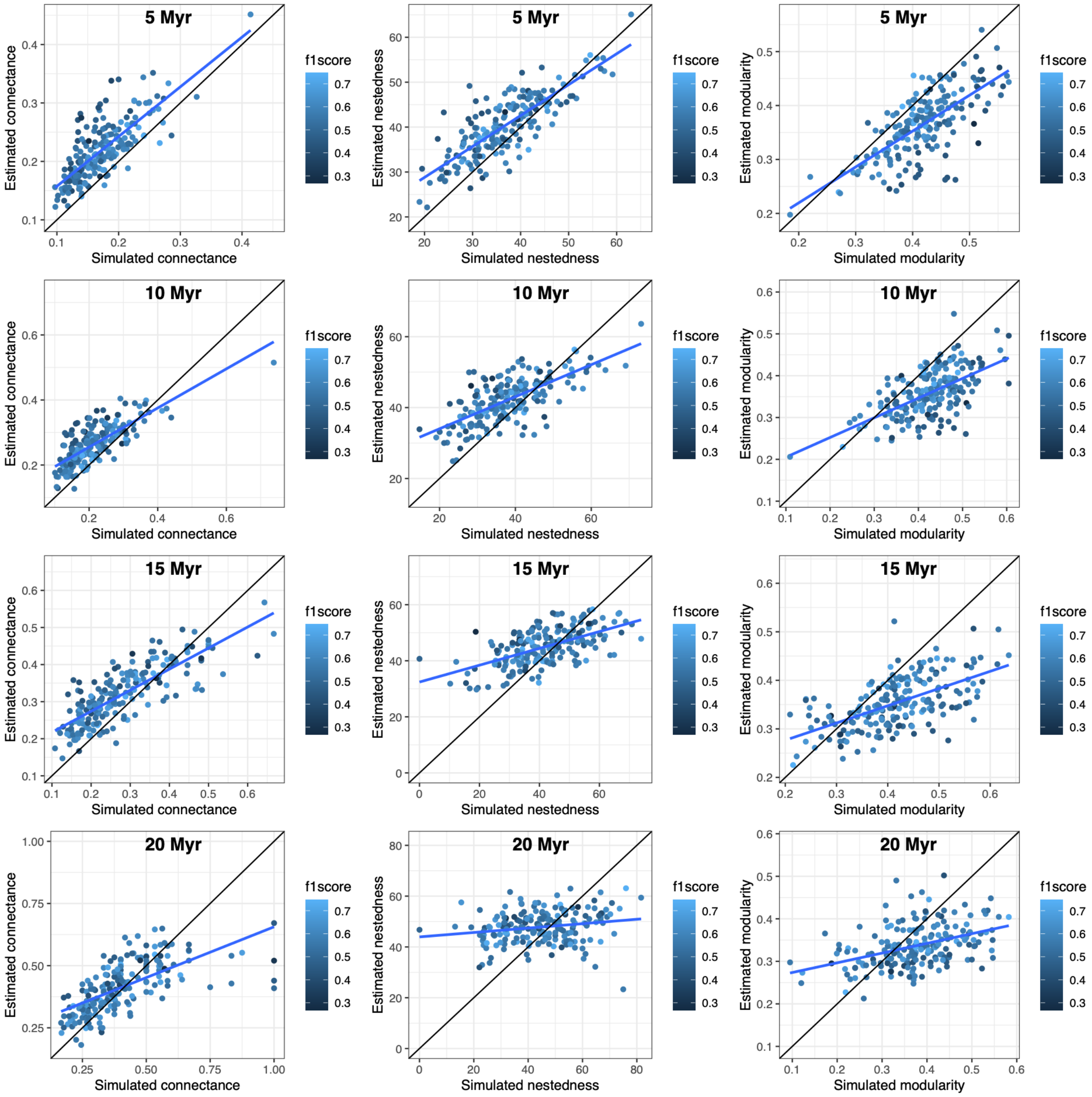

### e) Random evolution

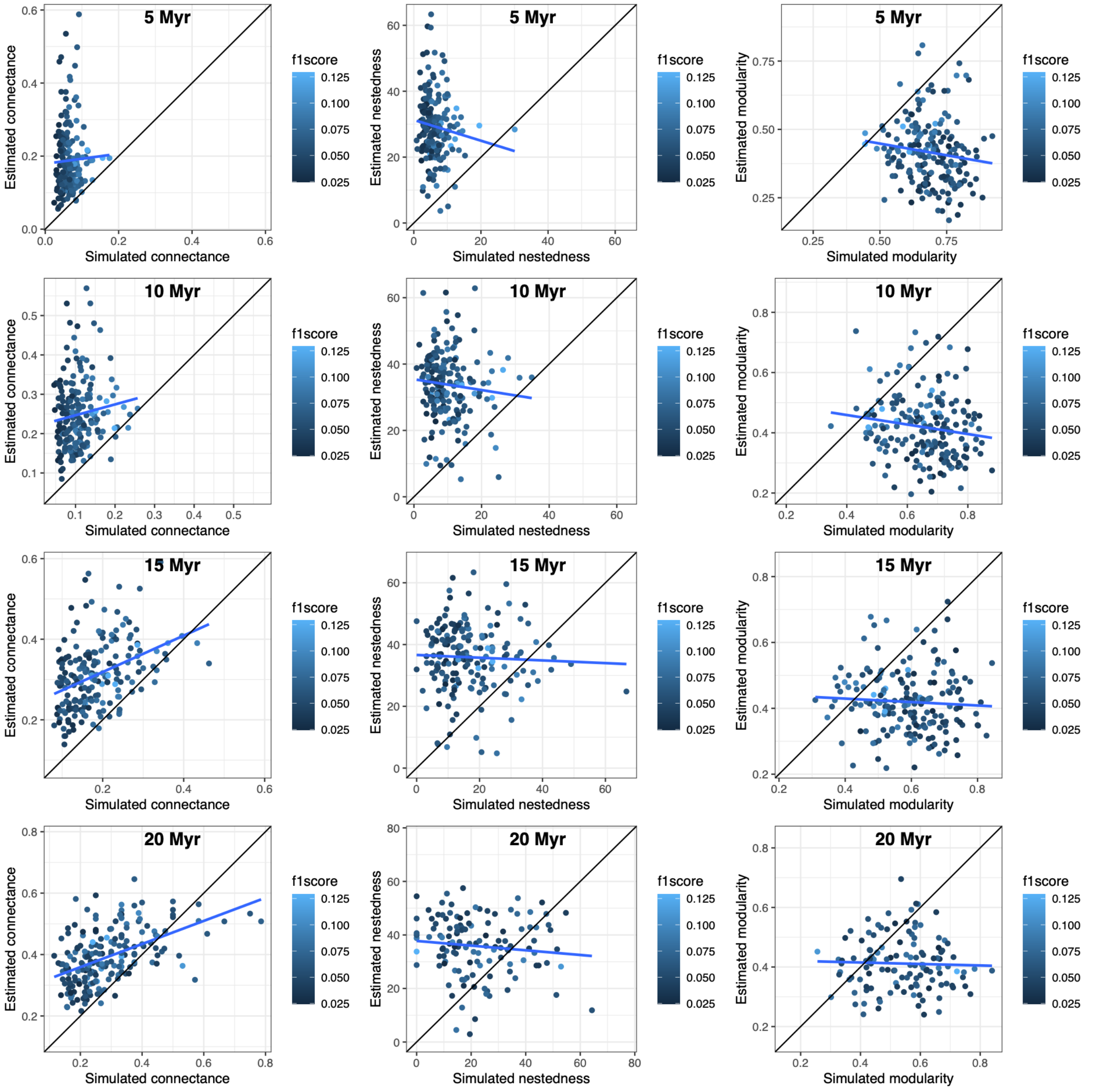

**Supplementary Figure 6: Estimations of the phylogenetic signals in ancestral networks as a function of the simulated ones for the different types of simulations using ELEFANT.**

The estimations of the phylogenetic signals in guild A (hosts; left panels) and B (symbionts; right panels) are represented at different times in the past (from 5 to 20 Myr ago).

Phylogenetic signals are measured using Mantel tests with Pearson correlations.

Each dot corresponds to one simulation, and for each simulation, we reported the estimated values averaged over 100 'augmented' ancestral networks.

Dot colors indicate the significance of the simulated phylogenetic signal, while shapes indicate the significance of the inferred signals.

The black lines represent the line y=x, while the blue lines correspond to the fit of a linear model between the estimated and simulated values.

### a) Network evolution by anagenetic shifts

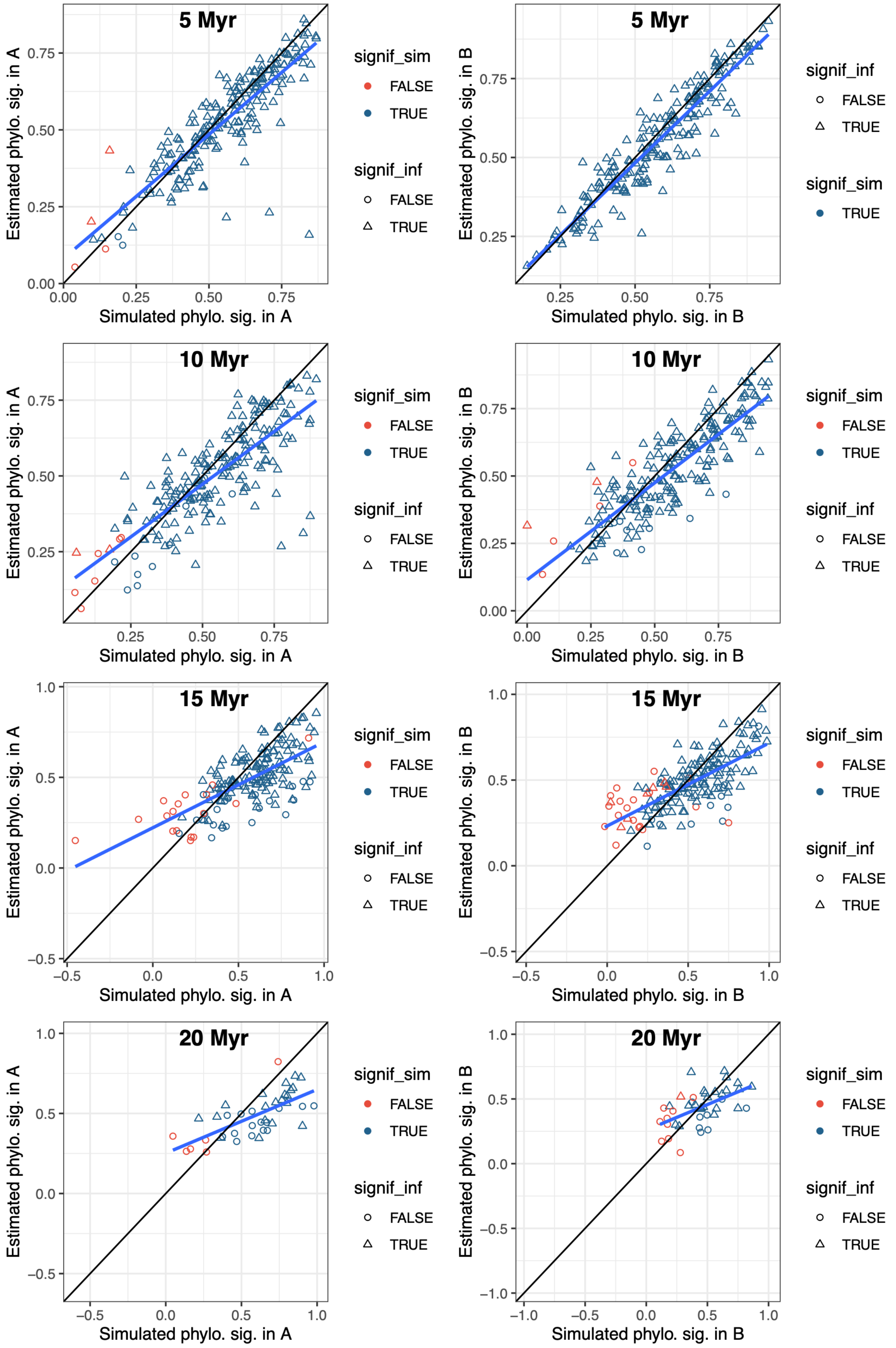

**b) Network evolution by cladogenetic shifts**

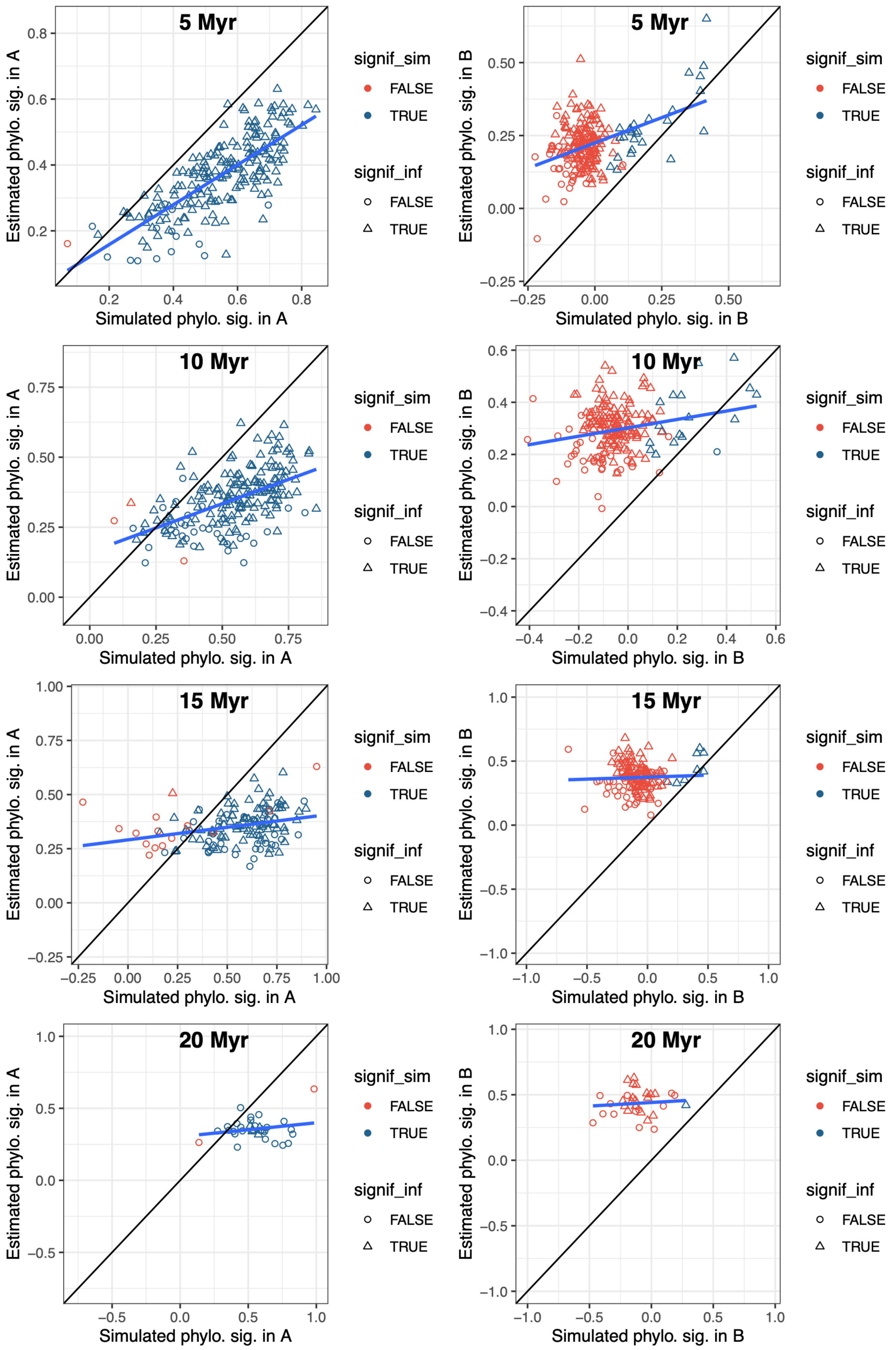

#### c) Evolution of latent traits by Brownian motions

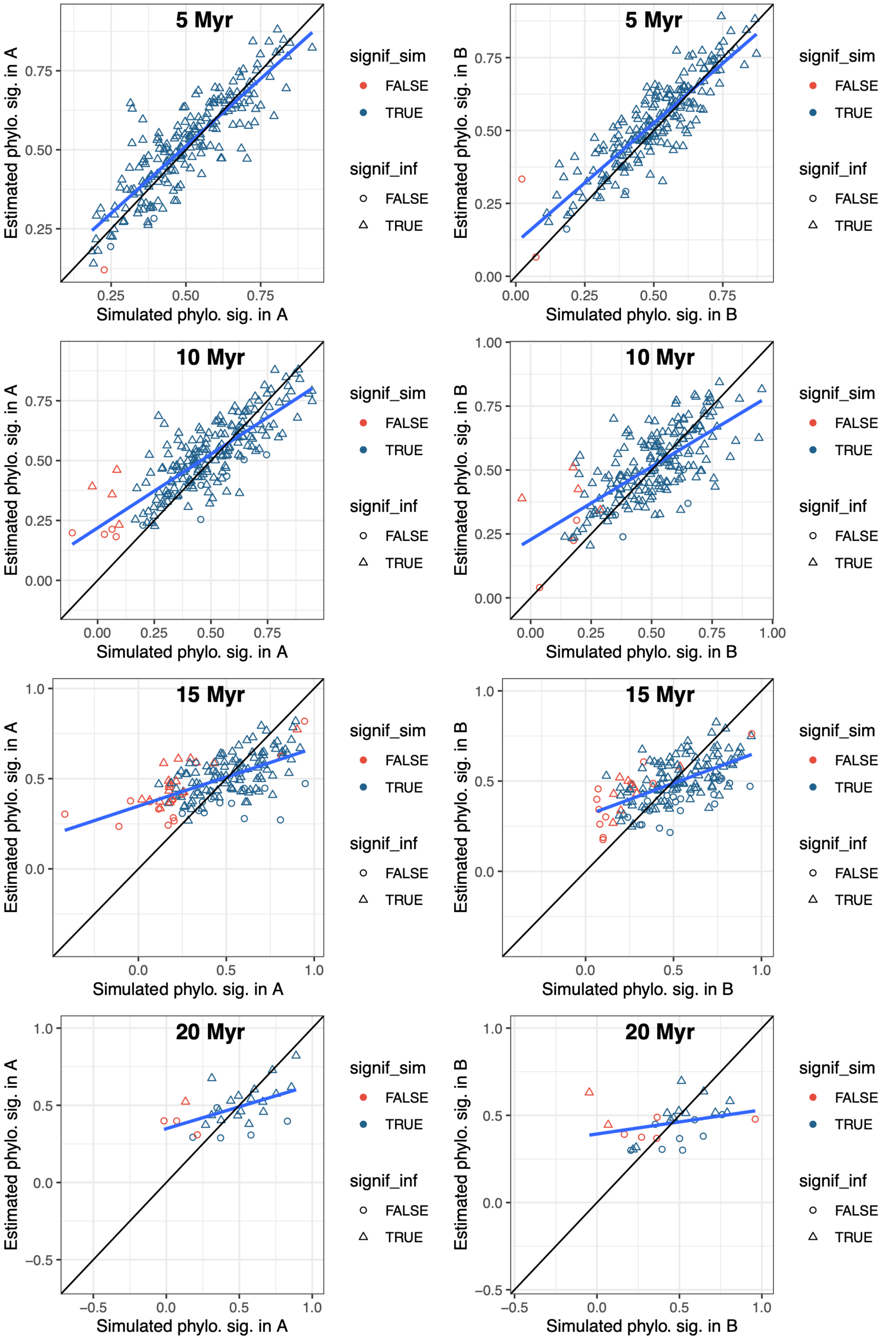

**d) Host repertoire evolution**

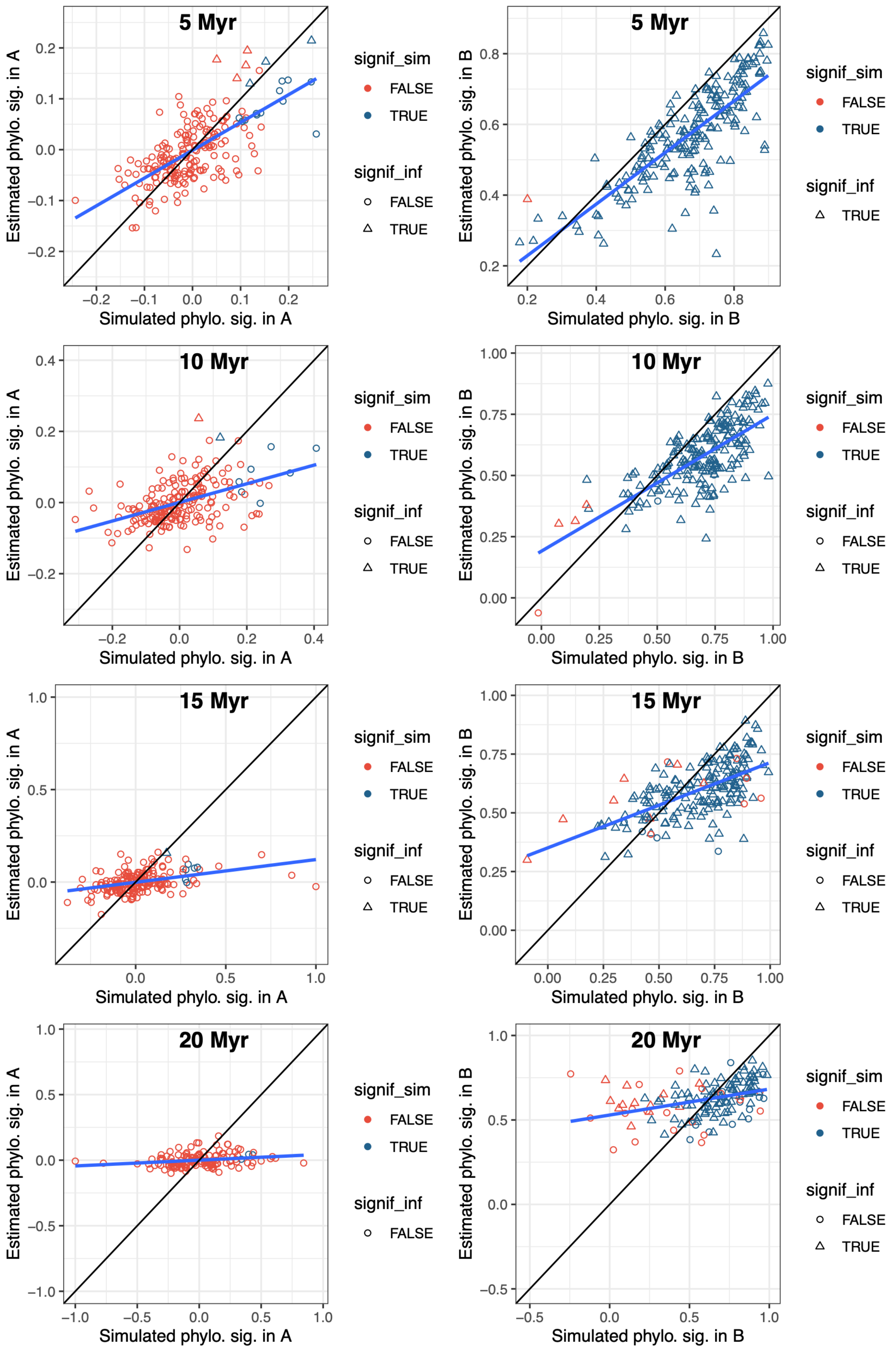

**e) Random evolution**

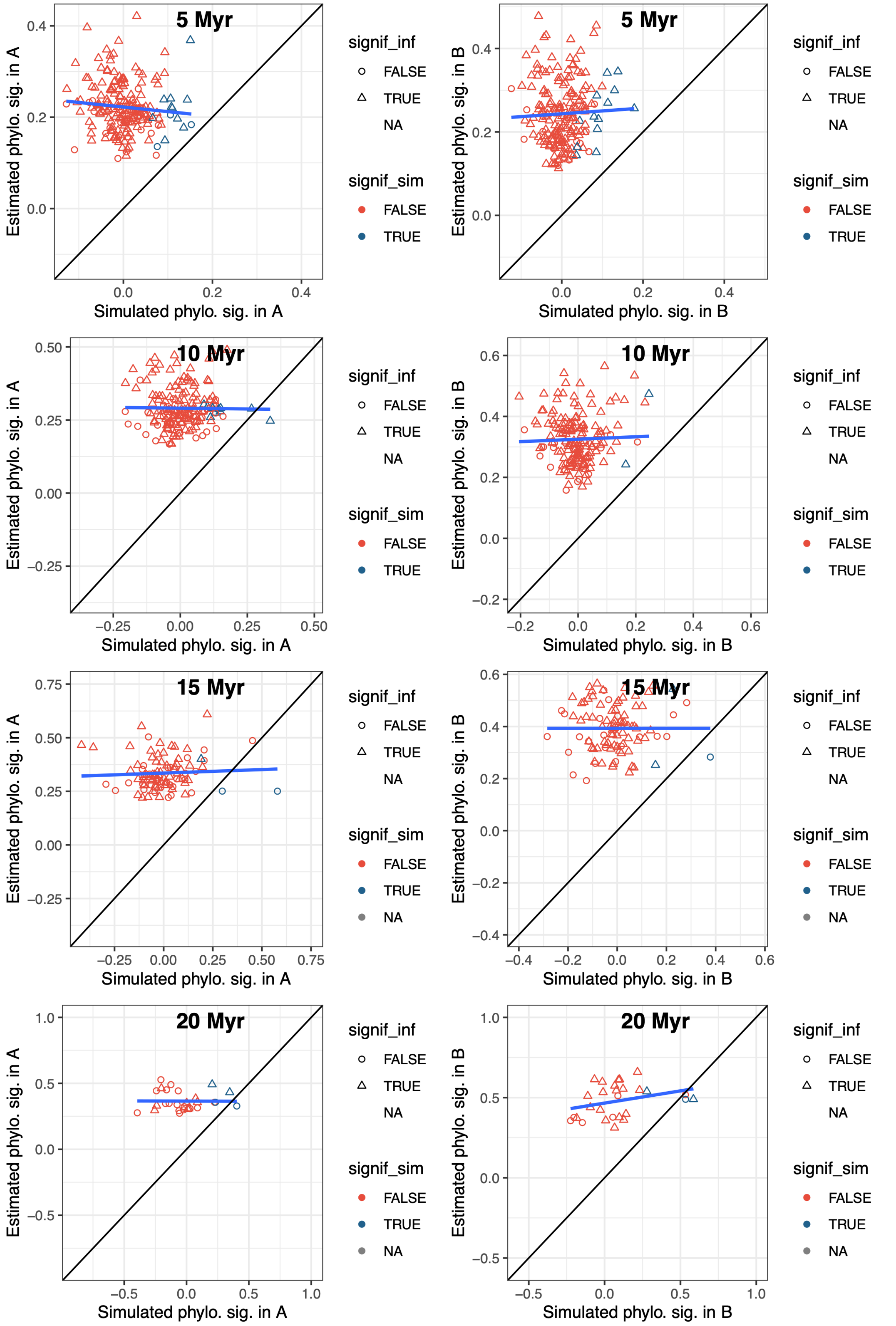

**Supplementary Figure 7: Temporal trends in the simulated or estimated global metrics of the ancestral networks for the different types of simulations using ELEFANT.**

The simulated and estimated values for the number of species in guild A (hosts; panels A and B), in guild B (symbionts; panels C and D), the connectance (panels E and F), the nestedness (panels G and H), and the modularity (panels I and J) are represented as a function of time.
Each dot corresponds to one simulation, and for each simulation, we reported the estimated values averaged over 100 'augmented' ancestral networks. Dots corresponding to the same simulation at different times are connected by a grey line.
The dot color represents the cross-validation scores. The blue lines correspond to the fits of linear models.

### a) Network evolution by anagenetic shifts

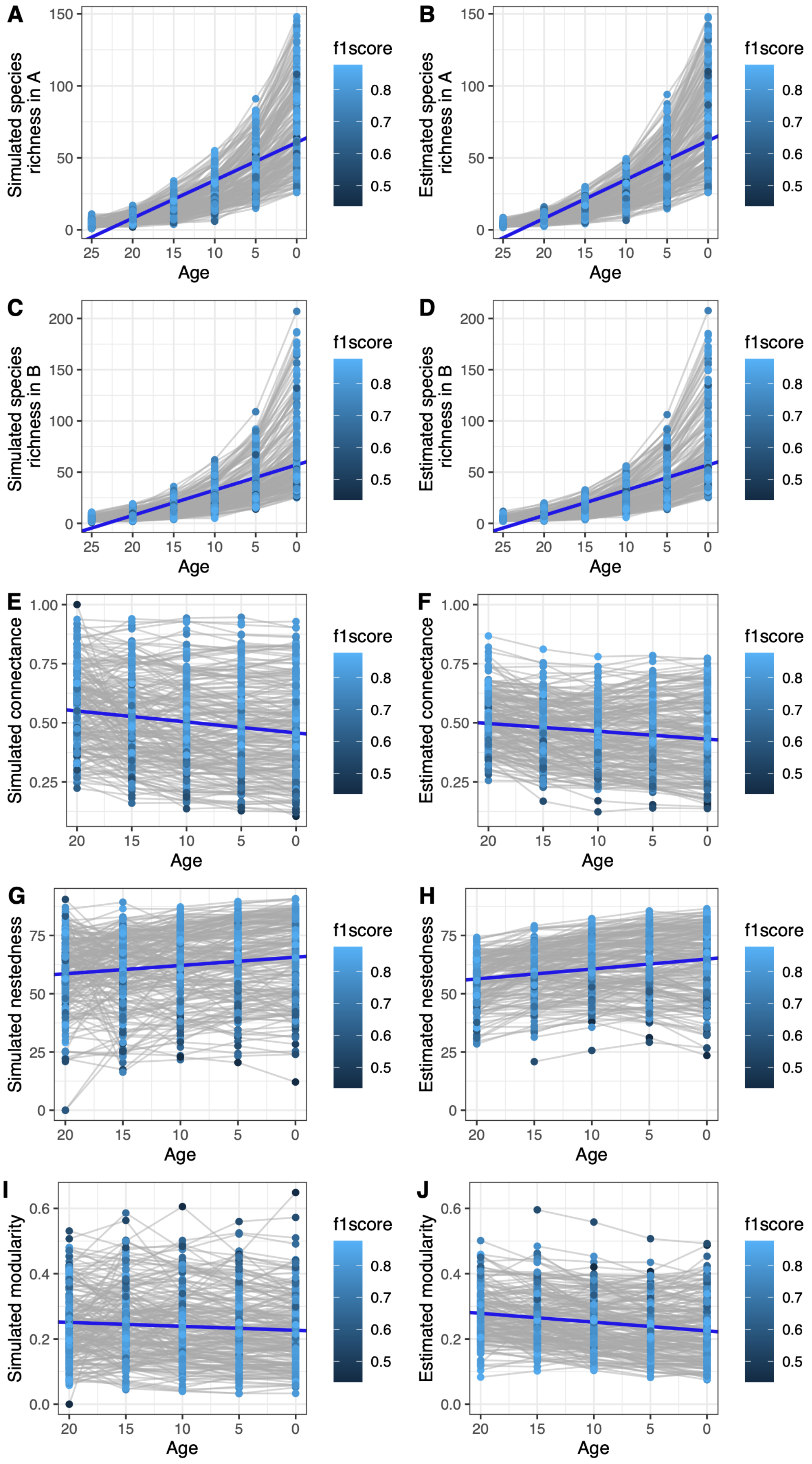

### b) Host repertoire evolution

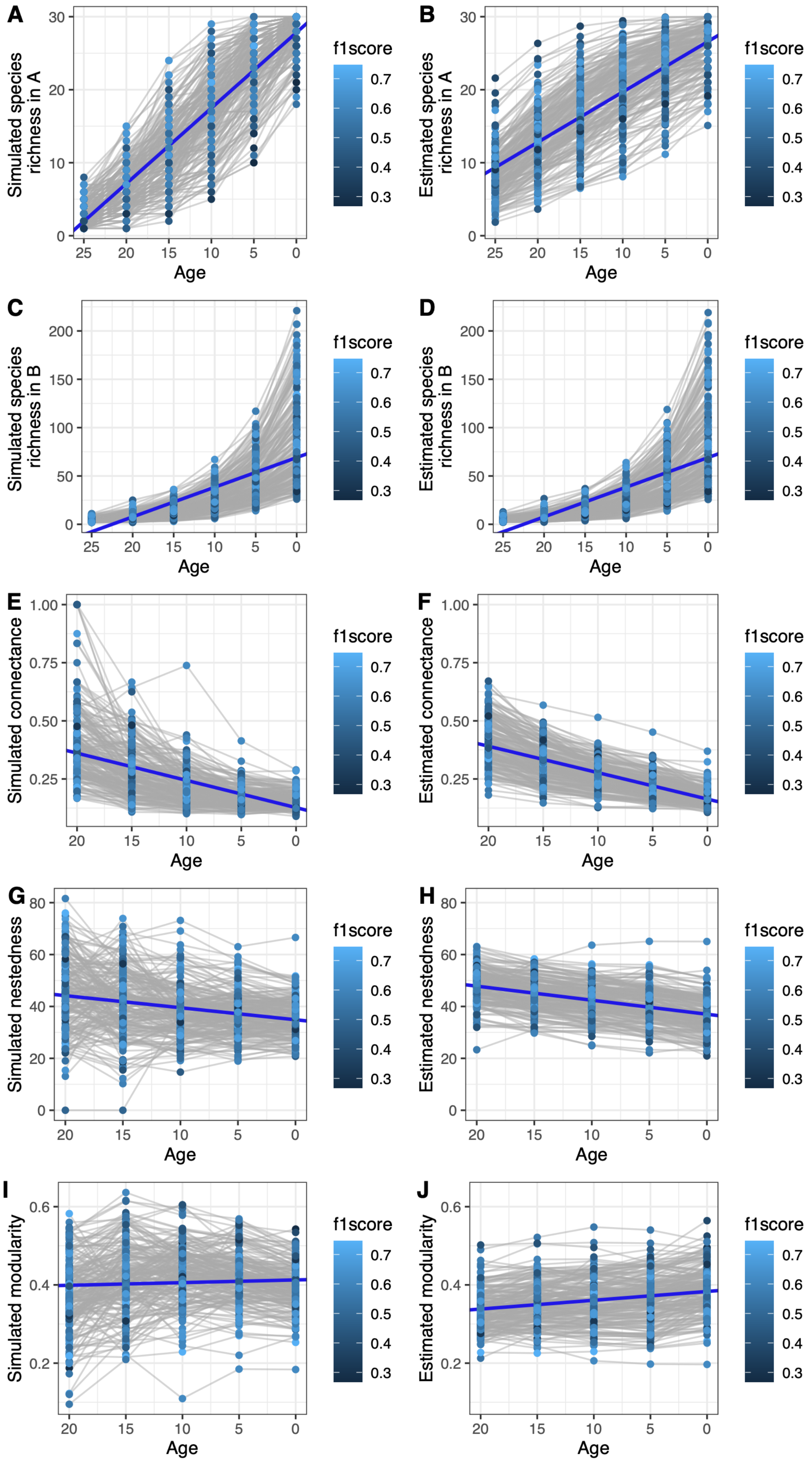

### c) Network evolution by cladogenetic shifts

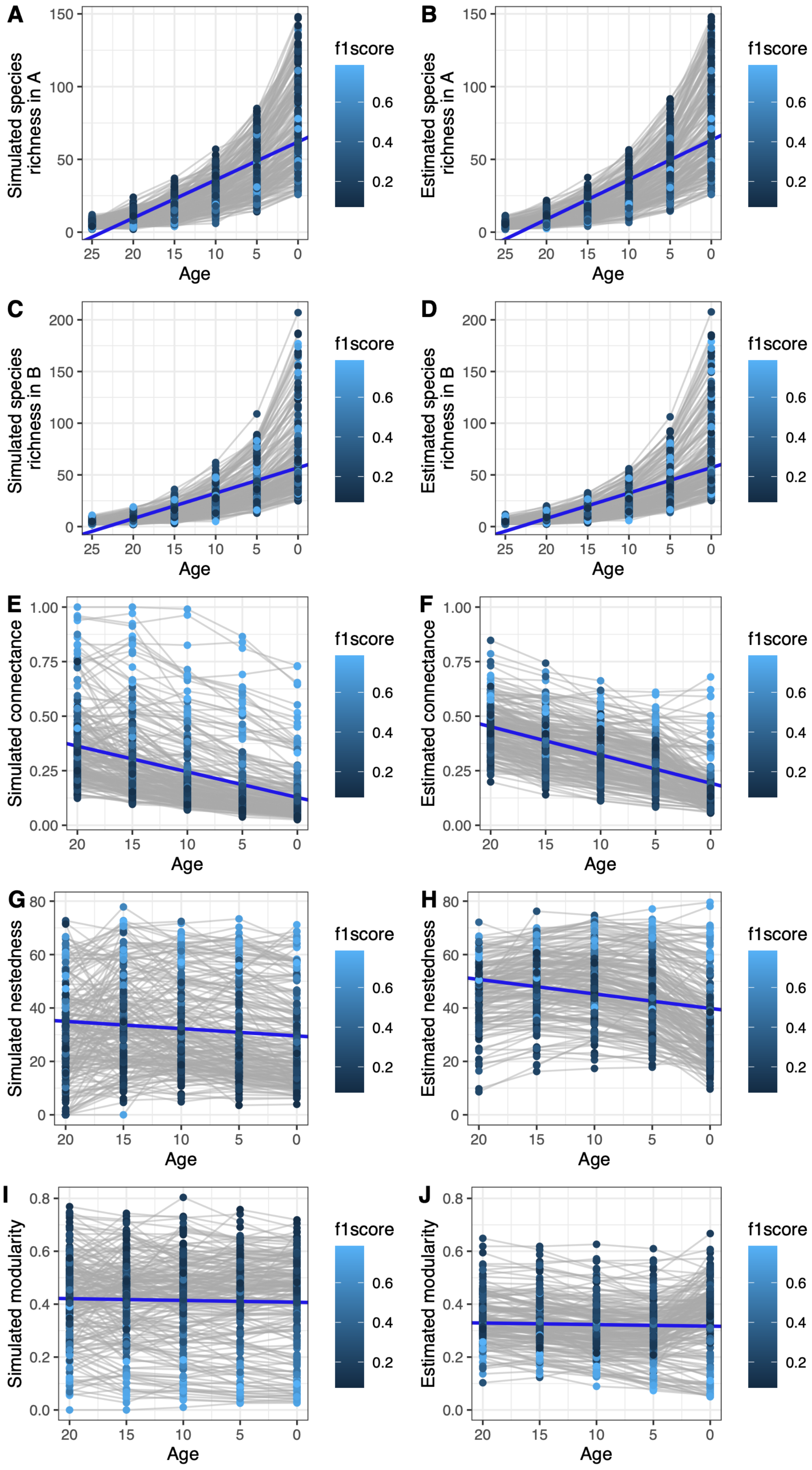

**d) Evolution of latent traits by Brownian motions**

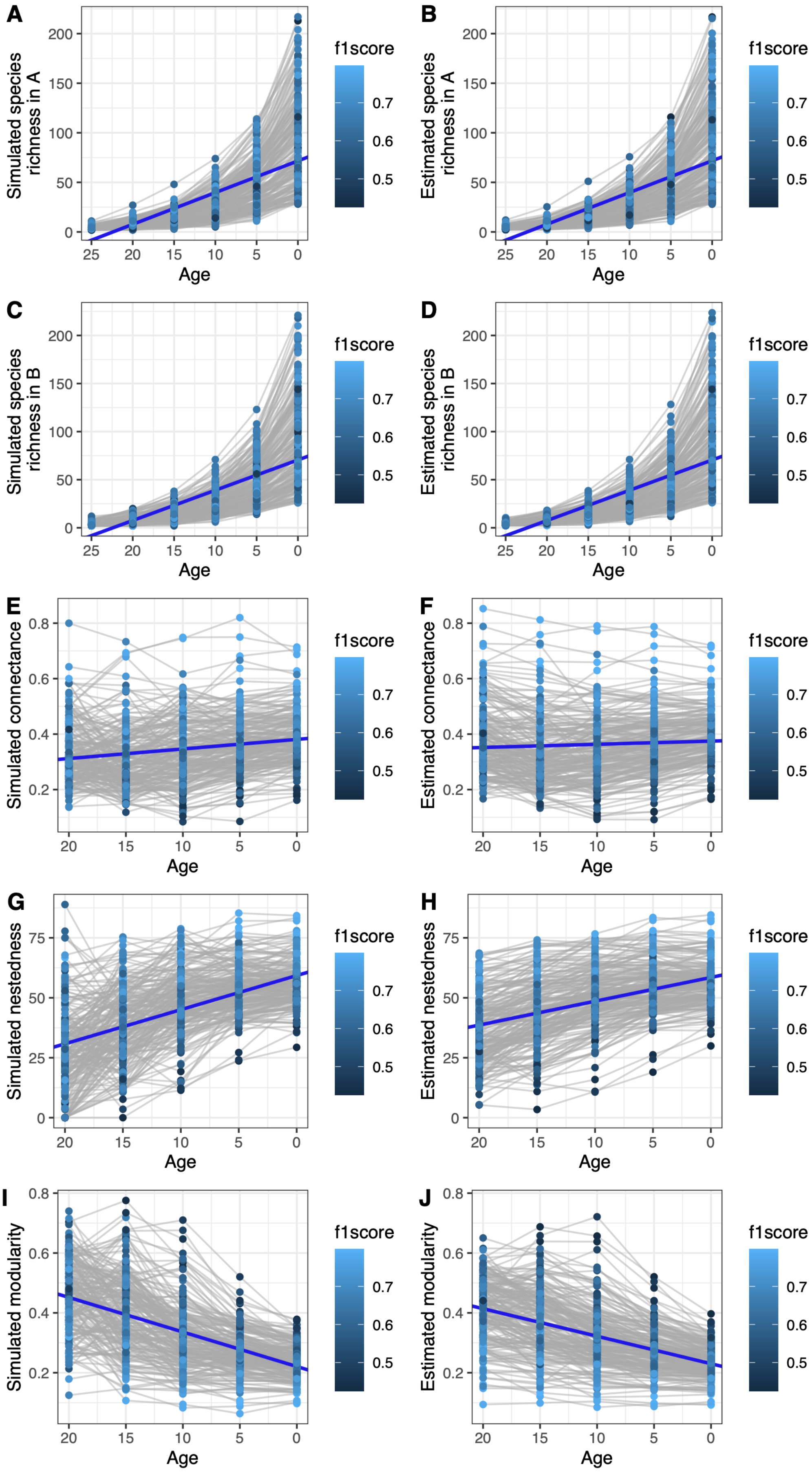

e) Random evolution

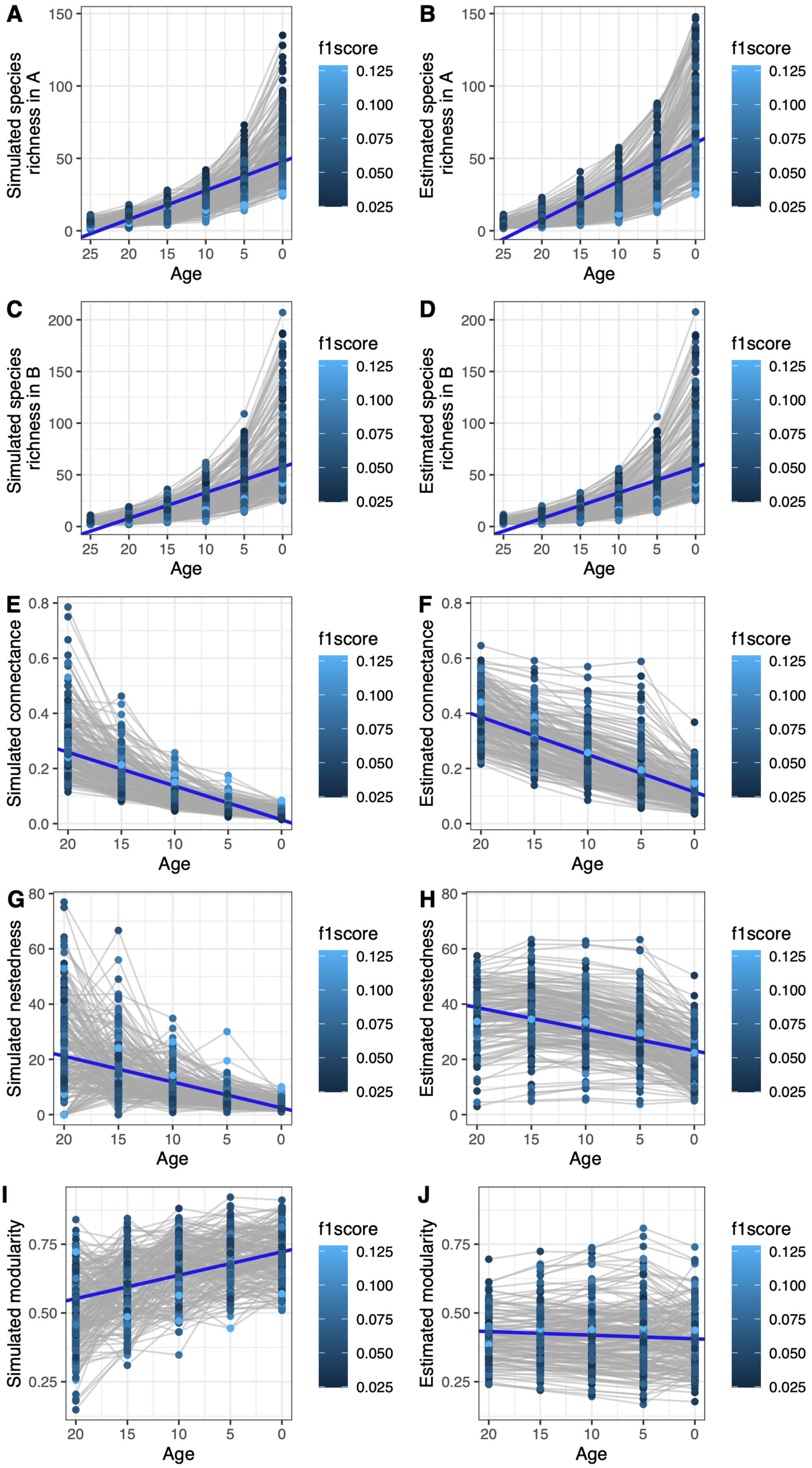

**Supplementary Figure 8: Results of the empirical applications are qualitatively similar without data augmentation or when using different rates of diversification for data augmentation:**

We replicated the analyses using ELEFANT without data augmentation (a, c, e, g) or with different rates of diversification data augmentation (b, d, f, h; see Table S4 for the rate values).

The different subplots represent the temporal variations of ancestral network properties. Squares indicate a significant departure from the null models (95% central range shown in orange), whereas the crosses indicate non-significance. Error bars represent the 95% central range of values obtained from the set of augmented ancestral networks.

**a) Mycorrhizal networks without data augmentation**

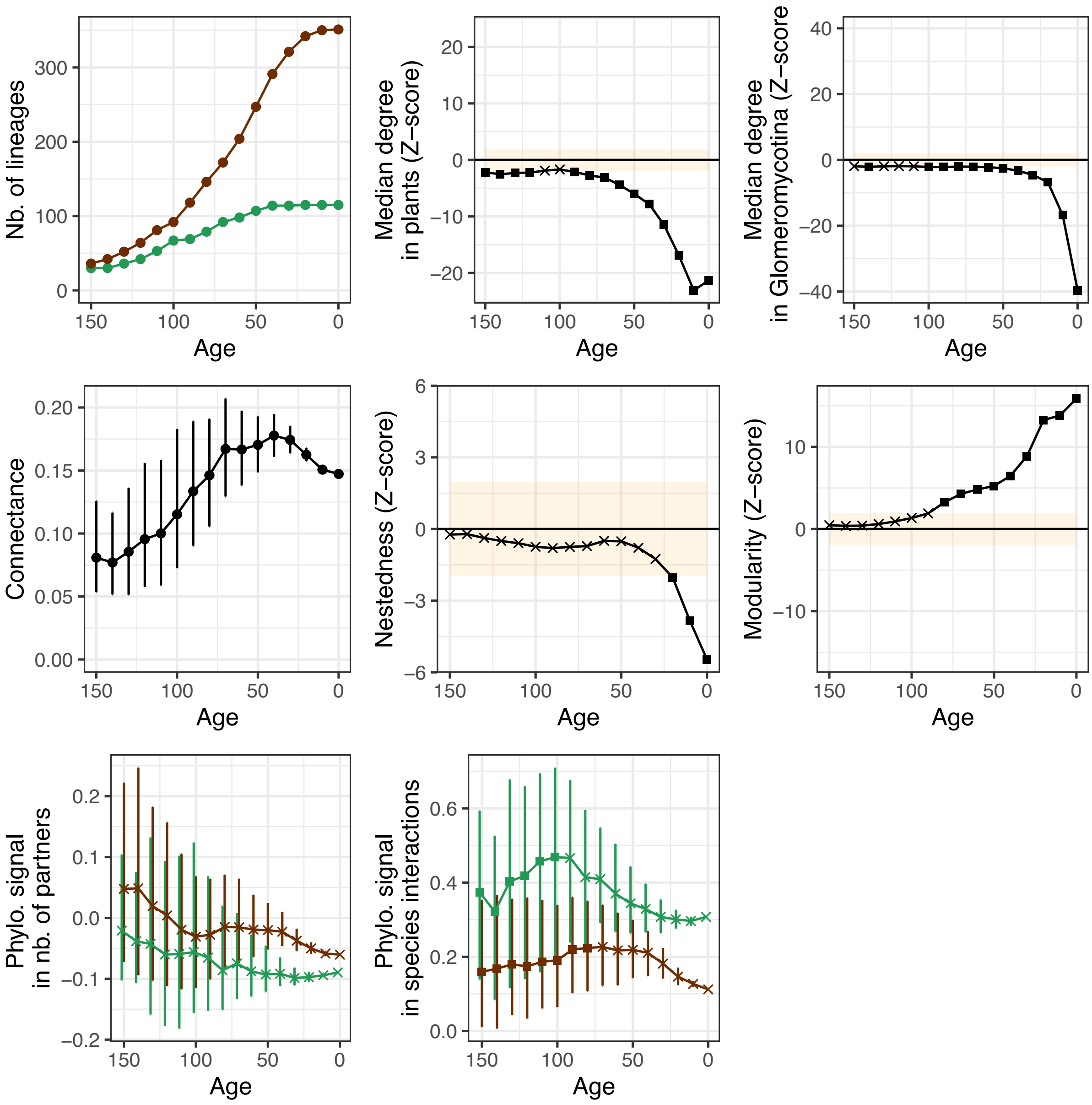

**b) Mycorrhizal networks with different rates for data augmentation**

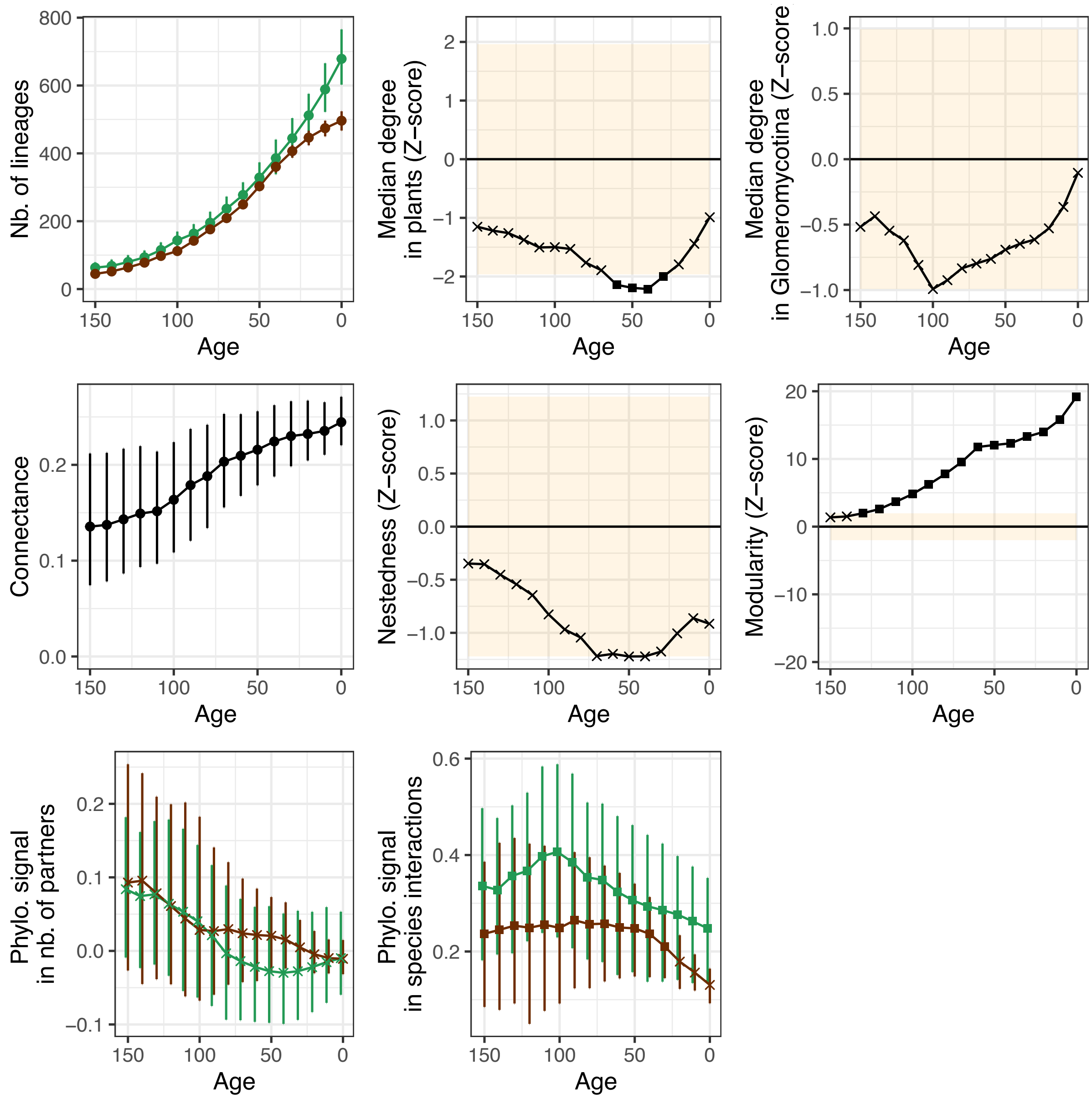

**c) Bat pollination networks without data augmentation**

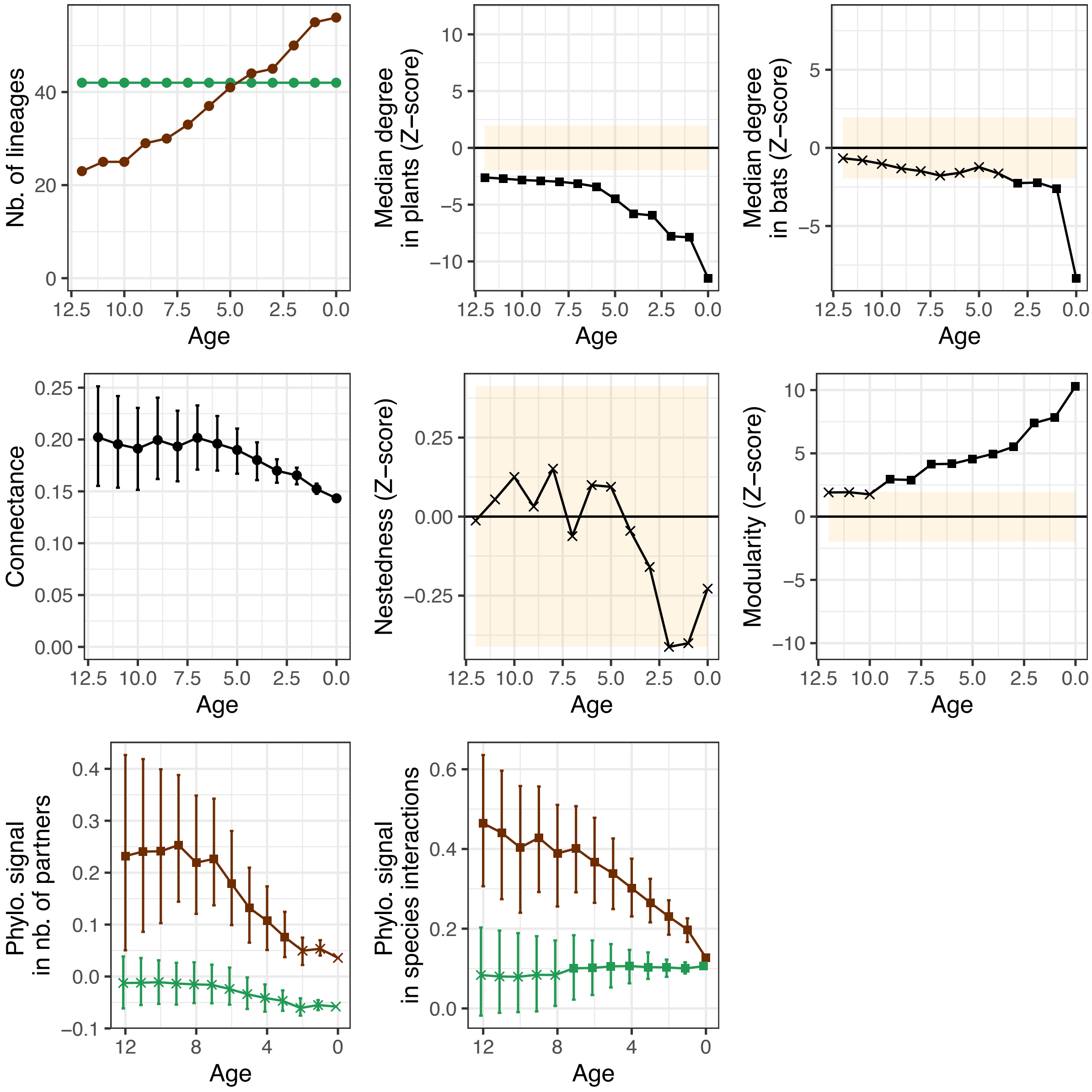

**d) Bat pollination networks with different rates for data augmentation**

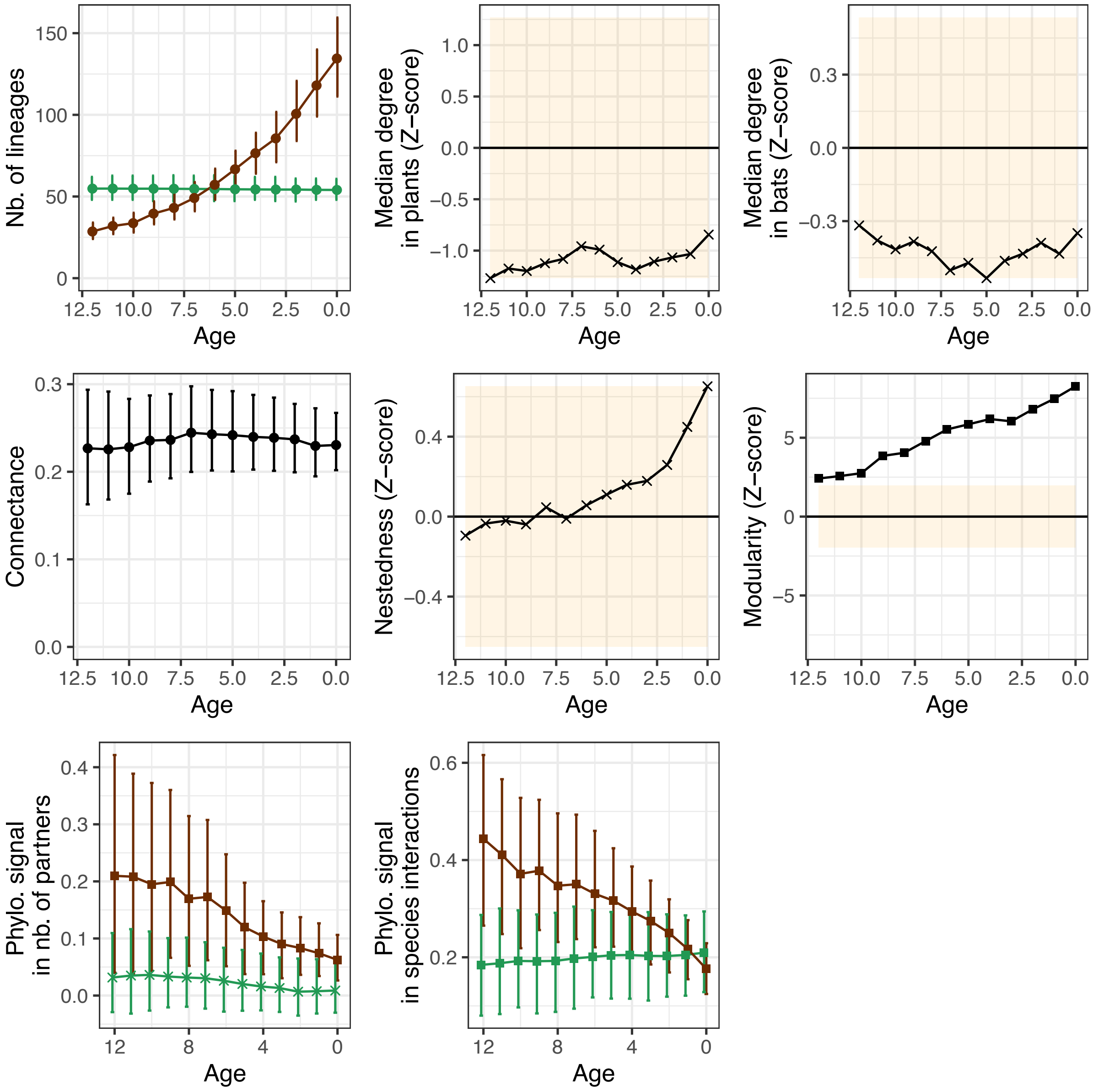

**e) Plant-bird seed dispersal networks in Europe without data augmentation**

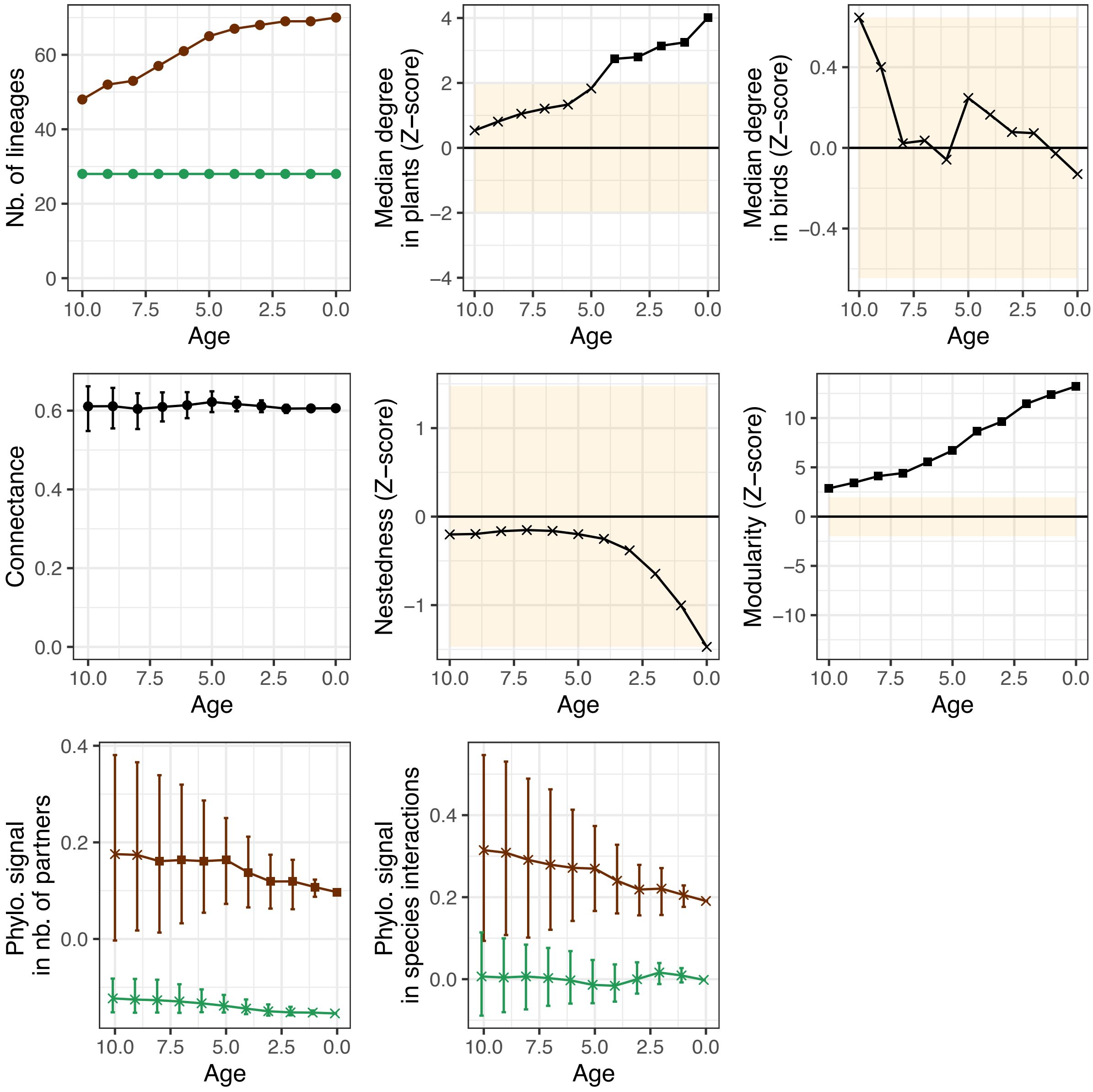

**f) Plant-bird seed dispersal networks in Europe with different rates for data augmentation**

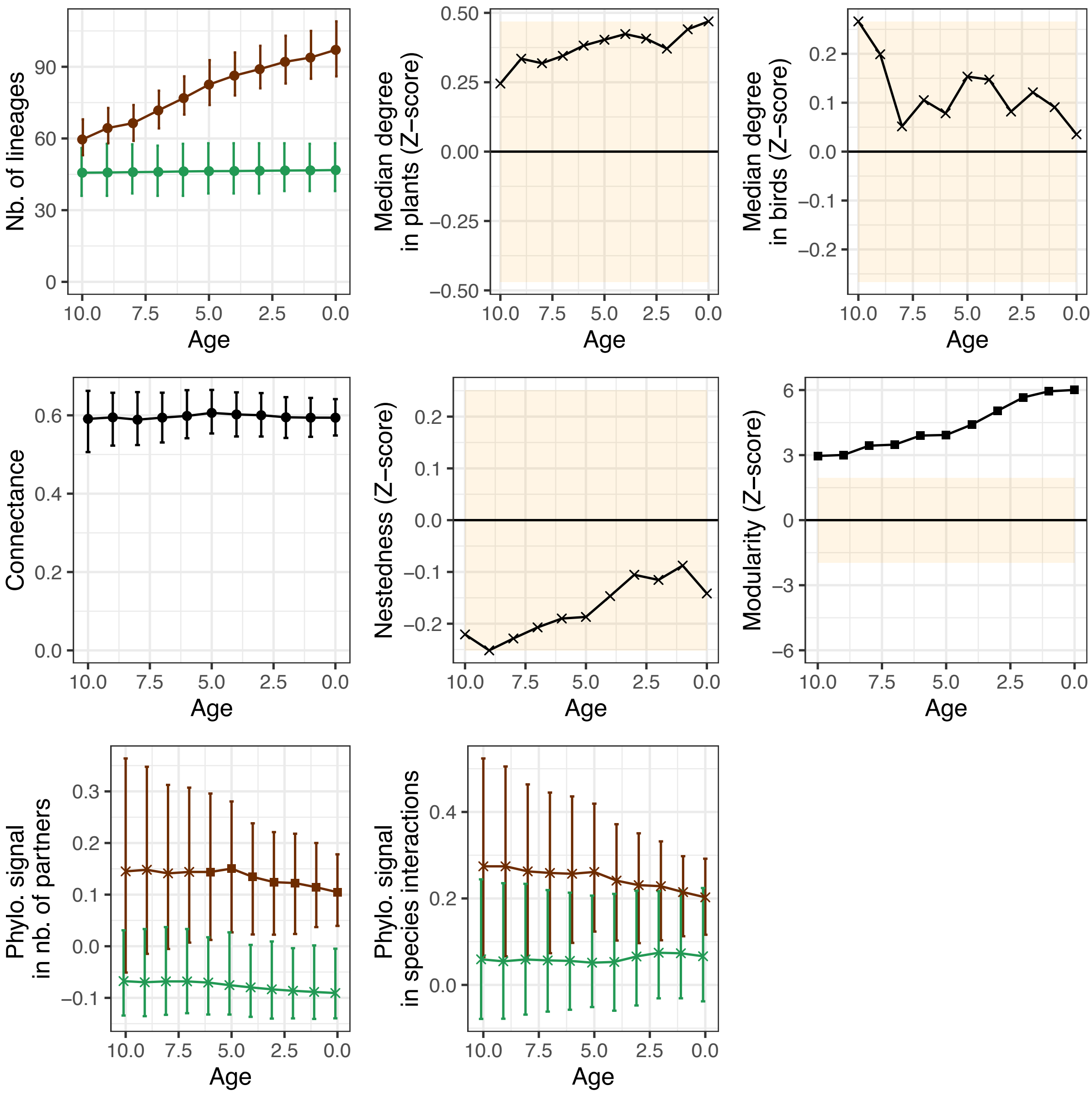

### g) Plant-bird seed dispersal networks in the Americas without data augmentation

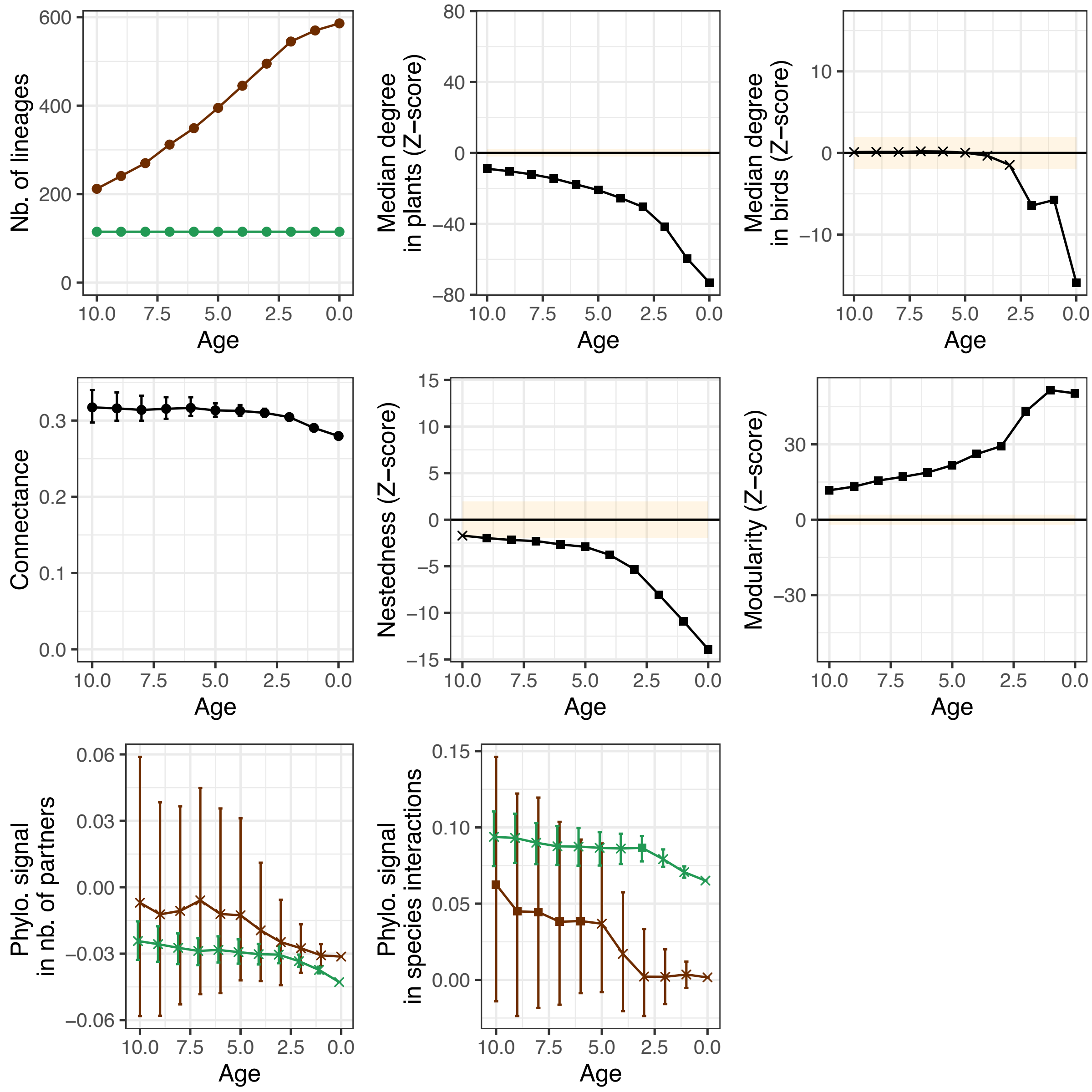

**h) Plant-bird seed dispersal networks in the Americas with different rates for data augmentation**

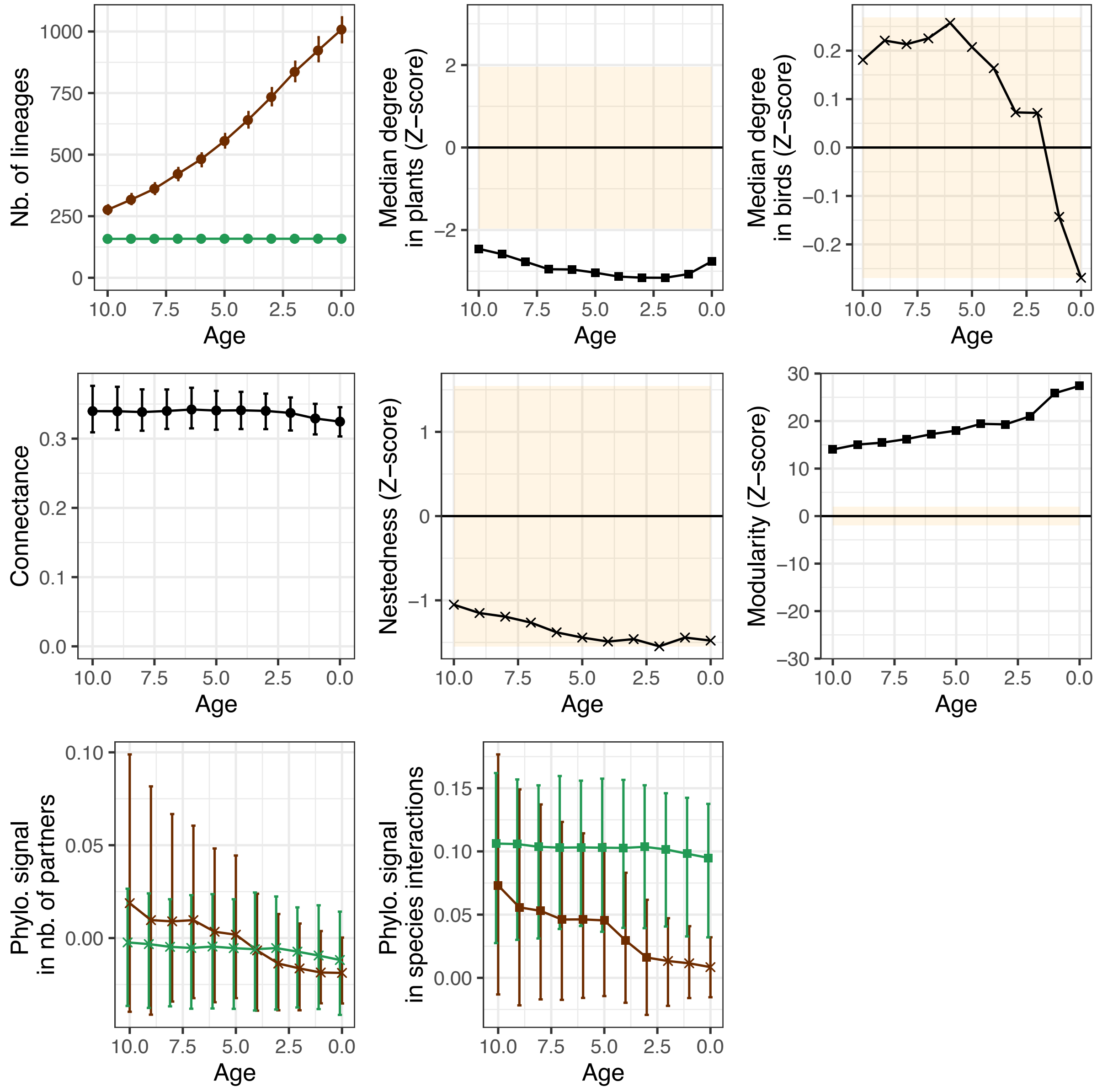

**Supplementary Figure 9: Relationship between the sum of the latent traits and the degree of generalism of each species in the empirical mutualistic networks:**

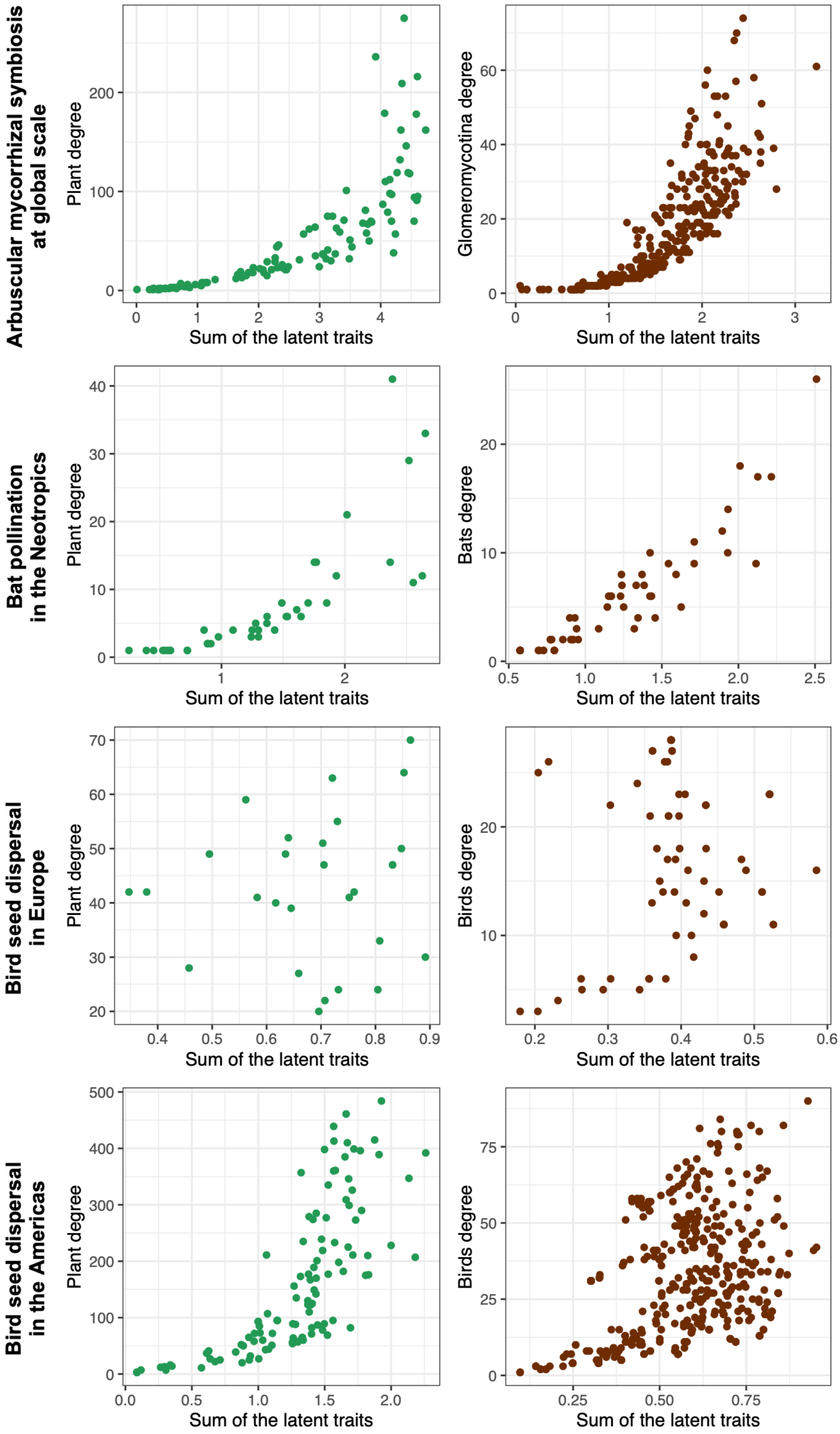

**Supplementary Figure 10: Effect of biogeography on the latent traits of species across different empirical mutualistic networks.**
For each latent trait, the barplot shows the result of a PGLS testing the effect of the species bioregions on the latent trait values. Effects were considered significant at p<0.01 to account for multiple testing.

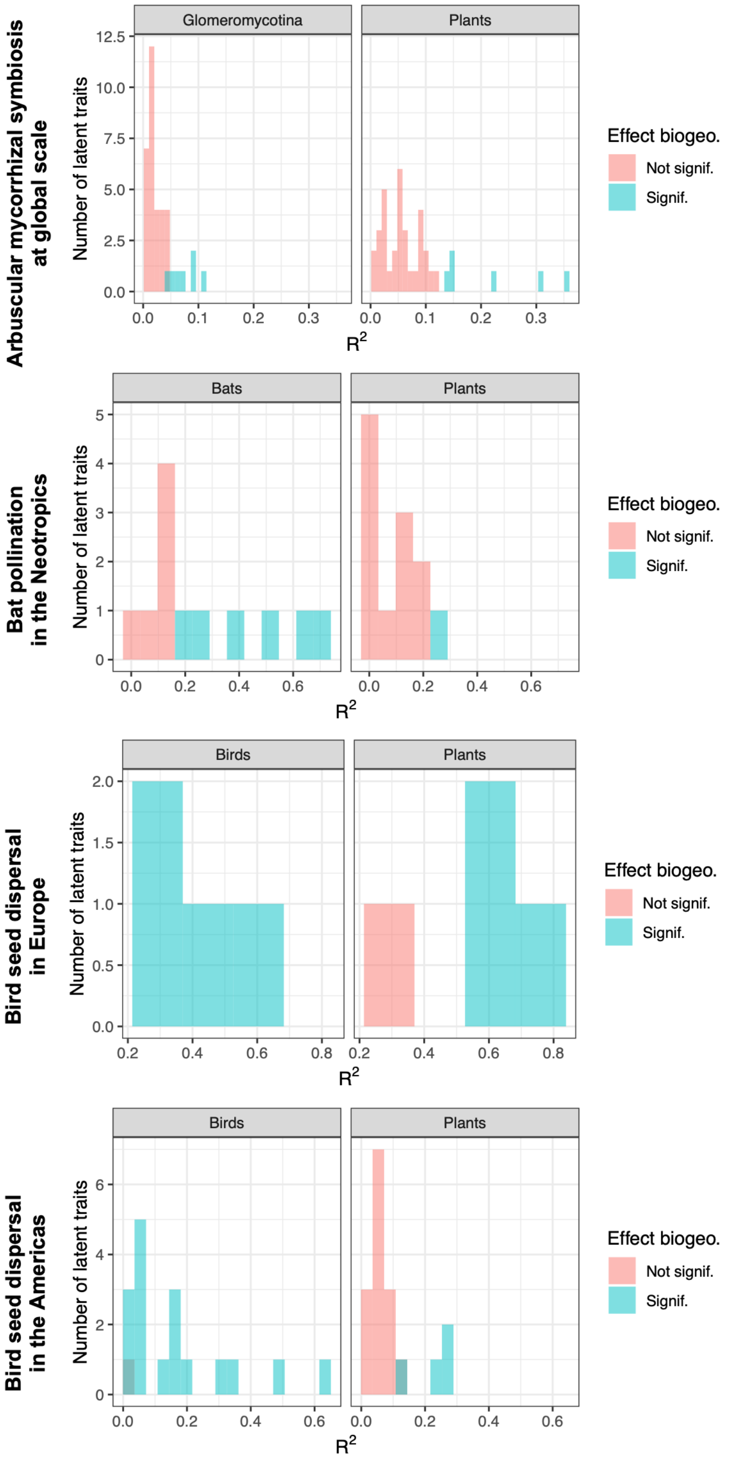

**Supplementary Figure 11: Present-day phylogenetic signals in present-day networks for the different types of simulations.**

Phylogenetic signals are measured using Mantel tests with Pearson correlations, which are reported in guild A (hosts; left panels) and B (symbionts; right panels).

**a) Network evolution by anagenetic shifts**

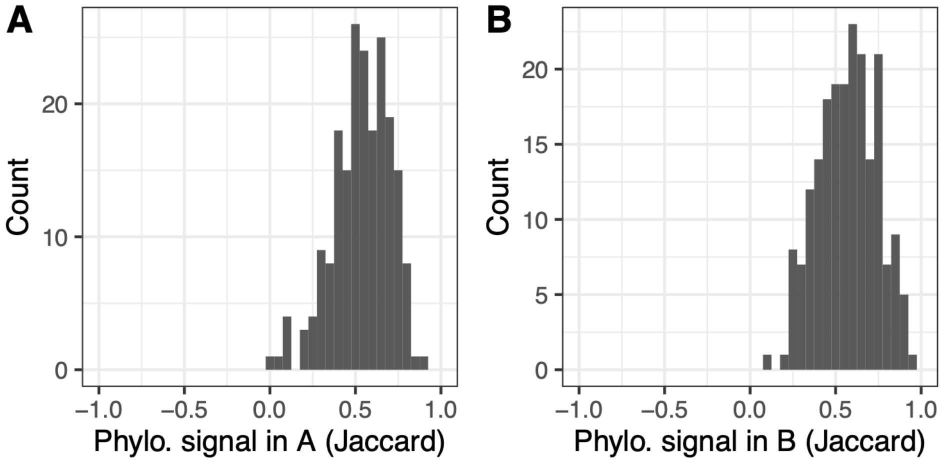


**b) Network evolution by cladogenetic shifts**

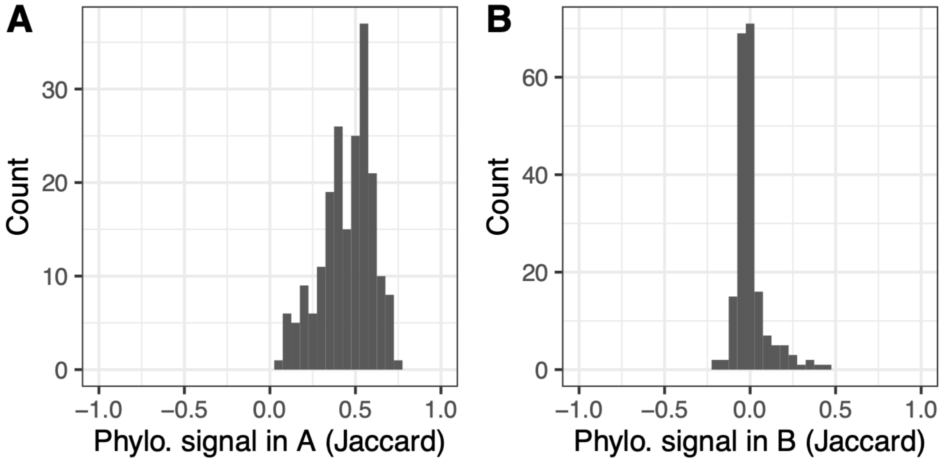


**c) Evolution of latent traits by Brownian motions**

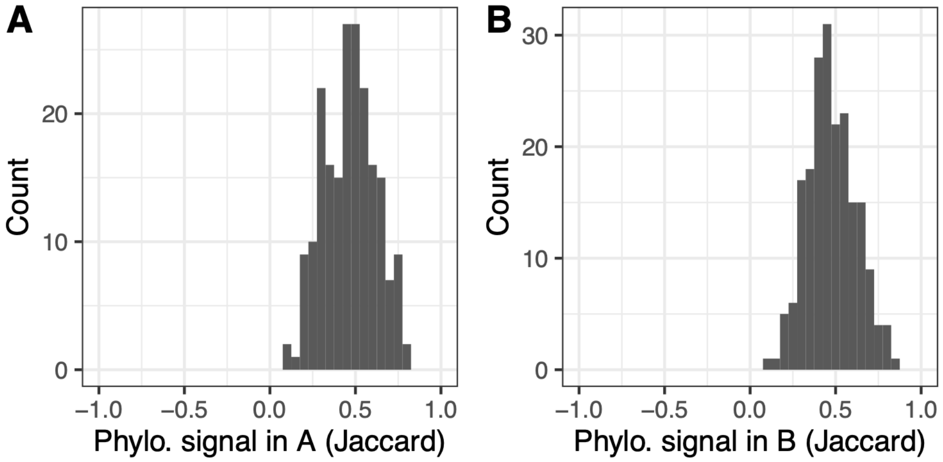

**d) Host repertoire evolution**

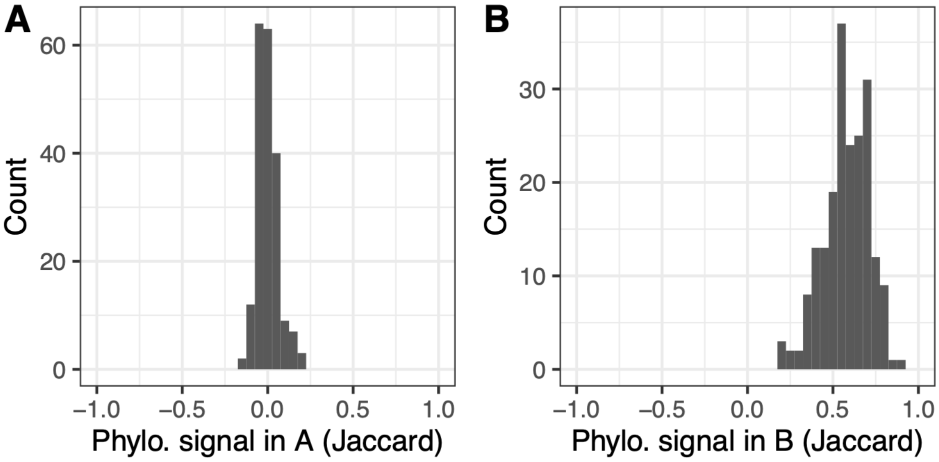


**e) Random evolution**

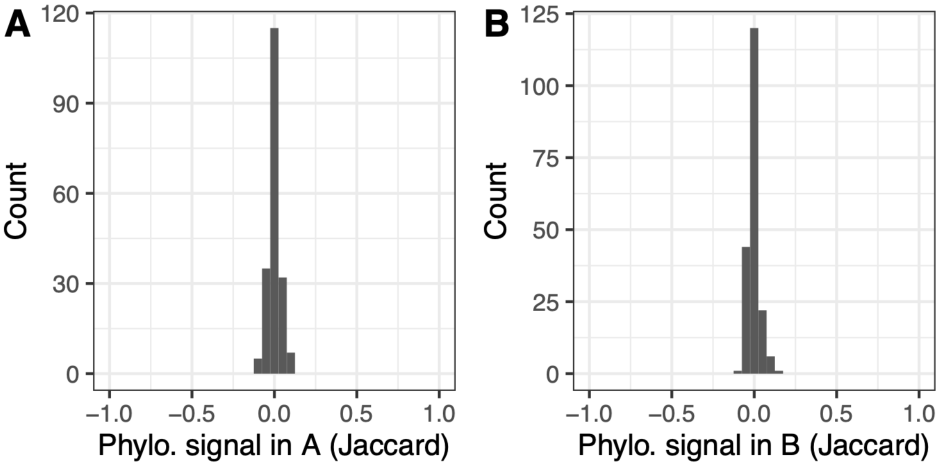

**Supplementary Figure 12: Minimum number of latent traits (*d*) explaining at least 90% of the present-day interactions in the different types of simulations using ELEFANT.**

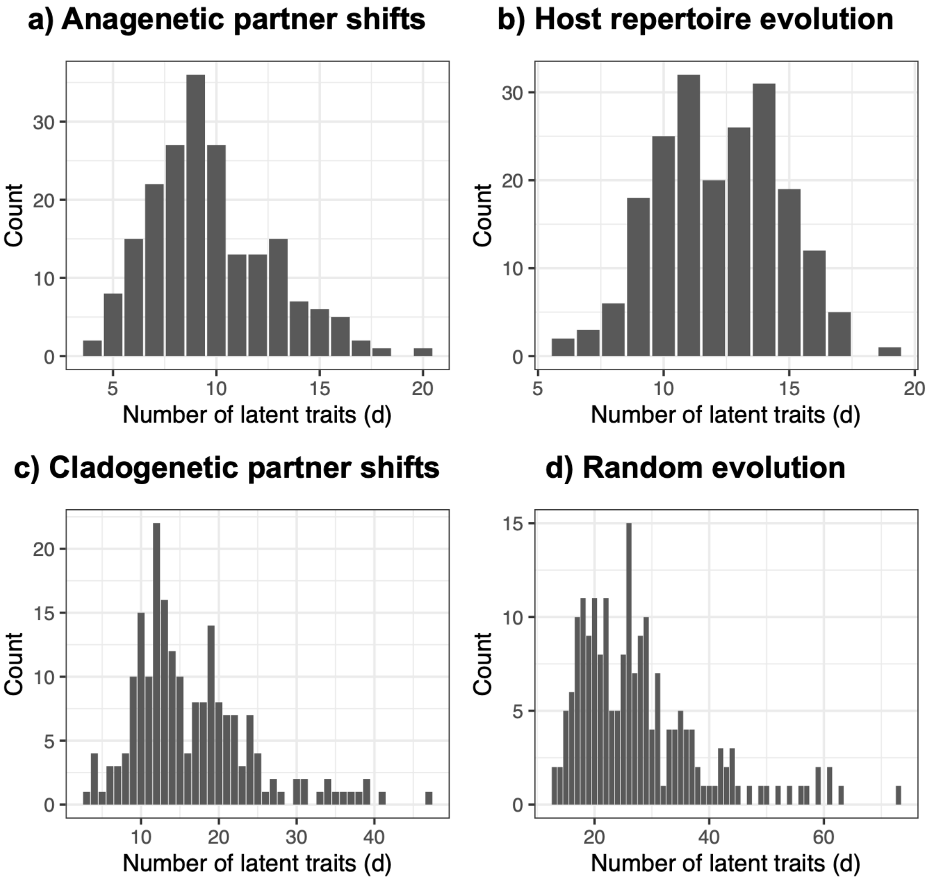

**Supplementary Figure 13: Ancestral global metrics are generally better estimated when adding extinct and unsampled lineages using data augmentation:**

The plots indicate the $R^2$ coefficients of determination of the ancestral global metrics estimated at different time in the past for the different types of simulations ($R^2$ close to 1 indicate that the estimated global metrics are very close to the simulated ones).
The color of the dots indicates the types of simulations: network evolution by anagenetic shifts (blue), host repertoire evolution (green), network evolution by cladogenetic shifts (yellow), evolution of latent traits by Brownian motions (red), or random evolution (purple).
The shape of the dots indicates whether ELEFANT included (round) or not (triangle) data augmentation (DA). The differences with or without DA are represented by grey lines.

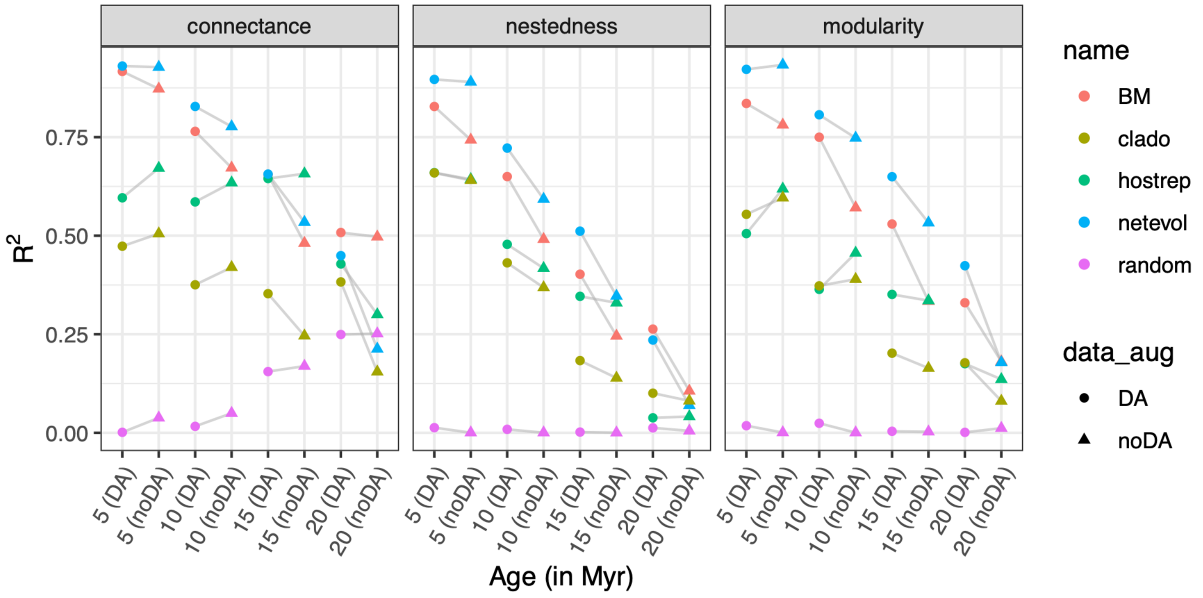

**Supplementary Figure 14: Robustness of ancestral network reconstruction to threshold choice.**

For simulations of network evolution under anagenetic changes, we ran ELEFANT using a range of threshold values (0.1-0.9) and measured the average accuracy of the estimated global metrics of the reconstructed ancestral networks ($R^2$), as well as the F1-scores of ancestral interactions (blue points). These values were compared to the average accuracies obtained using the Youden's J threshold (orange triangles).

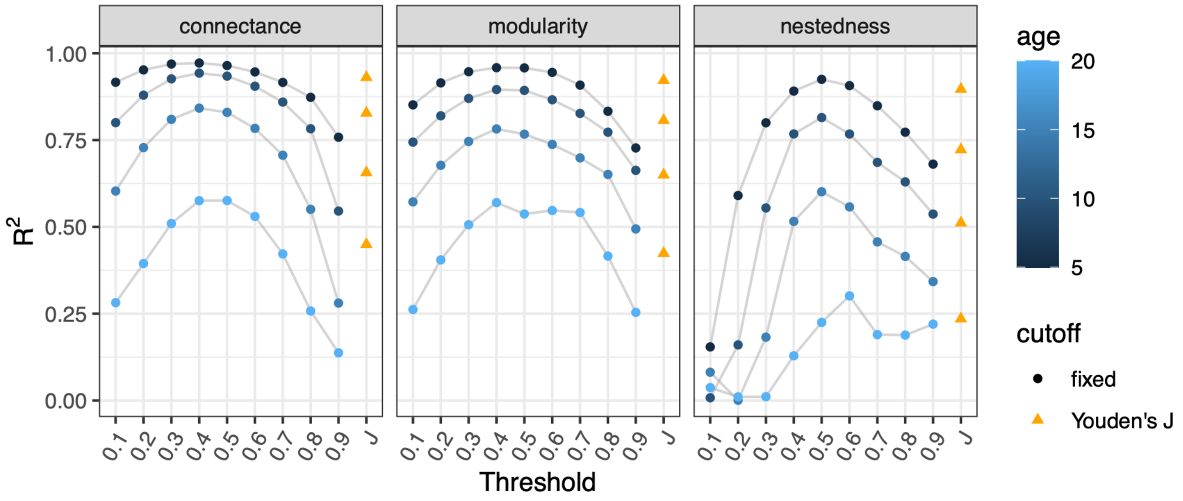


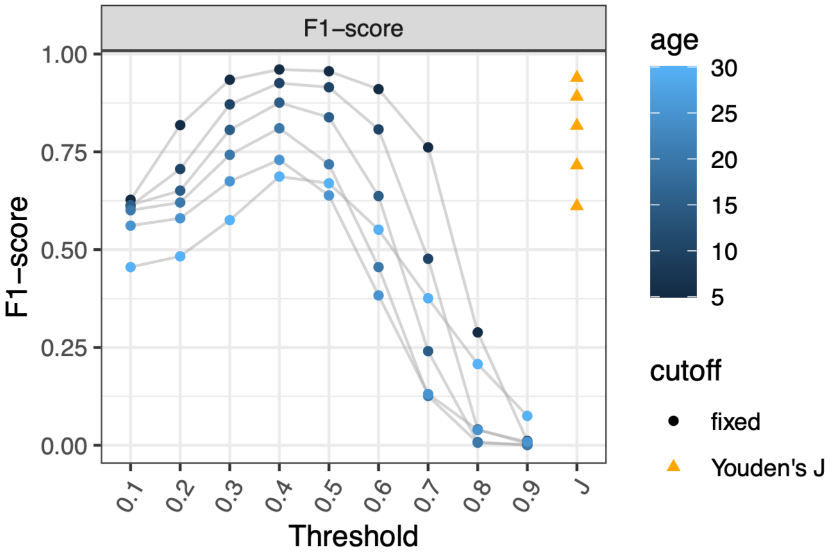

**Supplementary Figure 15: Results of the empirical applications are qualitatively similar when using fixed thresholds of 0.3, 0.5, or 0.6 instead of the Youden's J threshold.**

We repeated the ELEFANT analyses using fixed thresholds of 0.3, 0.5, and 0.6 to infer the presence of interactions, instead of the Youden's J threshold.

The different subplots represent the temporal variations of ancestral network properties. Squares indicate a significant departure from the null models (95% central range shown in orange), whereas the crosses indicate non-significance. Error bars represent the 95% central range of values obtained from the set of augmented ancestral networks.

**a) Mycorrhizal networks with an interaction threshold of 0.5**

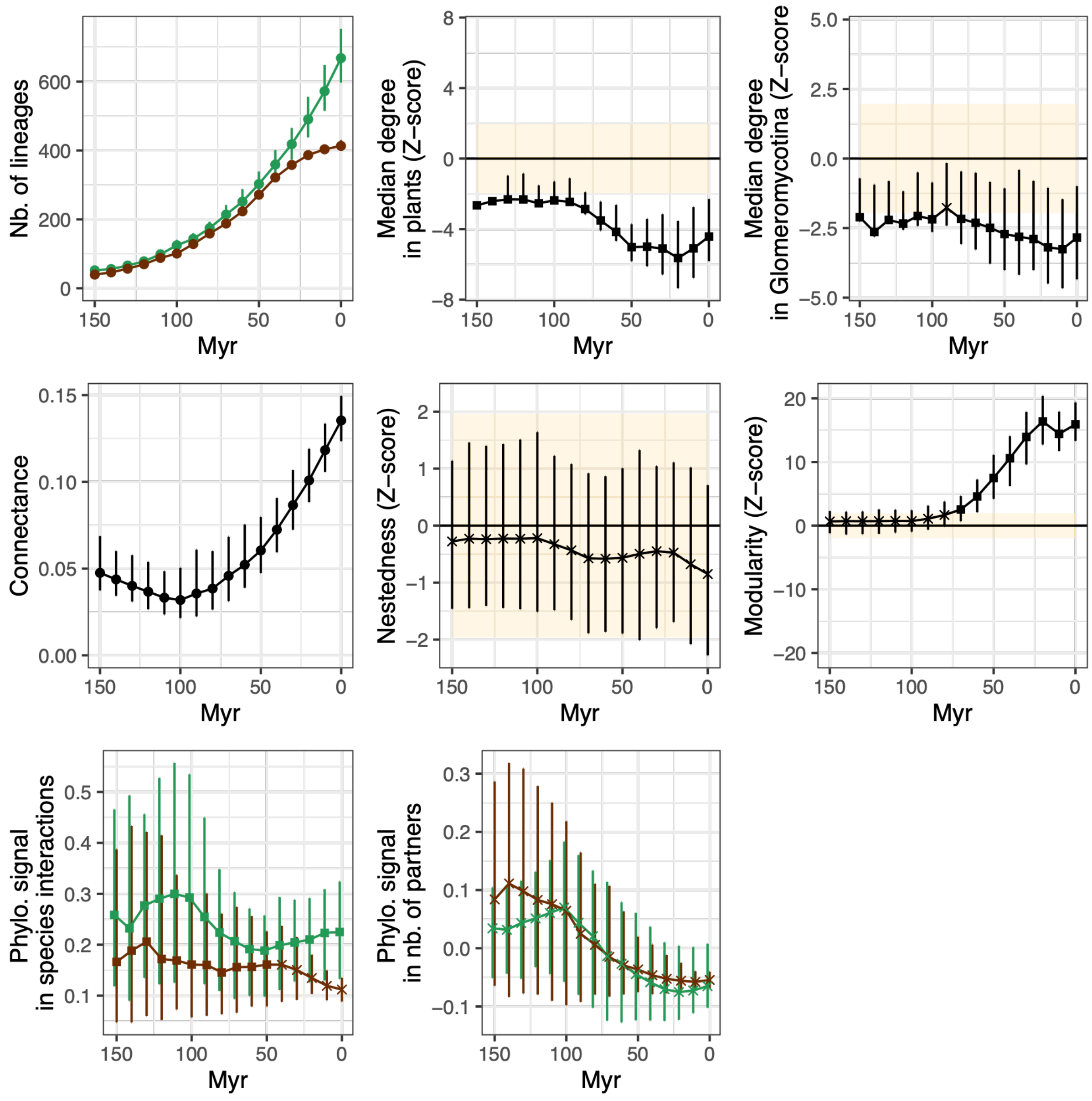

**b) Mycorrhizal networks with an interaction threshold of 0.3**

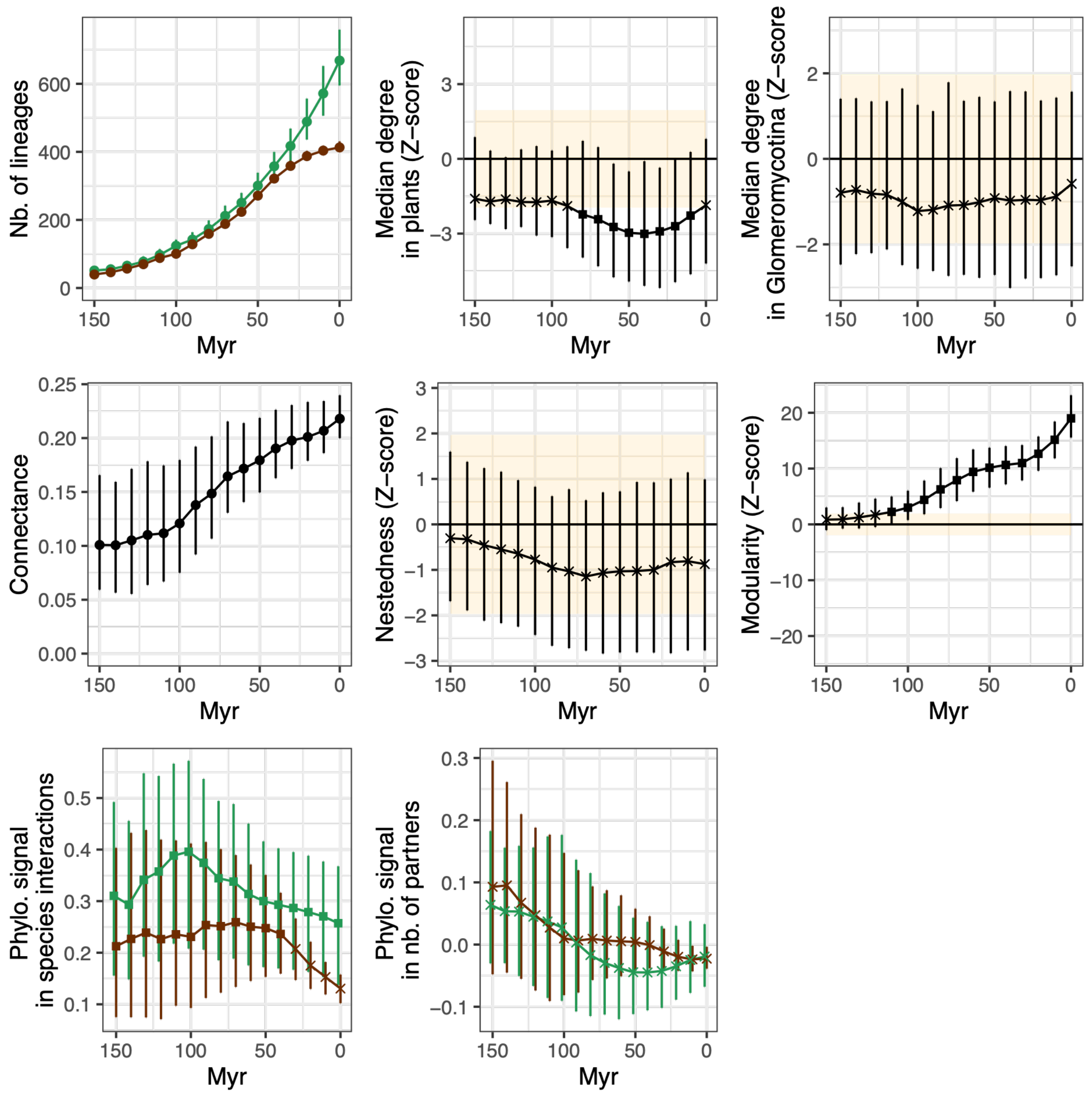

**c) Mycorrhizal networks with an interaction threshold of 0.6**

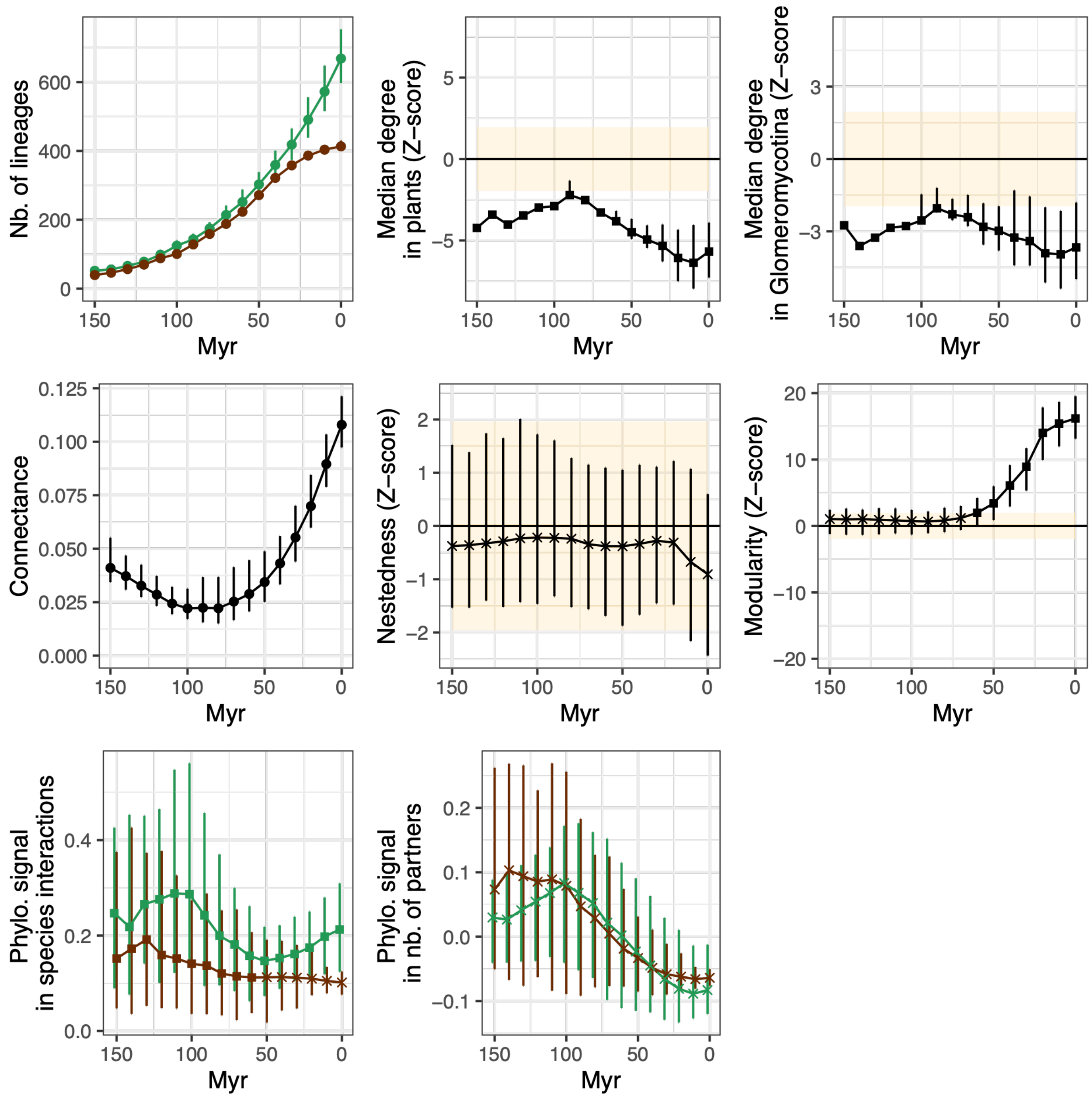

**d) Bat pollination networks with an interaction threshold of 0.5**

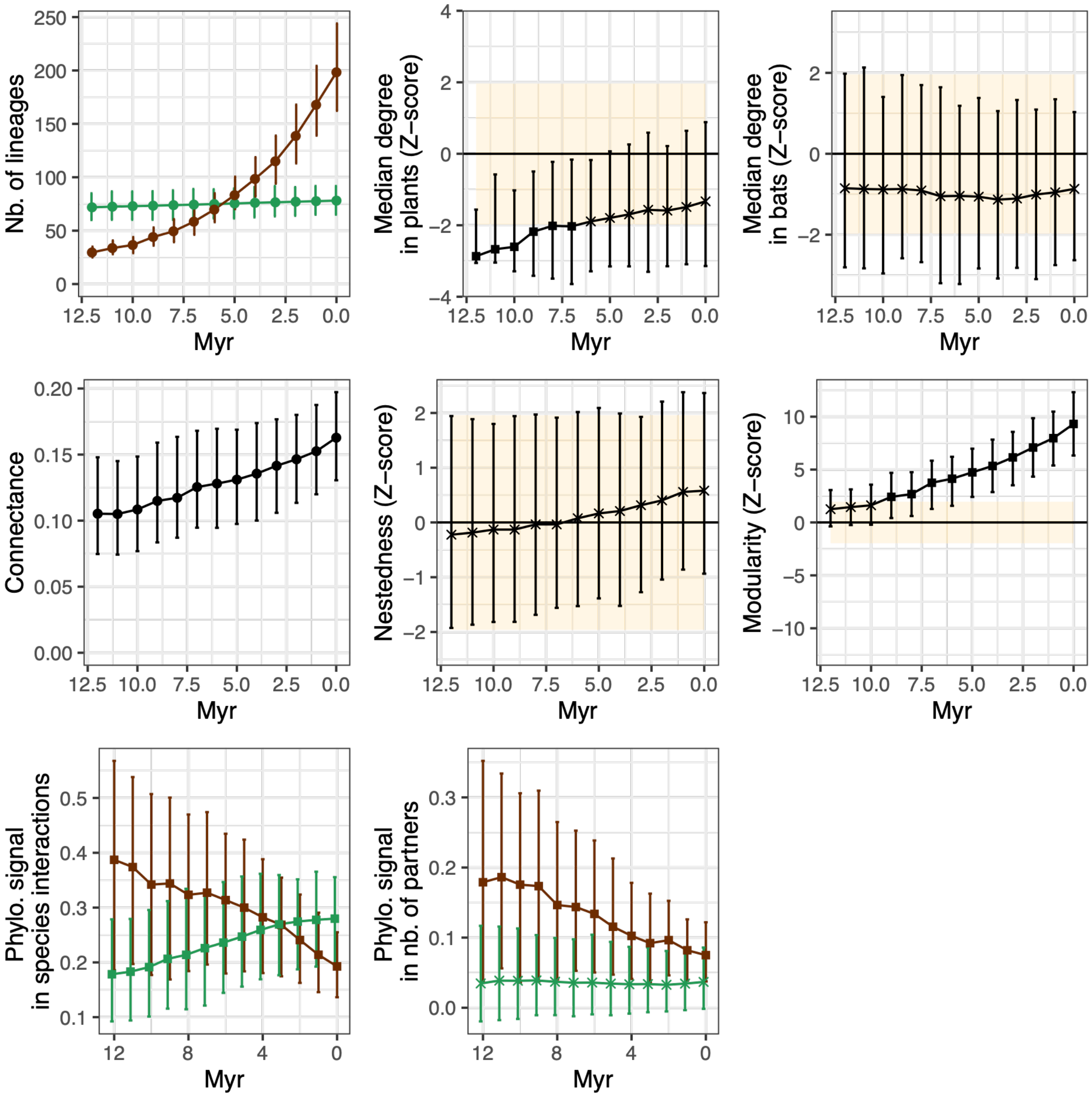

**e) Bat pollination networks with an interaction threshold of 0.3**

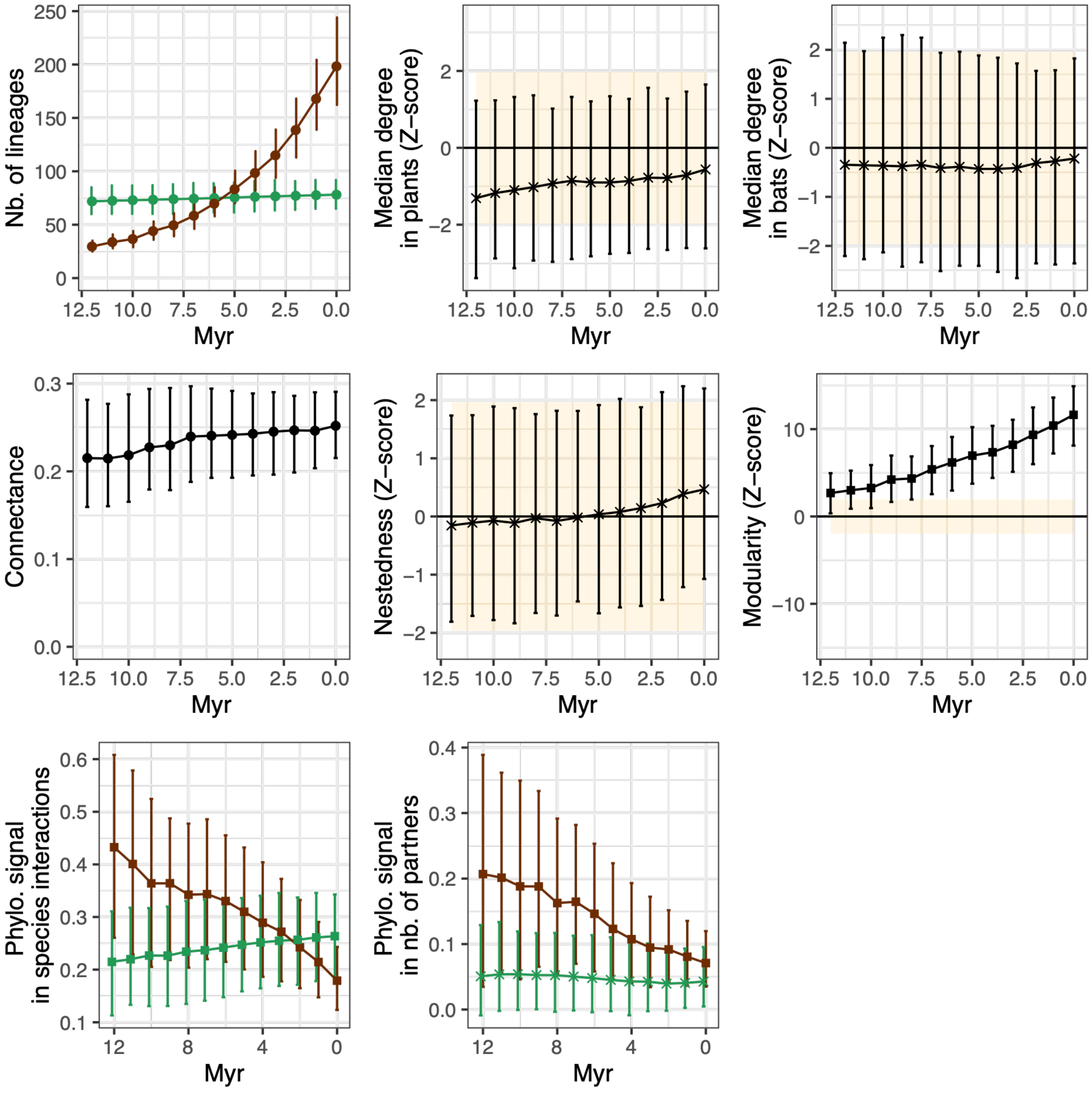

**f) Bat pollination networks with an interaction threshold of 0.6**

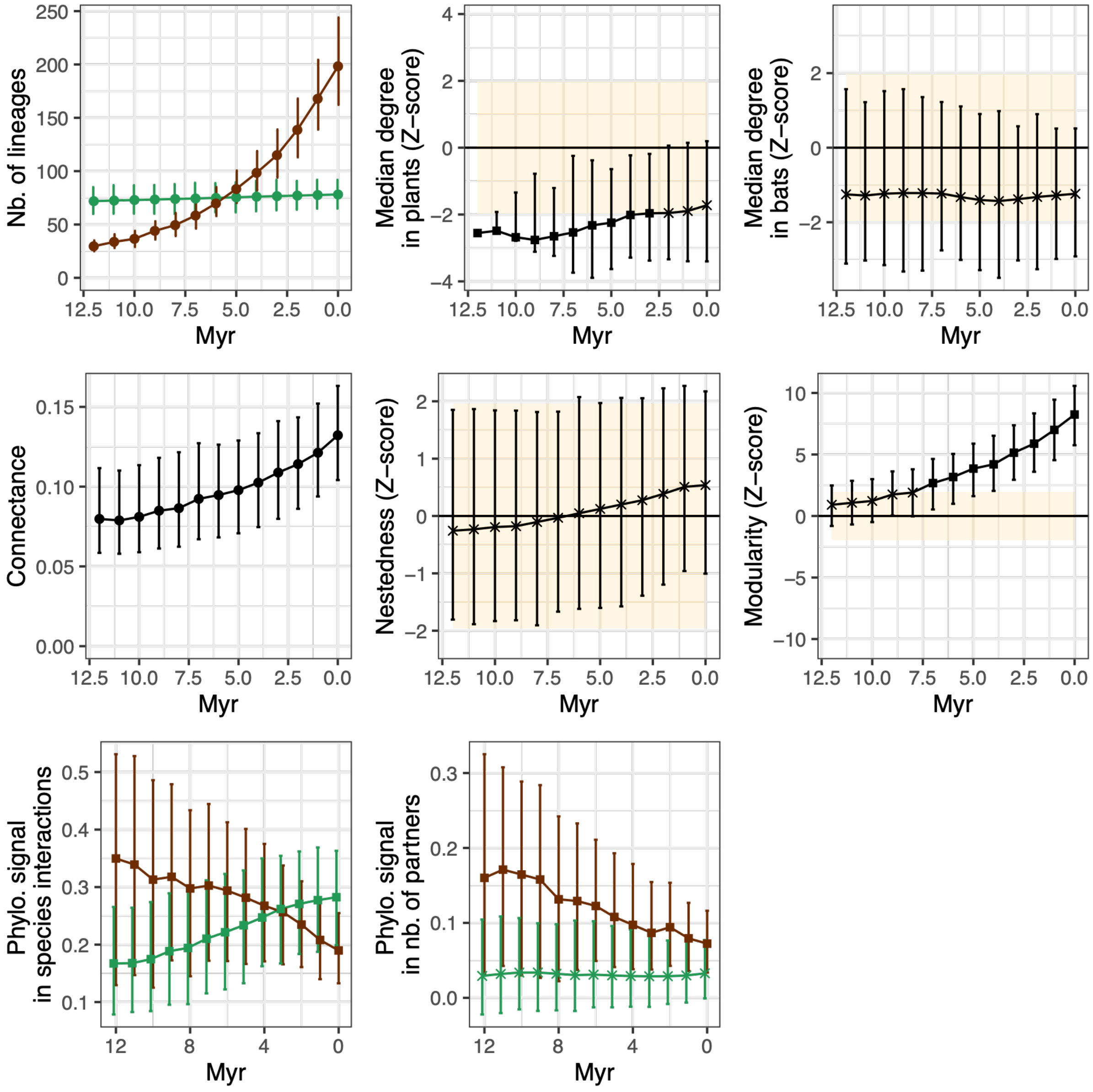

**g) Plant-bird seed dispersal networks in Europe with a threshold of 0.5**

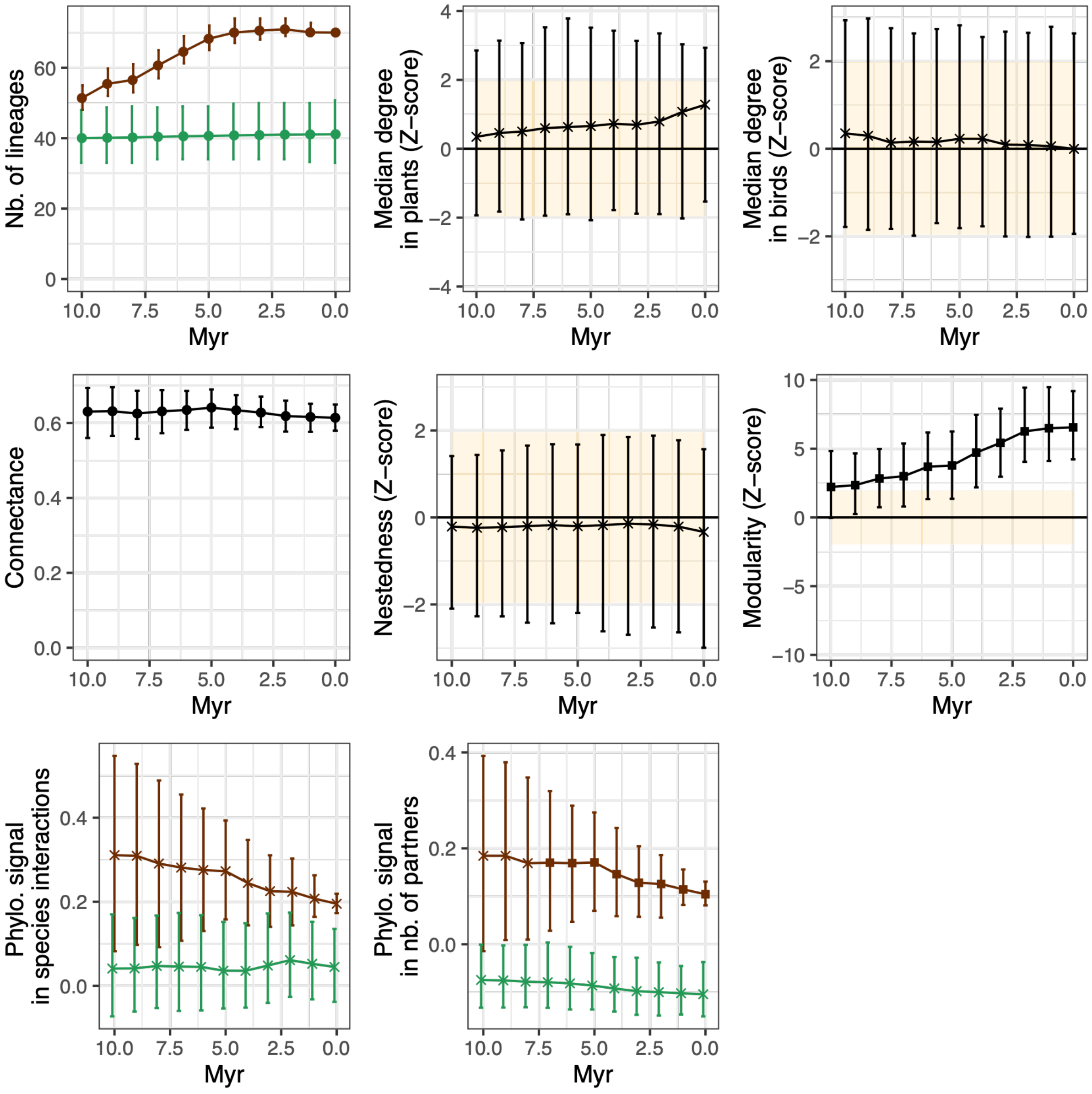

**h) Plant-bird seed dispersal networks in Europe with a threshold of 0.3**

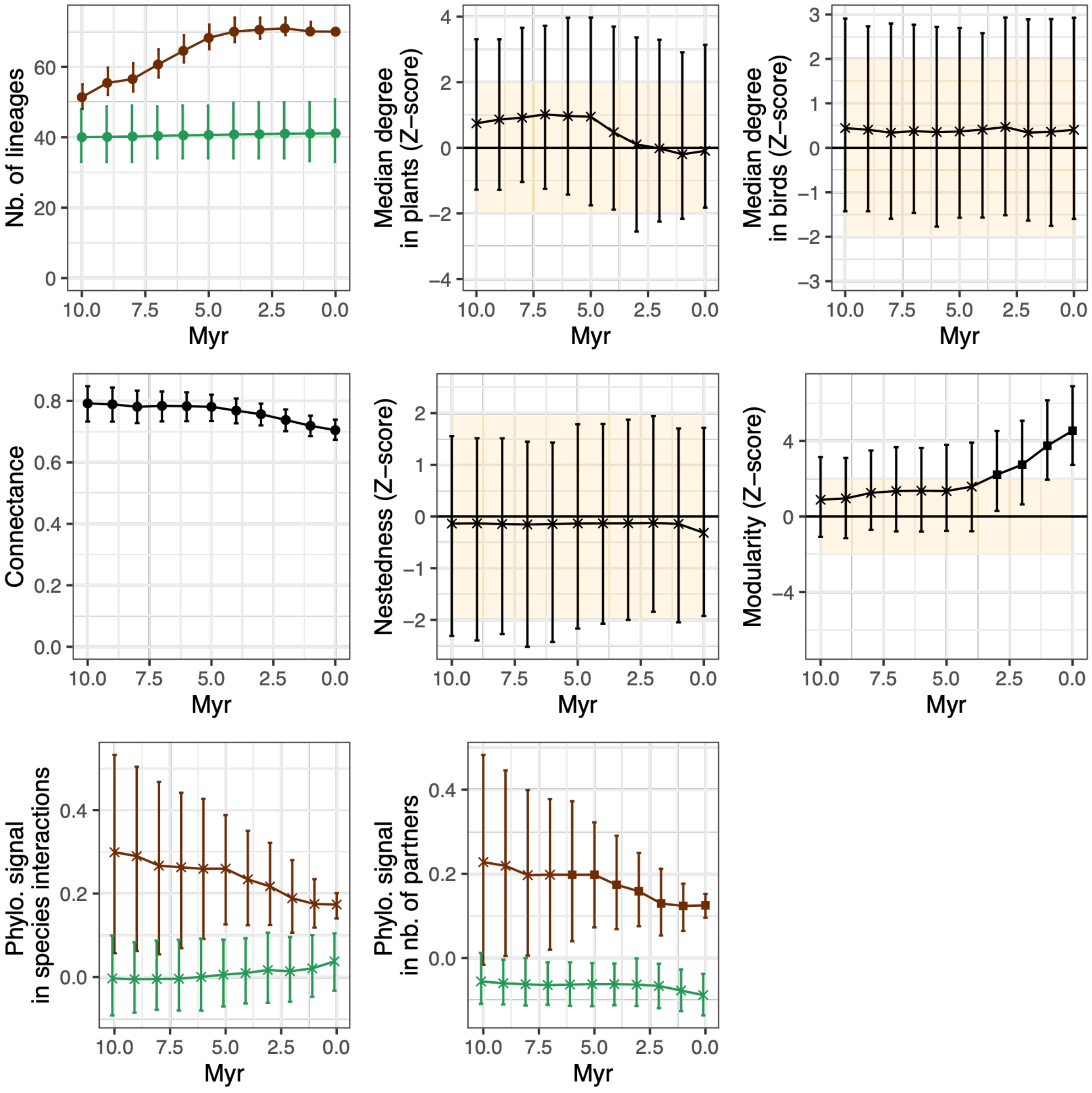

**i) Plant-bird seed dispersal networks in Europe with a threshold of 0.6**

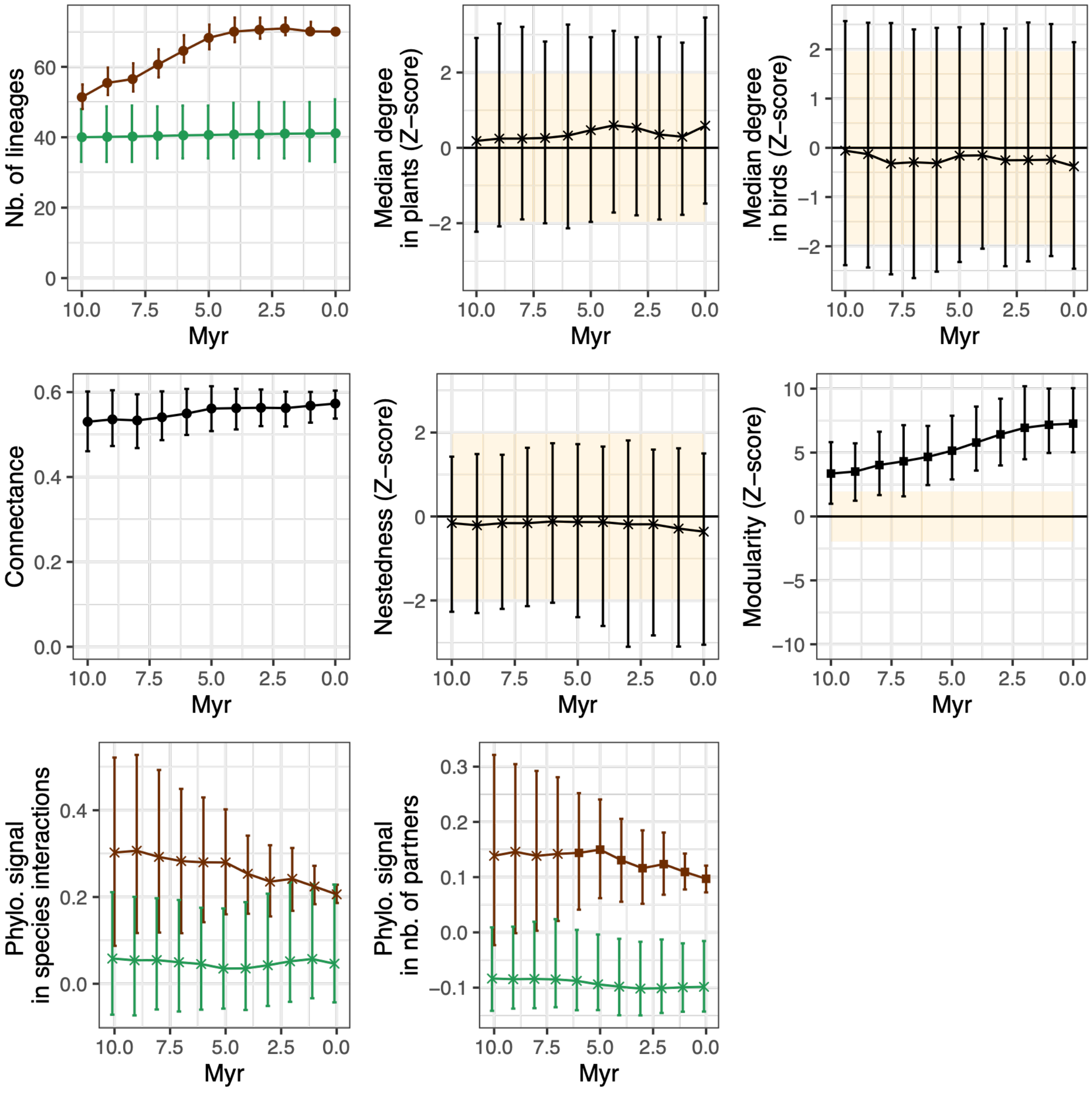

**j) Plant-bird seed dispersal networks in the Americas with a threshold of 0.5**

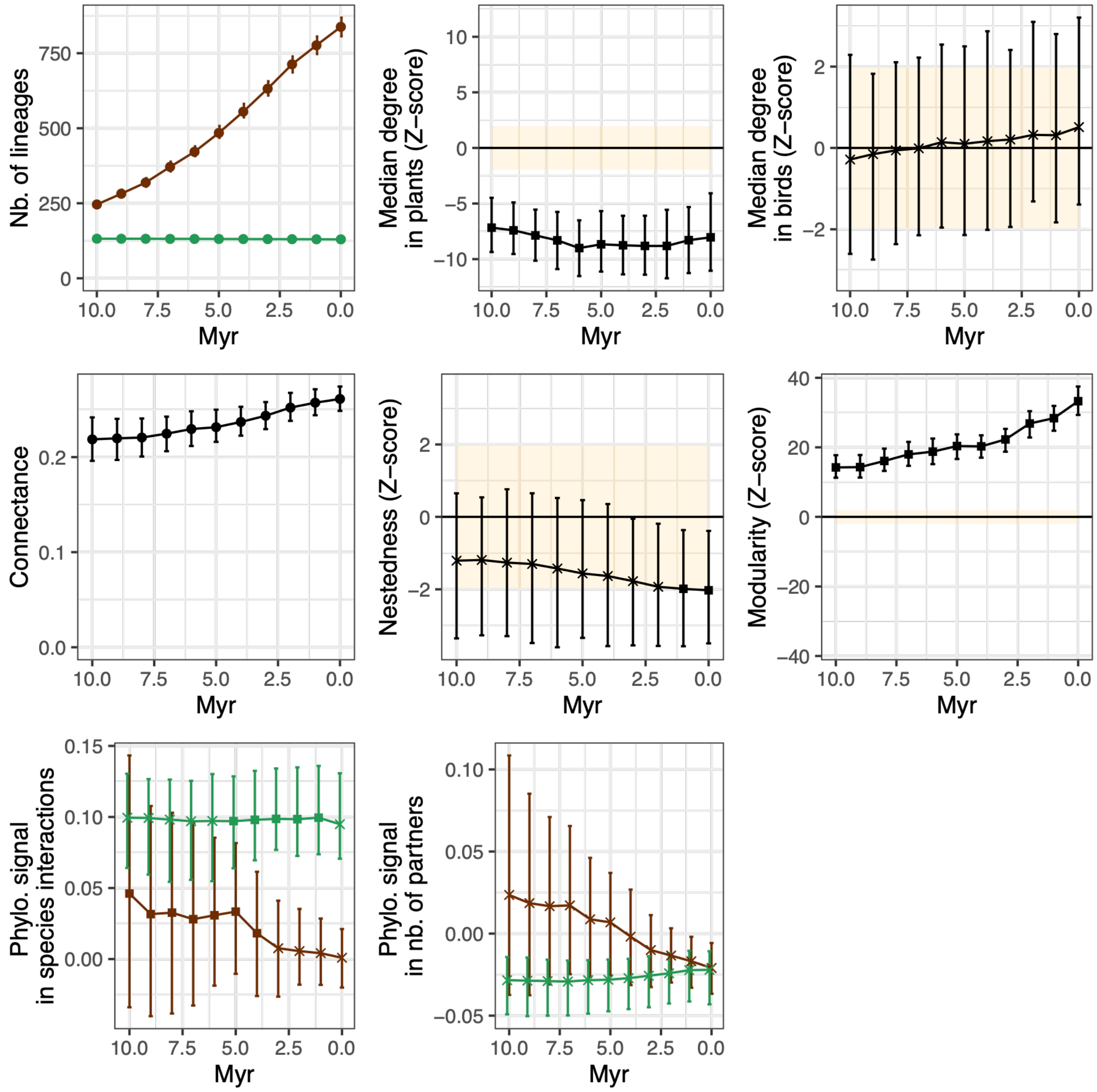

**k) Plant-bird seed dispersal networks in the Americas with a threshold of 0.3**

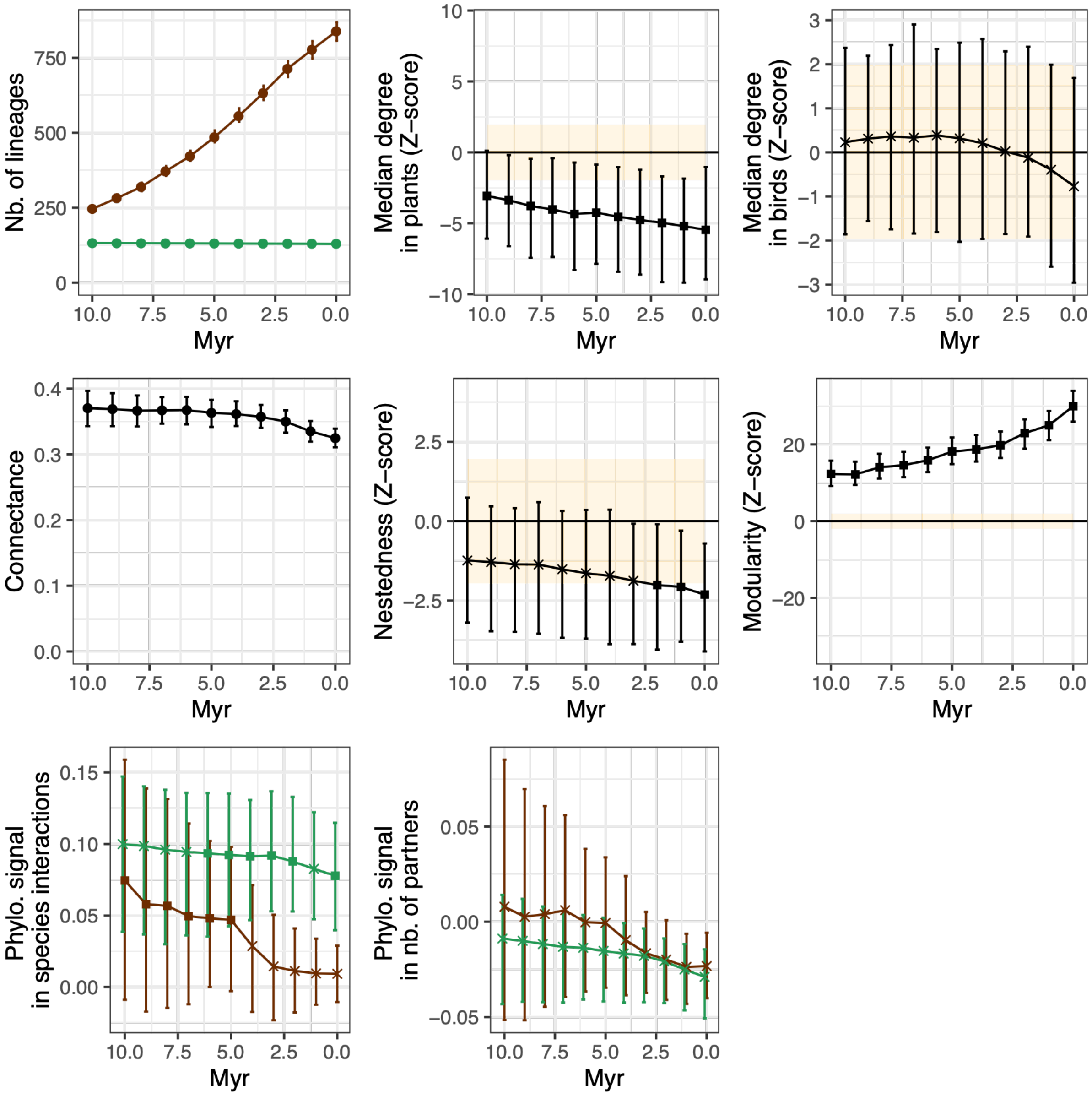

**l) Plant-bird seed dispersal networks in the Americas with a threshold of 0.6**

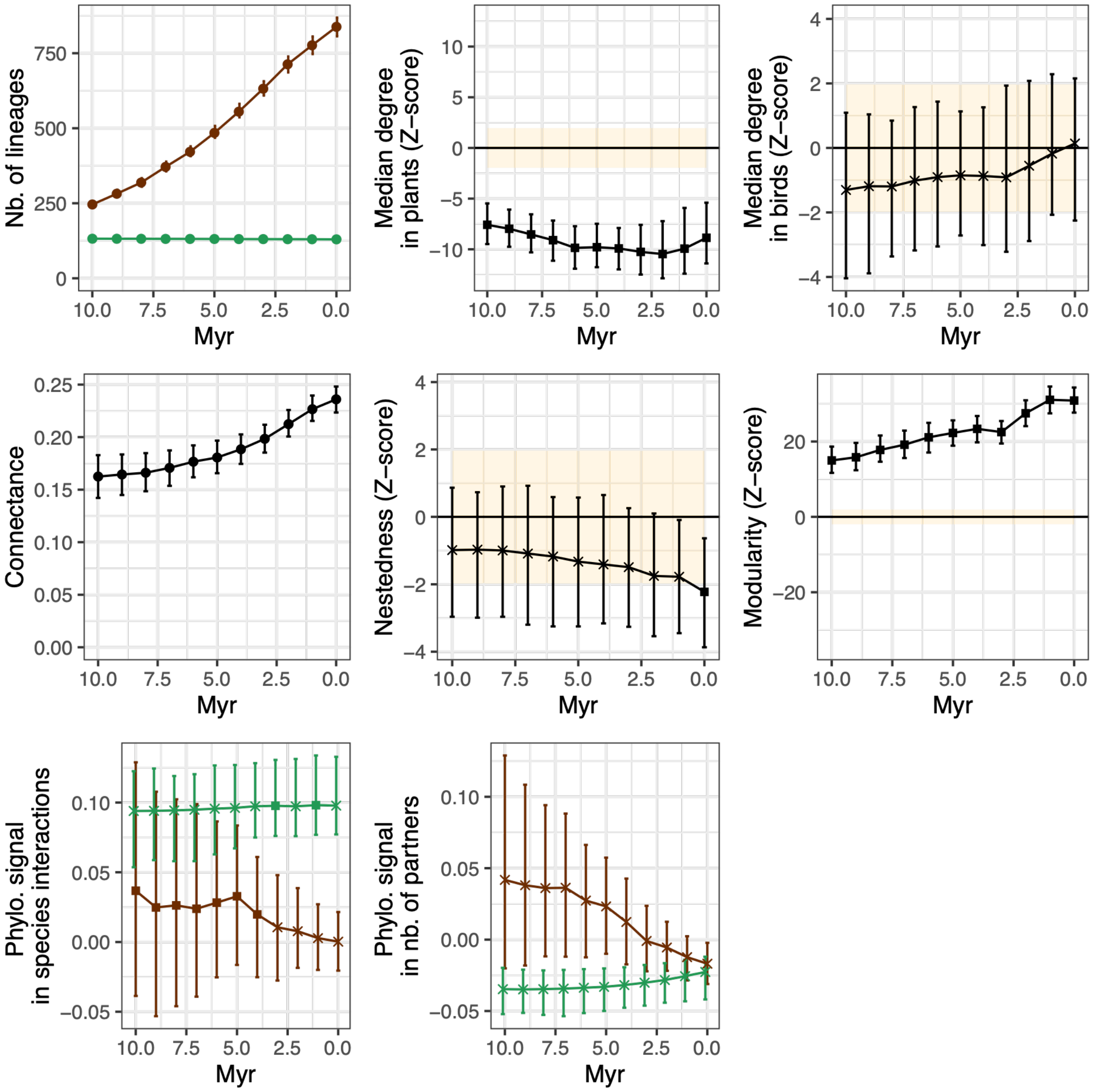

**Supplementary Figure 16: Examples of augmented trees with extinct and unsampled species resulting from our implemented step of data augmentation.**

Extinct or unsampled lineages are plotted in green. These lineages are absent from the reconstructed phylogenetic tree.

**a) Example with both extinct or unsampled lineages (λ=0.10, μ=0.05, and ρ=0.50)**

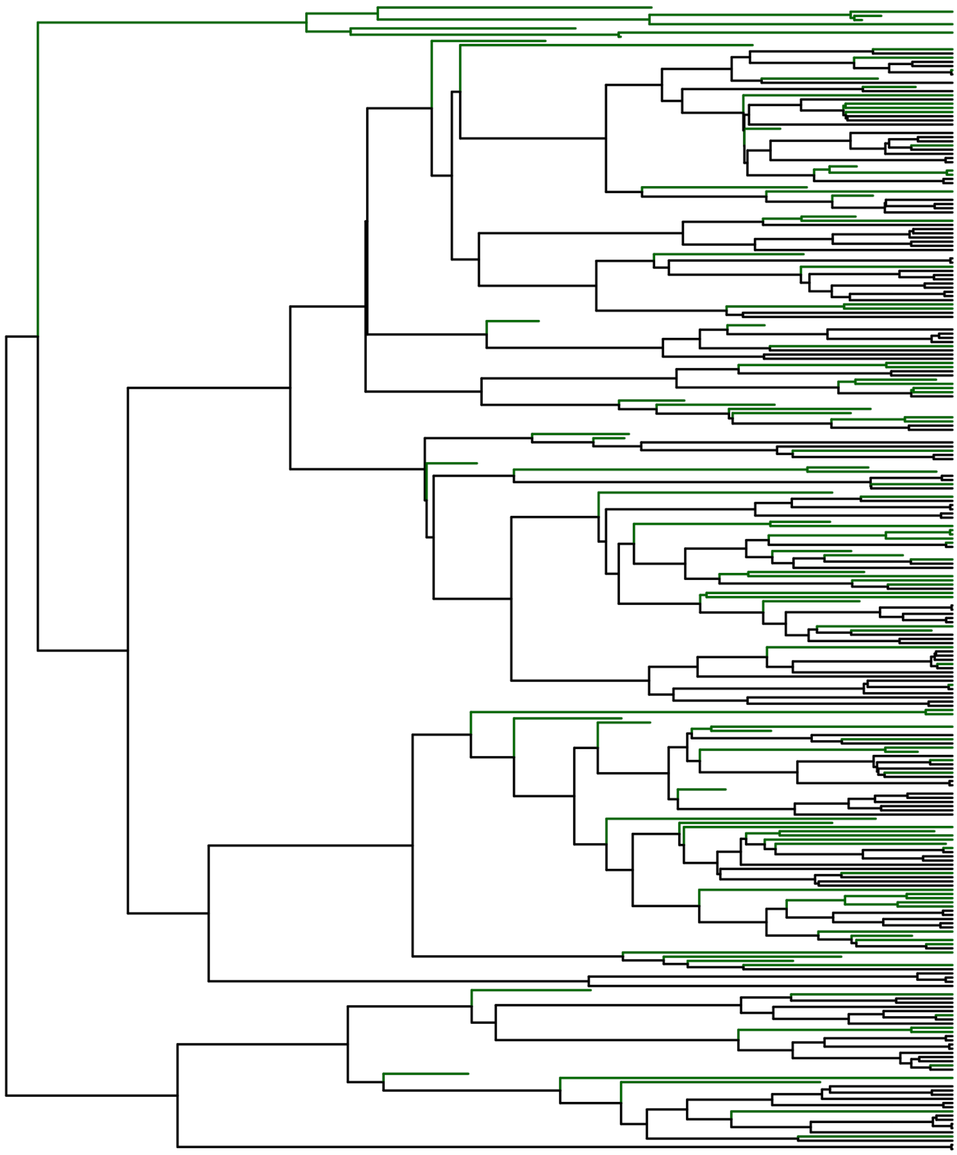

**b) Example with only extinct species (all present-day species are assumed to be sampled; λ=0.20, μ=0.15, and ρ=1)**

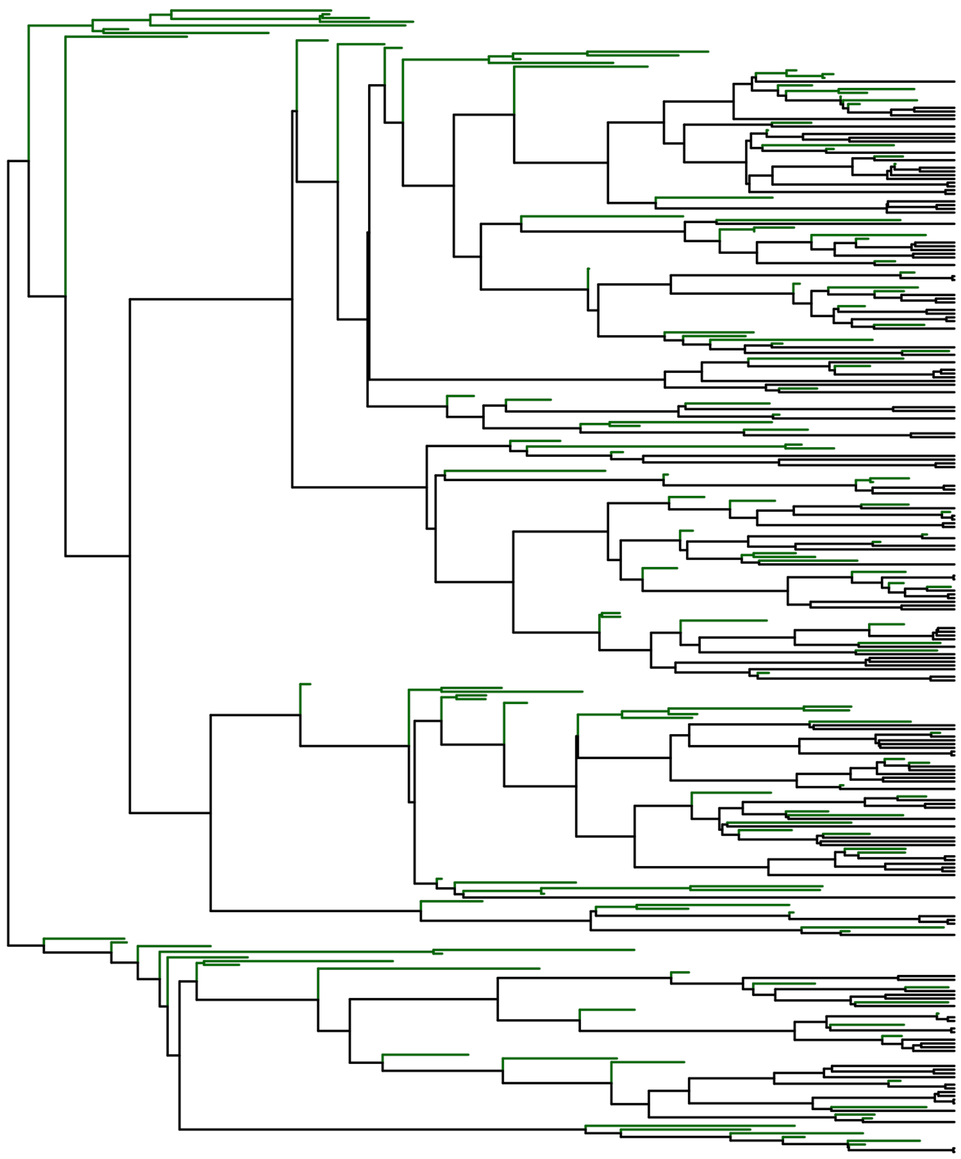

**Supplementary Tables:**

**Supplementary Table 1: Cross-validations confirm the ability of ELEFANT to recover subsampled interactions**

Based on the cross-validation results, this table reports the true positive rate (i.e., the proportion of true interactions correctly recovered by ELEFANT), the false positive rate (i.e., the proportion of inferred interactions that are actually false positives), and the cross-validation score (F1-score at the present time).

| Network | True positive rate | False positive rate | Cross-validation score (F1-score at present) |
|---|---|---|---|
| Arbuscular mycorrhizal symbiosis at global scale | 0.50 | 0.76 | 0.32 |
| Bat pollination in the Neotropics | 0.51 | 0.82 | 0.27 |
| Bird seed dispersal in Europe | 0.68 | 0.29 | 0.70 |
| Bird seed dispersal in the Americas | 0.55 | 0.60 | 0.46 |

**Supplementary Table 2: Angiosperms preferentially interact with Glomeraceae since their emergence, whereas early-diverging plant lineages interact with early-diverging Glomeromycotina.**

The table indicates the ratio of interactions observed between Angiosperms and Glomeraceae or between non-Angiosperms and non-Glomeraceae. For comparison, the theoretical ratio is computed given the number of species in each category and assuming random interactions.

| Age (in Myr) | Observed ratio | Theoretical ratio |
|---|---|---|
| 0 (present) | 0.75 | 0.62 |
| 10 | 0.76 | 0.62 |
| 20 | 0.76 | 0.62 |
| 30 | 0.77 | 0.62 |
| 40 | 0.77 | 0.63 |
| 50 | 0.76 | 0.61 |
| 60 | 0.73 | 0.59 |
| 70 | 0.72 | 0.58 |
| 80 | 0.69 | 0.55 |
| 90 | 0.66 | 0.54 |
| 100 | 0.65 | 0.53 |
| 110 | 0.59 | 0.5 |
| 120 | 0.55 | 0.49 |
| 130 | 0.54 | 0.5 |
| 140 | 0.55 | 0.52 |
| 150 | 0.61 | 0.57 |
| 160 | 0.66 | 0.62 |
| 170 | 0.67 | 0.62 |

**Supplementary Table 3: Effect of the biogeography on the composition of network modules:**

| Network | Metric | Mean | s.d. |
|---|---|---|---|
| Arbuscular mycorrhizal symbiosis at global scale | Number of bioregions per module for the plants | 4,00 | 1,58 |
| | Number of bioregions per module for the Glomeromycotina | 4,40 | 1,52 |
| | % of pairs of species from the same bioregion per module for the plants | 39% | 21% |
| | % of pairs of species from the same bioregion per module for the Glomeromycotina | 44% | 25% |
| Bat pollination in the Neotropics | Number of bioregions per module for the plants | 2,20 | 0,45 |
| | Number of bioregions per module for the bats | 2,80 | 0,45 |
| | % of pairs of species from the same bioregion per module for the plants | 50% | 19% |
| | % of pairs of species from the same bioregion per module for the bats | 45% | 13% |
| Bird seed dispersal in the Americas | Number of bioregions per module for the plants | 3,40 | 1,52 |
| | Number of bioregions per module for the birds | 3,20 | 1,30 |
| | % of pairs of species from the same bioregion per module for the plants | 54% | 28% |
| | % of pairs of species from the same bioregion per module for the birds | 45% | 22% |
| Bird seed dispersal in Europe | Number of bioregions per module for the plants | 2,25 | 0,50 |
| | Number of bioregions per module for the birds | 4,50 | 0,58 |
| | % of pairs of species from the same bioregion per module for the plants | 47% | 21% |
| | % of pairs of species from the same bioregion per module for the birds | 24% | 12% |

**Supplementary Table 4: Diversification rates for the empirical applications of ELEFANT**

The rates used in the main analyses were based on the most likely values reported in the literature or based on our own inferences, while the "other rates" scenarios explored other conditions accounting for the uncertainty in diversification rate estimates in the literature.

| System | Type | Speciation rate (λ) | Extinction rate (μ) | Sampling fraction (ρ) |
|---|---|---|---|---|
| Mycorrhizal network | Main analyses | Plants: 0.02<br>Fungi: 0.012 | Plants: 0.003<br>Fungi: 0.001 | Plants: 0.19<br>Fungi: 0.86 |
| Mycorrhizal network | No data augmentation | X | X | X |
| Mycorrhizal network | Other rates | Plants: 0.02<br>Fungi: 0.012 | Plants: 0.005<br>Fungi: 0.002 | Plants: 0.18<br>Fungi: 0.70 |
| Bat pollination network | Main analyses | Plants: 0.02<br>Bats: 0.20 | Plants: 0.003<br>Bats: 0.01 | Plants: 0.63<br>Bats: 0.31 |
| Bat pollination network | No data augmentation | X | X | X |
| Bat pollination network | Other rates | Plants: 0.02<br>Bats: 0.20 | Plants: 0.005<br>Bats: 0.02 | Plants: 0.84<br>Bats: 0.47 |
| Plant-bird seed dispersal network in Europe | Main analyses | Plants: 0.02<br>Birds: 0.15 | Plants: 0.003<br>Birds: 0.01 | Plants: 0.78<br>Birds: 1 |
| Plant-bird seed dispersal network in Europe | No data augmentation | X | X | X |
| Plant-bird seed dispersal network in Europe | Other rates | Plants: 0.02<br>Birds: 0.15 | Plants: 0.005<br>Birds:0.02 | Plants: 0.70<br>Birds: 0.80 |
| Plant-bird seed dispersal network in the Americas | Main analyses | Plants: 0.02<br>Birds: 0.15 | Plants: 0.003<br>Birds: 0.01 | Plants: 0.93<br>Birds: 0.73 |
| Plant-bird seed dispersal network in the Americas | No data augmentation | X | X | X |
| Plant-bird seed dispersal network in the Americas | Other rates | Plants: 0.02<br>Birds: 0.15 | Plants: 0.005<br>Birds: 0.02 | Plants: 0.80<br>Birds: 0.60 |
